# A systematic review and meta-analysis of the potential non-human animal reservoirs and arthropod vectors of the Mayaro virus

**DOI:** 10.1101/2021.07.28.454243

**Authors:** Michael Celone, Bernard Okech, Barbara A. Han, Brett M. Forshey, Assaf Anyamba, James Dunford, George Rutherford, Neida K. Mita Mendoza, Elizabet Lilia Estallo, Ricardo Khouri, Isadora Cristina de Siqueira, Simon Pollett

## Abstract

Improving our understanding of Mayaro virus (MAYV) ecology is critical to guide surveillance and risk assessment. We conducted a PRISMA-adherent systematic review of the published and grey literature to identify potential arthropod vectors and non-human animal reservoirs of MAYV. We searched PubMed, Embase, Web of Science, SciELO and grey-literature sources including PAHO databases and dissertation repositories. Studies were included if they assessed MAYV virological/immunological measured occurrence in field-caught, domestic, or sentinel animals or in field-caught arthropods. We conducted an animal seroprevalence meta-analysis using a random effects model. We compiled granular georeferenced maps of non-human MAYV occurrence and graded the quality of the studies using a customized framework. Overall, 57 studies were eligible out of 1523 screened, published between the years 1961 and 2020. Seventeen studies reported MAYV positivity in wild mammals, birds, or reptiles and five studies reported MAYV positivity in domestic animals. MAYV positivity was reported in 12 orders of wild-caught vertebrates, most frequently in the orders Charadriiformes and Primate. Sixteen studies detected MAYV in wild-caught mosquito genera including *Haemagogus, Aedes, Culex, Psorophora, Coquillettidia,* and *Sabethes*. Vertebrate animals or arthropods with MAYV were detected in Brazil, Panama, Peru, French Guiana, Colombia, Trinidad, Venezuela, Argentina, and Paraguay. Among non-human vertebrates, the Primate order had the highest pooled prevalence (PP) at 13.1% (95% CI: 4.3-25.1%). From the three most studied primate genera we found the highest prevalence was in *Alouatta* (PP: 32.2%, 95% CI: 0.0-79.2%), followed by *Callithrix* (PP: 17.8%, 95% CI: 8.6-28.5%), and *Cebus/Sapajus* (PP: 3.7%, 95% CI: 0.0-11.1%). We further found that MAYV occurs in a wide range of vectors beyond *Haemagogus* spp. The quality of evidence behind these findings was variable and prompts calls for standardization of reporting of arbovirus occurrence. These findings support further risk emergence prediction, guide field surveillance efforts, and prompt further *in-vivo* studies to better define the ecological drivers of MAYV maintenance and potential for emergence.

**Author Summary:** Mayaro virus (MAYV) is an emerging tropical public health threat in the Americas. We conducted a georeferenced, quality-graded systematic review to evaluate the current evidence regarding MAYV occurrence in non-human vertebrates and arthropods. Overall, 57 studies were eligible out of 1523 screened, published between the years 1961 and 2020. Seventeen studies reported MAYV positivity in wild mammals, birds, or reptiles and five studies reported MAYV positivity in domestic animals. MAYV positivity was reported in 12 orders of wild-caught vertebrates, most frequently in the orders Charadriiformes and Primate. Our systematic review identified 12 orders of wild-caught vertebrates and seven mosquito genera with evidence of MAYV occurrence. Primates had the highest pooled MAYV prevalence according to a seroprevalence meta-analysis. The graded quality of evidence behind these findings was variable and prompts calls for standardization of reporting of MAYV and perhaps other emerging arbovirus occurrence in animals and vectors. This study provides important information for public health authorities and disease ecologists concerned with the growing threat of MAYV in Latin America. Our analysis provides a foundation for future laboratory and field studies focused on the MAYV transmission cycle.

## Introduction

First detected in Trinidad in 1954 [1], Mayaro virus (MAYV) is a zoonotic *Alphavirus* that is endemic in several Latin American countries. Like Chikungunya virus (CHIKV), MAYV may cause complications such as debilitating arthralgia but often presents with a non-specific constellation of symptoms and signs that may be clinically indistinguishable from other vector borne diseases such as dengue or Zika [2]. There is no current licensed vaccine or antiviral treatment for MAYV infections, and the current standard of clinical treatment is supportive care only [2, 3].

MAYV has caused periodic outbreaks in humans in Brazil [4, 5], Bolivia [6], and Venezuela [7], while surveillance studies and serological surveys have detected MAYV in humans in several countries throughout the Americas including Peru [8], Suriname [9], Mexico [10], Colombia [11], French Guiana [12], and Haiti [13]. These findings demonstrate widespread circulation of the virus throughout the region. A recent 2019 epidemiological alert by the Pan American Health Association (PAHO) has emphasized the need for increased awareness of and extended surveillance for this emerging virus in the Americas [3]. However, the precise areas of risk from MAYV throughout the Americas remain unclear. Understanding the ecology and distribution of MAYV remains a major obstacle in predicting areas that are at high risk of transmission to humans and domestic animals.

Current evidence suggests that MAYV is maintained in nature through a sylvatic transmission cycle involving mosquito vectors and non-human animal reservoirs. Therefore, human MAYV cases reported to date likely represent direct sylvatic spillovers. Residing near forested areas [12] and hunting in the rainforest [14] have been identified as risk factors for MAYV infection in humans, highlighting the importance of the sylvatic transmission cycle and the potential for spillover events.

Identification of the non-human vertebrate animals (i.e., reservoirs) involved in MAYV transmission is an important step in delineating the human populations at greatest risk. The spillover of MAYV into humans represents a complex interaction of processes involving the density and distribution of reservoirs and vectors, as well as the prevalence and intensity of infection among reservoirs [15].

Identifying the non-human to, the challenges associated with establishing evidence of infection in wild animal populavertebrates that may serve as MAYV reservoirs is a difficult task due to a myriad of issues including, but not limited tions [16, 17]. High seroprevalence of a pathogen in an animal population does not necessarily implicate a given host as an efficient reservoir; conversely, low seroprevalence at a single point in time cannot definitively rule out an animal as a reservoir [17]. Due to the relatively short viremia of MAYV (approximately 3-10 days) molecular assays may be unsuccessful in detecting virus [18], necessitating the use of serological assays such as hemagglutination-inhibition (HI) assays, enzyme-linked immunosorbent assays (ELISA), or plaque-reduction neutralization tests (NT).

Several studies have been conducted to clarify the precise vertebrate hosts that may serve as MAYV reservoirs. High seroprevalence among non-human primates (NHPs) in Brazil [19], French Guiana [12], and Panama [20] provides evidence that NHPs may play an important role in the MAYV transmission cycle. MAYV antibodies have also been detected in mammals including rodents and marsupials [21] as well as several avian species [19]. Unfortunately, there is significant heterogeneity in the study methods used to identify potential MAYV reservoirs and there remains a high level of uncertainty surrounding the role of various non-human vertebrate species in the MAYV transmission cycle.

Studies have also been conducted in wild-caught mosquito populations as well as in controlled laboratory conditions in order to identify potential arthropod vectors of MAYV. One study in Brazil [19] suggested that the canopy-dwelling *Haemagogus janthinomys* mosquito is an important vector of MAYV. Additional mosquito species including *Aedes aegypti, Ae. albopictus*, and several anopheline species have been shown to be competent vectors in laboratory settings [22-24], posing a potential but as yet theoretical risk of urban MAYV cycles. The occurrence of MAYV in the city of Manaus has also led to concerns about the involvement of *Aedes* mosquitoes in a MAYV urban transmission cycle [25].

Although many non-human vertebrate animals and arthropod species have been proposed as capable MAYV reservoirs or vectors, our understanding of the MAYV transmission cycle and ecology remains limited. Collating and evaluating the current evidence regarding the potential MAYV reservoirs and vectors are important steps in characterizing MAYV transmission ecology and identifying the communities at greatest risk for MAYV outbreaks. Therefore, the goal of this systematic review is to evaluate the current evidence regarding MAYV occurrence in non-human vertebrates and arthropods. We present here the first structured evaluation of the potential vector and non-human reservoir range of MAYV, including the development of custom criteria for grading the quality of evidence of arbovirus occurrence in invertebrate and vertebrate non-human hosts.

## Methods

This systematic review and meta-analysis were conducted according to the PRISMA 2020 Checklist [26] (see **S1 Table**). A protocol was developed but was not uploaded to PROSPERO.

### Information Sources

We conducted a systematic review of original research articles, reports, and dissertations that attempted to identify potential non-human animal reservoirs or arthropod vectors of MAYV. We first searched Embase, Web of Science, PubMed, and SciELO databases for English, Spanish, and Portuguese language articles published between 1954 (the year MAYV was first isolated) and March 21, 2020. We searched all databases using the highly sensitive search term “Mayaro”. A PubMed alert using the search term “Mayaro” was also set to capture any additional studies that were published between the initial search and May 2021. This database search was extended using bioRxiv (https://www.biorxiv.org/) and medRxiv (https://www.medrxiv.org/) pre-print databases. We complemented these database search results with ‘grey literature,’ including hand-searched bibliographies of MAYV review articles (including systematic reviews), dissertations from several Brazilian university repositories, the Pan American Health Organization (PAHO) Institutional Repository for Information Sharing database (<u>iris.paho.org</u>), the GIDEON database (https://www.gideononline.com/), and GenBank [27] (https://www.ncbi.nlm.nih.gov/genbank/). In addition, we searched conference handbooks that are available online (2004-2019) from the American Society of Tropical Medicine and Hygiene (https://www.astmh.org/annual-meeting/past-meetings).

### Eligibility Criteria

We included studies that evaluated past or current MAYV infection in non-human vertebrates using methods including virus isolation, molecular detection, and serosurveys. We also included studies that screened arthropods for MAYV using virus isolation and molecular detection. Original research studies were considered for eligibility if they assessed MAYV positivity in field-caught, captive, or sentinel non-human vertebrates or field-caught arthropods. Studies that met any of the following exclusion criteria were not included: studies involving only humans; studies not reporting original data (e.g., review articles, perspective pieces, editorials, recommendations, and guidelines); duplicate studies; *in vitro* studies such as vector cell-line or mammal cell line experiments; laboratory-based vector competence studies that did not explicitly demonstrate the detection of MAYV in a wild-caught vector; *in-vivo* lab-reared animal studies or any laboratory-based study that experimentally inoculated an animal to test theoretical reservoir status.

### Selection process

All articles were organized using EndNote software version X9 (Clarivate, Philadelphia, Pennsylvania, USA), and data were abstracted into a Microsoft Excel table. Two reviewers independently screened all titles and abstracts to determine articles that could immediately be excluded and articles that should be included in the second stage of review. Results were compared to reconcile any differences between the two reviewers. The first and second reviewers then independently read the full text of potentially eligible articles identified through screening and selected the articles that were candidates for inclusion in the study. Results were compared to reconcile any differences between the two reviewers. A third-party reviewer adjudicated when consensus was not reached between the two reviewers during the first or second stage review. From those studies deemed eligible, data were extracted from articles by one reviewer using the data abstraction tool in Microsoft Excel.

### Data abstraction

Relevant information was abstracted by one reviewer in an Excel sheet. Information for each article was abstracted across several domains including publication details (author and affiliation, study title, study funding), study methods (date and location of study, study design, laboratory methods to assess MAYV positivity), and study results (sample size, taxonomic classification, proportion of animals testing positive for MAYV, location of vertebrates/arthropods testing positive for MAYV). A second reviewer randomly selected and reviewed five articles for review to validate the data abstraction process.

### Grading quality of evidence

We developed a customized grading system to assess the quality of each study included in our review. Several published studies have employed a similar grading system to assess evidence quality of included articles [28-30]. We assigned each study in our systematic review a grade for each of four quality items: clarity of research question/objective (*Was the research question/objective clearly described and stated?*); description of study methods (*Were the study methods presented in a reproducible way?*); description of sampling methods (*Was the sampling method described in detail?*); and validity of diagnostic tests (*Was MAYV positivity measured in a valid way?*). For each quality item, eligible studies were assigned a score of 3 (strong evidence), 2 (moderate evidence), 1 (weak evidence), or unable to judge. Studies were deemed unable to judge if the information provided was insufficient to assign quality scores (e.g., a single GenBank entry or conference abstract).

A score of 3 was assigned for the *description of sampling methods* item if authors thoroughly described the type of trap used, the habitats in which traps were set, how often traps were checked, and the results of trapping (i.e., were animals reported to the species level). For studies that assessed MAYV in vertebrate animals, a score of 3 was assigned for the *validity of diagnostic tests* item if MAYV positivity was assessed using RT-PCR, viral culture, or high-specificity serological method (i.e., plaque reduction NT); a score of 2 was assigned if MAYV positivity was assessed using non-specific serological assay (i.e., HI and ELISA); and a score of 1 was assigned if MAYV positivity was based on presumptive exposure only with no confirmatory assay. For studies that assessed MAYV in arthropods, a score of 3 was assigned for this item if MAYV positivity was assessed using viral culture; a score of 2 was assigned if MAYV positivity was assessed using RT-PCR or metagenomics; and a score of 1 was assigned if MAYV positivity was based on presumptive exposure only with no confirmatory assay. A score of “NA” was assigned for the *validity of diagnostic tests* item if studies did not detect MAYV positivity in any animal or arthropod samples.

Quality review scores were recorded in two different Excel documents for animal reservoir studies and arthropod vector studies, respectively. Two reviewers independently graded the evidence quality for each study and results were compared to reconcile any differences between the two reviewers. A third-party reviewer adjudicated if consensus was not reached between the two reviewers.

## Data analysis

### Descriptive Analysis

Descriptive statistics were presented by species for potential animal reservoirs showing the total sample size, proportion infected, and locations of infected animals. Descriptive statistics were presented by species for potential arthropod vectors showing the total sample size and total pools tested for virus (if applicable), the number of MAYV isolates or PCR-positive pools, and locations of infected arthropods. Maps were developed using ArcGIS software [31] to display the geographic distribution of MAYV-positive animals and vectors.

### Pooled Analysis

Due to the heterogeneity of study designs and outcome measurements, a quantitative meta-analysis across all eligible studies was not possible. Instead, we conducted a seroprevalence meta-analysis using the studies that reported MAYV seroprevalence (i.e., using serological methods including HI, ELISA, or NT) in non-human vertebrate animals. Pooled prevalence estimates were stratified by taxonomic order and an additional analysis was conducted among the various Primate genera. Orders were excluded from the analysis if the total sample size was less than 10 or if no MAYV-positive samples were reported within that order. Pooled seroprevalence was first calculated based on all available data, regardless of test method. This included the samples that tested MAYV-positive based on HI alone (when no confirmatory assay was performed) as well as the samples that were confirmed positive by an NT. Only monotypic reactions to MAYV were included in the meta-analysis in the absence of confirmatory NT. A sensitivity analysis was then conducted using only the MAYV-positive samples that were confirmed using NT. Positive samples that were based on HI alone (without confirmatory NT) were excluded from this analysis, although all MAYV-negative samples were retained. This sensitivity analysis was conducted to account for the low specificity of HI compared to NT [32] and provided a more conservative estimate of seroprevalence.

Due to the substantial differences across studies including sample size, study design, species sampling methods, and geographical location, a random effects model was used for analysis [33, 34]. The Freeman-Tukey double-arcsine transformation was implemented to calculate a proportion, based on the recommendation of Barendregt et al. [35]. A sensitivity analysis was conducted using a generalized linear mixed model (GLMM) with a logit transformation, due to the potential for misleading results with the double-arccosine transformation [36, 37]. Measures of variance ( τ^2^), heterogeneity (*I*^2^), and statistical significance are presented for each random effects model. An additional sensitivity analysis was conducted using a fixed effects model. Results of sensitivity analyses are presented in the Supplementary materials.

The *I^2^* statistic measures inconsistency across study results and is calculated as *I^2^* = 100% x (Q - df) / Q [38]. The *I^2^* statistic ranges between 0% and 100%, where a value of 0% represents no heterogeneity and larger values represent increased heterogeneity. Animal seroprevalence estimates with 95% confidence intervals (CIs) weighted by sample size are presented as forest plots. All analyses were conducted using the ‘*meta’* package in R statistical software version 4.0.2 (R Project for Statistical Computing, Vienna, Austria) [39, 40].

### Estimation of bias

An assessment of publication bias was carried out for meta-analyses that included five studies or more. Bias was assessed using funnel plots and tests for funnel plot asymmetry based on methods proposed by Egger [41]. If the Egger’s test revealed bias, the Trim and Fill technique was used to estimate the effect of missing studies on the outcomes of the meta-analysis [42].

### Georeferencing of MAYV occurrence

All available location information from each confirmed MAYV infection (animal and mosquito) was extracted from each article and georeferenced based on methods that have been described previously [43, 44]. Each occurrence of MAYV was designated as either a point or polygon location according to the spatial resolution provided in the study. When specific latitude and longitude coordinates were provided, they were verified in GoogleMaps and designated as a point location. If a neighborhood, town, village, or small city was explicitly mentioned in the article and fell within a 5×5 km grid cell, it was designated as a point location and its centroid coordinates were recorded. For studies that report a less precise spatial resolution such as states or counties, first level (ADM1) or second level (ADM2) administrative divisions were recorded as polygons. If the size of a specific named location was greater than a 5×5 km grid cell the occurrence was assigned to a custom polygon created in ArcGIS that encompassed the extent of that location. If place names were duplicated (i.e., the ADM1 and ADM2 units had the same name), the coarsest spatial resolution was used. Country shapefiles were accessed through the geoBoundaries Global Administrative Database [45].

## Results

### General Findings

We identified a total of 57 research items that met our eligibility criteria out of 1523 research items screened, including 46 research articles, seven dissertations, two GenBank entries, one laboratory report, and one abstract (see **Table 1** for a full list of eligible items and citations). Thirty-nine (68%) of the included items assessed MAYV infection in non-human vertebrates while 29 (51%) items assessed MAYV infection in arthropods. Of the 57 eligible items, 24 (42%) were included in the vertebrate seroprevalence meta-analysis, and the remaining items were only included in the qualitative analysis. A flow chart describing the article search and selection process is presented in **Fig 1**. Five articles were identified that met the inclusion criteria but were deemed to be reporting the same data as other included articles. These include de Thoisy *et al*., (2001) [46] and Talarmin *et al.*, (1998) [12] (both reporting the same data as de Thoisy *et al.,* (2003) [21]), Aitken *et al.,* (1960) [47] (reporting the same data as Aitken *et al.,* (1969) [48]), Batista *et al*., 2013 [49] (reporting the same data as Paulo *et al*., (2015) [50]), and Woodall (1967) [51] (reporting the same data as Taylor, (1967) [52]). These articles were excluded from this systematic review.

**Table 1.** Eligible Study Characteristics.

| Reference | Study Period | Country | Arthropods Tested (n) | Vertebrate non-human animals tested (n)<sup>a</sup> | MAYV infection reported |
| --- | --- | --- | --- | --- | --- |
| Aitken, 1969 [48] | 1953-1963 | Trinidad | 1,568,439 | --- | Yes |
| Araujo, 2003 [53] | 2002 | Brazil | --- | 555 | Yes |
| Araujo, 2004 [54] | 2003 | Brazil | --- | 202 | No |
| Araujo, 2004b [55] | 2003 | Brazil | --- | 495 | Yes |
| Araujo, 2012 [56] | 2007-2008 | Brazil | --- | 95 | Yes |
| Araujo, 2012b [57] | 2009 | Brazil | --- | 102 | Yes |
| Azevedo, 2009 [58] | 2008 | Brazil | 832 | --- | Yes |
| Batista, 2012 [59] | 2010 | Brazil | 122 | 65 | Yes |
| Calisher, 1974 [60] | 1967 | USA<sup>b</sup> | --- | 1,300 | Yes |
| Carrera, 2020 [61] | 2017 | Panama | 113 | --- | No |
| Casseb, 2010 [62] | 2009 | Brazil | --- | 2191 | Yes |
| Casseb, 2016 [63] | 2009 | Brazil | --- | 753 | Yes |
| Catenacci, 2017 [64] | 2006-2014 | Brazil | 239 | 142 | Yes |
| Cruz, 2009 [65] | 2006-2008 | Brazil | --- | 85 | No |
| Degallier, 1992 [66] | 1974-1988 | Brazil | 2,005,069 | 6,248 | Yes |
| De Thoisy, 2003 [21] | 1994-1995 | French Guiana | --- | 579 | Yes |
| Diaz, 2007 [67] | 1994 | Argentina, Paraguay | --- | 90 | No |
| Esposito, 2015 [68] | 1960 | Brazil | NA<sup>d</sup> | --- | Yes |
| Ferreira, 2020 [69] | 2017-2018 | Brazil | 10,569 | --- | Yes |
| Galindo, 1966 [70] | 1959-1962 | Panama | 377,492 | 2,444 | Yes |
| Galindo, 1967 [71] | 1966 | Panama | 11,829 | --- | Yes |
| Galindo, 1983 [72] | 1972-1979 | Panama | NA<sup>c</sup> | NA<sup>c</sup> | Yes |
| GenBank KY618129 | 1991 | Brazil | NA<sup>d</sup> | --- | Yes |
| GenBank KY618130 | 2011 | Brazil | NA<sup>d</sup> | --- | Yes |
| Gibrail, 2015 [73] | 2011-2014 | Brazil | --- | 50 | No |
| Gomes, 2019 [74] | 2018 | Brazil | --- | 213 | Yes |
| Groot, 1961 [75] | 1958-1960 | Colombia | 41,564 | --- | Yes |
| Groot, 1964 [11] | 1956-1961 | Colombia | --- | 34 | Yes |
| Henriques, 2008 [76] | 2002-2005 | Brazil | 37,519 | --- | No |
| Hoch, 1981 [19] | 1978-1979 | Brazil | 10,667 | 1785 | Yes |
| Kubiszeski, 2017 [77] | 2014-2015 | Brazil | 778 | --- | Yes |
| Laroque, 2014 [78] | 2008-2010 | Brazil | --- | 131 | Yes |
| Maia, 2019 [79] | 2017 | Brazil | 4786 | --- | Yes |
| Martinez, 2020 [80] | 2018-2019 | Colombia | 169 | --- | No |
| Medlin, 2016 [81] | 2005-2007 | Costa Rica | --- | 94 | No |
| Medina, 2015 [82] | 1999 | Venezuela | --- | NA<sup>d</sup> | Yes |
| Moreira-Soto, 2018 [83] | 2012-2017 | Brazil | --- | 103 | Yes |
| Nunes, 2009 [84] | 2005 | Brazil | --- | 181 | No |
| Paulo, 2015 [50] | 2012-2014 | Brazil | --- | 43 | Yes |
| Pauvolid-Correa, 2010 [85] | 2007 | Brazil | --- | 135 | No |
| Pauvolid-Correa, 2015 [86] | 2009-2011 | Brazil | --- | 748 | Yes |
| Pauvolid-Correa, 2008 [87] | 2007 | Brazil | 1,759 | NA<sup>e</sup> | No |
| Perez, 2019 [88] | 2007-2008 | Peru | --- | 90 | Yes |
| Pinheiro, 1974 [89] | 1971-1974 | Brazil | NA<sup>c</sup> | NA<sup>c</sup> | Yes |
| Pinheiro, 2019 [90] | 2017 | Brazil | 867 | --- | No |
| Powers, 2006 [91] | N/A | N/A | NA<sup>d</sup> | NA<sup>d</sup> | Yes |
| Price, 1978 [92] | 1972-1974 | Trinidad | --- | 997 | No |
| Ragan, 2019 [93] | N/A | N/A | --- | NA<sup>c</sup> | No |
| Sanmartin, 1973 [94] | 1967 | Colombia | 27,437 | 480 | No |
| Scherer, 1975 [95] | 1970-1971 | Peru | 1,500 | NA<sup>c</sup> | No |
| Serra, 2016 [96] | 2013 | Brazil | 4,556 | --- | Yes |
| Seymour, 1983 [20] | 1974-1976 | Panama | --- | 304 | Yes |
| Silva, 2017 [97] | 2016 | Brazil | 3,750 | --- | No |
| Srihongse, 1974 [98] | 1967 | Panama/Colombia | --- | 2026 | Yes |
| Tauro, 2019 [99] | 2017 | Brazil | 125 | --- | No |
| Taylor, 1967 [52] | N/A | Brazil/Trinidad | NA<sup>c</sup> | NA<sup>c</sup> | Yes |
| Turell, 2019 [100] | 2001-2002 | Peru | --- | 20 | No |

<sup>a</sup> Includes wild-caught, sentinel, and domestic animals.
<sup>b</sup> Migratory birds captured in Louisiana.
<sup>c</sup> Unable to determine the total number of animals or arthropods tested for MAYV.
<sup>d</sup> Genomic sequence only. No additional information provided.
<sup>e</sup> Horse seroprevalence data collected but recorded in another study.

**Fig 1.**
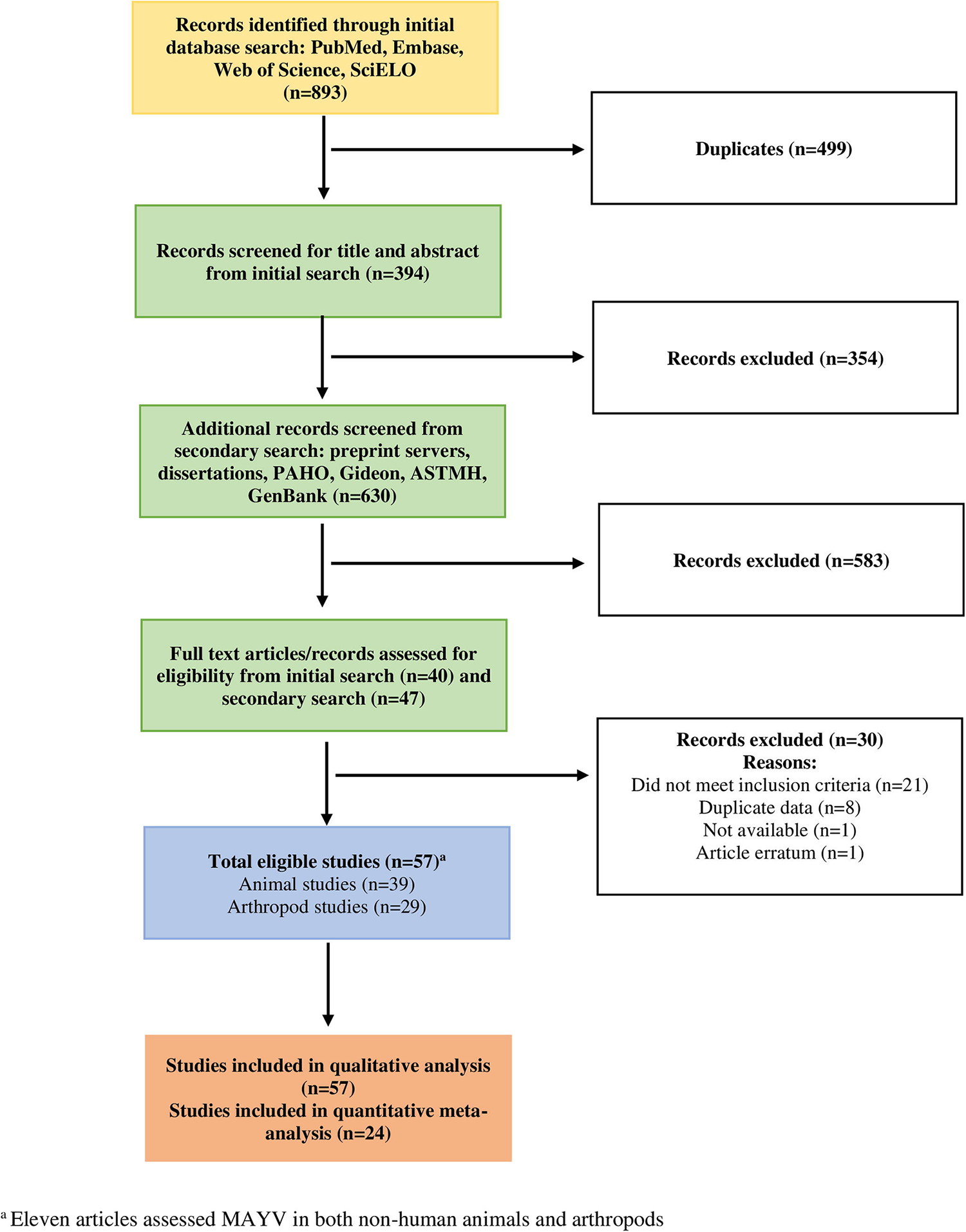
Flow diagram for search and selection of articles.

Studies were conducted in the following countries: Brazil (n=34), Panama (n=5), Colombia (n=4), Peru (n=3), Trinidad and Tobago (n=2), French Guiana (n=1), Venezuela (n=1), Costa Rica (n=1), and the United States of America (n=1). Several studies reported data from multiple countries including Argentina/Paraguay (n=1), Panama/Colombia (n=1), and Brazil/Trinidad and Tobago (n=1). The majority of studies were conducted after the year 2000 (n=33), although some studies were conducted between 1950-1969 (n=9), 1970-1989 (n=8), or 1990-1999 (n=4). Quality scores for all included studies are reported in Table 2.

**Table 2.**
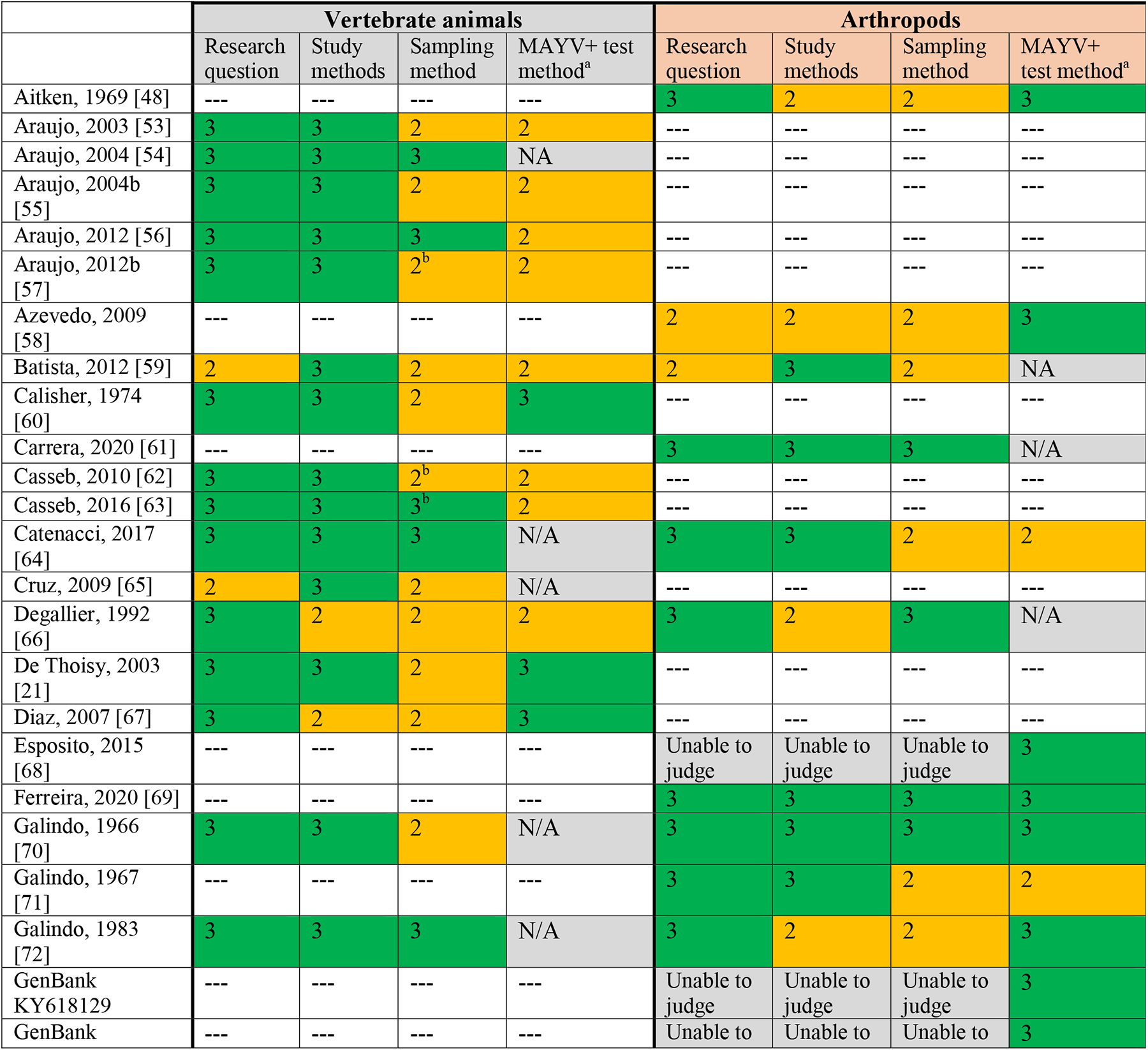
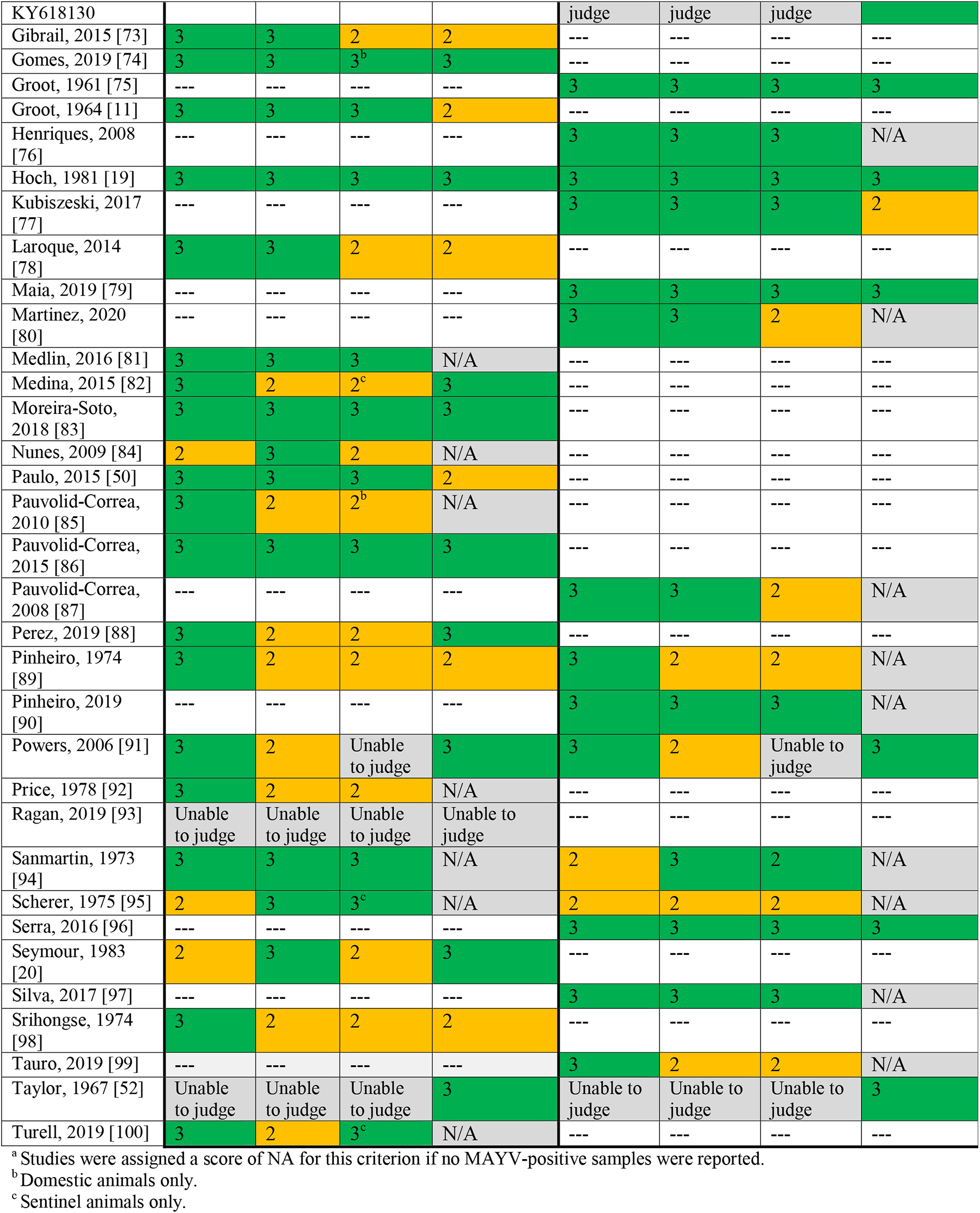
Quality Review Scores.

### MAYV in wild-caught non-human vertebrate animals

Thirty-nine (68%) studies in our systematic review assessed MAYV infection in wild-caught non-human vertebrate animals (including birds, mammals, and reptiles). Seventeen (44%) of these studies identified at least one non-human vertebrate that was positive for MAYV infection. Of the 27 taxonomic orders studied, 12 (44.4%) had evidence of MAYV infection: Artiodactyla (even-toed ungulates), Caprimulgiformes (nightbirds), Carnivora, Charadriiformes (shorebirds), Cingulata (armadillos), Columbiformes (pigeons and doves), Didelphimorphia (opossums), Passeriformes (passerine birds), Pilosa (sloths and anteaters), Primate, Rodentia, and Squamata (scaled reptiles). The greatest number of MAYV-positive animal species were found in the order Charadriiformes (n=16 positive species) and the order Primate (n=15 positive species). (See **S2 Table** for complete mammal data and **S3 Table** for complete avian data).

**Table 3** reports NHP species that were detected with MAYV antibodies. Only studies with positive results are shown on Table 3; other negative studies are listed in the **S2 Table**. High MAYV seroprevalence was confirmed by NT among *Alouatta seniculus* monkeys in individual studies in French Guiana [21] (n=51/98) and among *Callithrix argentata* monkeys in Brazil [19] (n=32/119). In addition, 29 *Cebus libidinosus* monkeys from wildlife screening centers were detected with MAYV antibodies according to HI, although only six were reported as monotypic reactions [78]. Diagnosis in these monkeys was not confirmed by NT. An additional *Cebus libidinosus* monkey presented a heterotypic reaction to MAYV (titer of 1:20) and four additional viruses according to HI (including a titer of 1:640 for Oropouche virus) [73]. However, based on the study’s protocol, confirmatory NT was only performed for viruses with titers > 1:40.

**Table 3.**
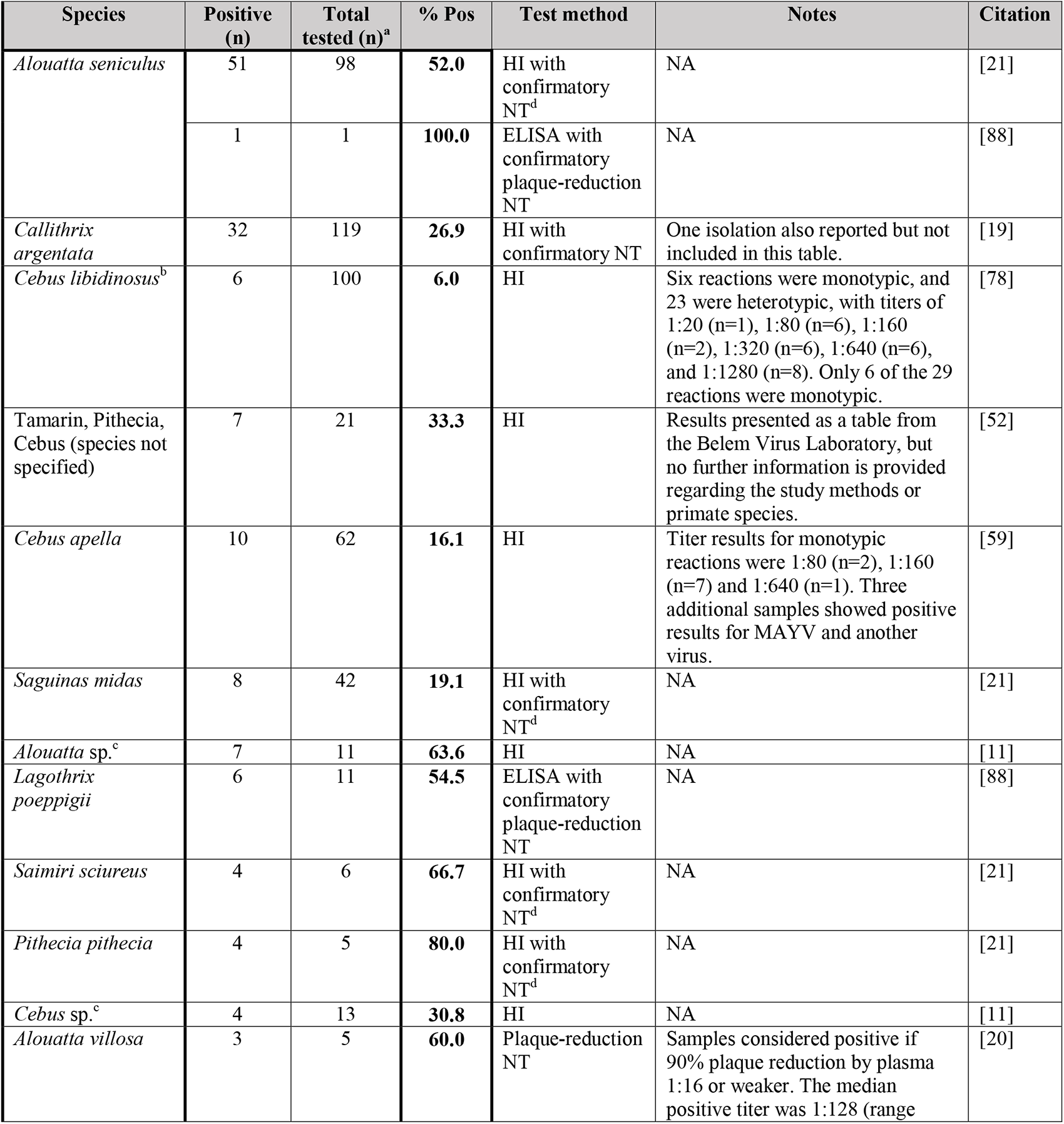
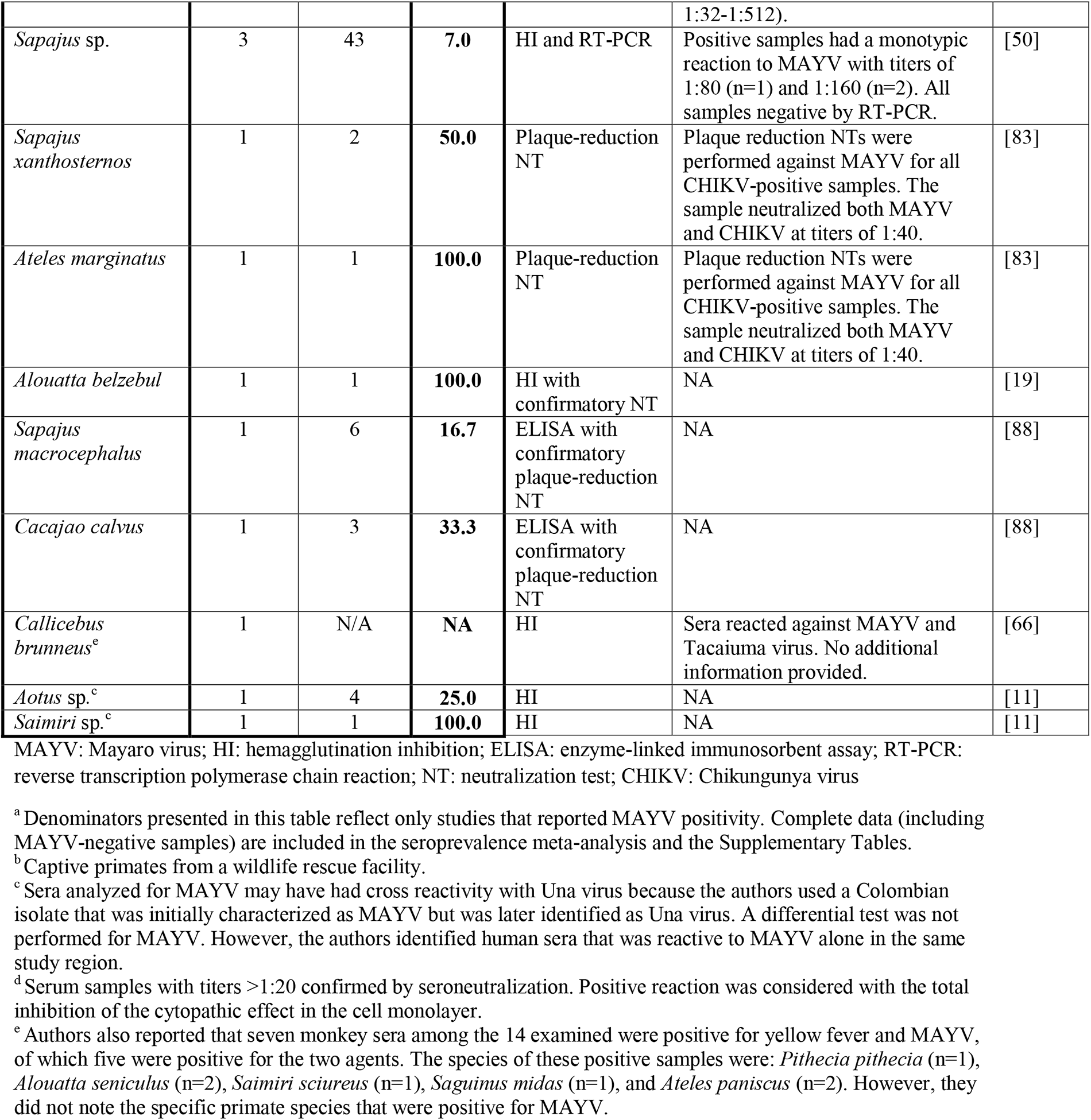
Evidence of MAYV infection in non-human primates.

Among the 12 additional NHP species with evidence of past MAYV infection, nine were confirmed by NT and three by HI alone. In addition, MAYV positivity was reported in the following NHP genera, although animals were not reported to species: *Aotus* (n=1/4), *Alouatta* (n=7/11), *Cebus* (n=4/13), *Sapajus* (n=3/43), and *Saimiri* (n=1/1). The authors reporting MAYV positivity in the *Aotus*, *Alouatta*, *Cebus*, and *Saimiri* genera noted that these results should be interpreted with caution due to potential for cross-reactivity with Una virus (UNAV) [11]. In one study conducted in Brazil, two of 11 Chikungunya virus (CHIKV)-positive serum samples (in the species *Sapajus xanthosternos* and *Ateles marginatus*) neutralized MAYV with titers of 1:40 in plaque reduction NTs [83]. These two samples were considered MAYV-positive and included in our meta-analysis. One additional study [67] detected neutralizing antibodies against both UNAV and MAYV in 21 *Alouatta caraya* monkeys. However, all 21 monkeys were diagnosed with UNAV based on a 4-fold titer difference between the two viruses. Therefore, we considered these monkeys MAYV-negative and did not include them in our meta-analysis. Finally, in 1963 the Belem Virus laboratory reported MAYV infection in seven NHPs based on HI tests alone [52]. These monkeys were described as Tamarin, Pithecia, and Cebus although no further information was provided regarding sampling method, testing protocol, or primate species.

MAYV antibodies were also detected in 21 bird species from the order Charadriiformes (n=16) and Passeriformes (n=5). All MAYV-positive birds were found in Brazil, with the exception of one MAYV isolate from a migrating bird captured in Louisiana USA [60]. A high MAYV-seroprevalence (n=34/122) was reported by the Belem Laboratory in 1963 among *Columbigallina* birds, although no additional information was provided regarding sampling method or bird species. MAYV antibodies were also detected in seven avian families that were not identified to genus or species. Only one study that detected MAYV antibodies in birds performed confirmatory NT [19]. All other diagnoses (with the exception of the virus isolation) were made by HI tests alone. See **Table 4** for additional information regarding avian species that were infected with MAYV.

**Table 4.**
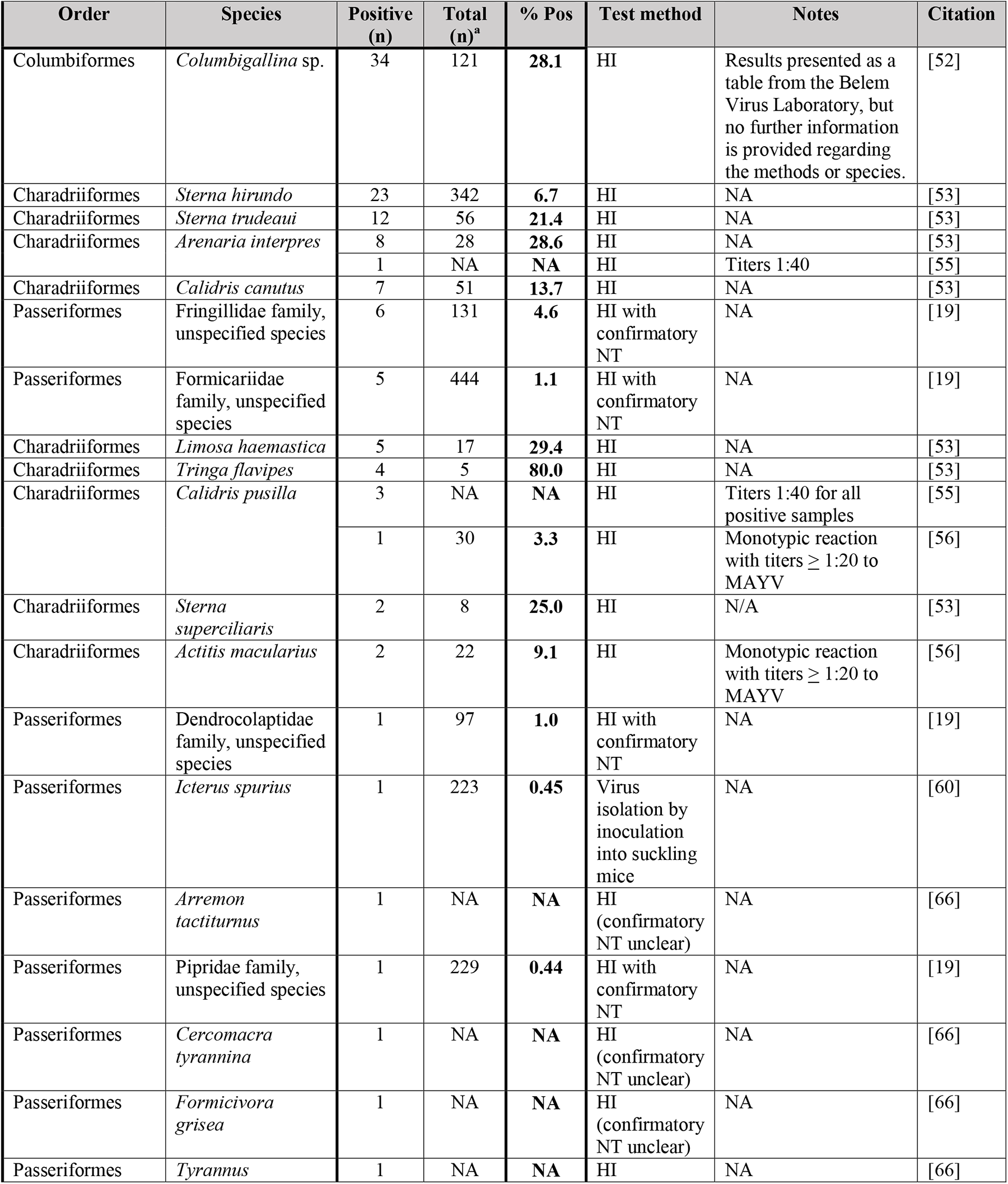
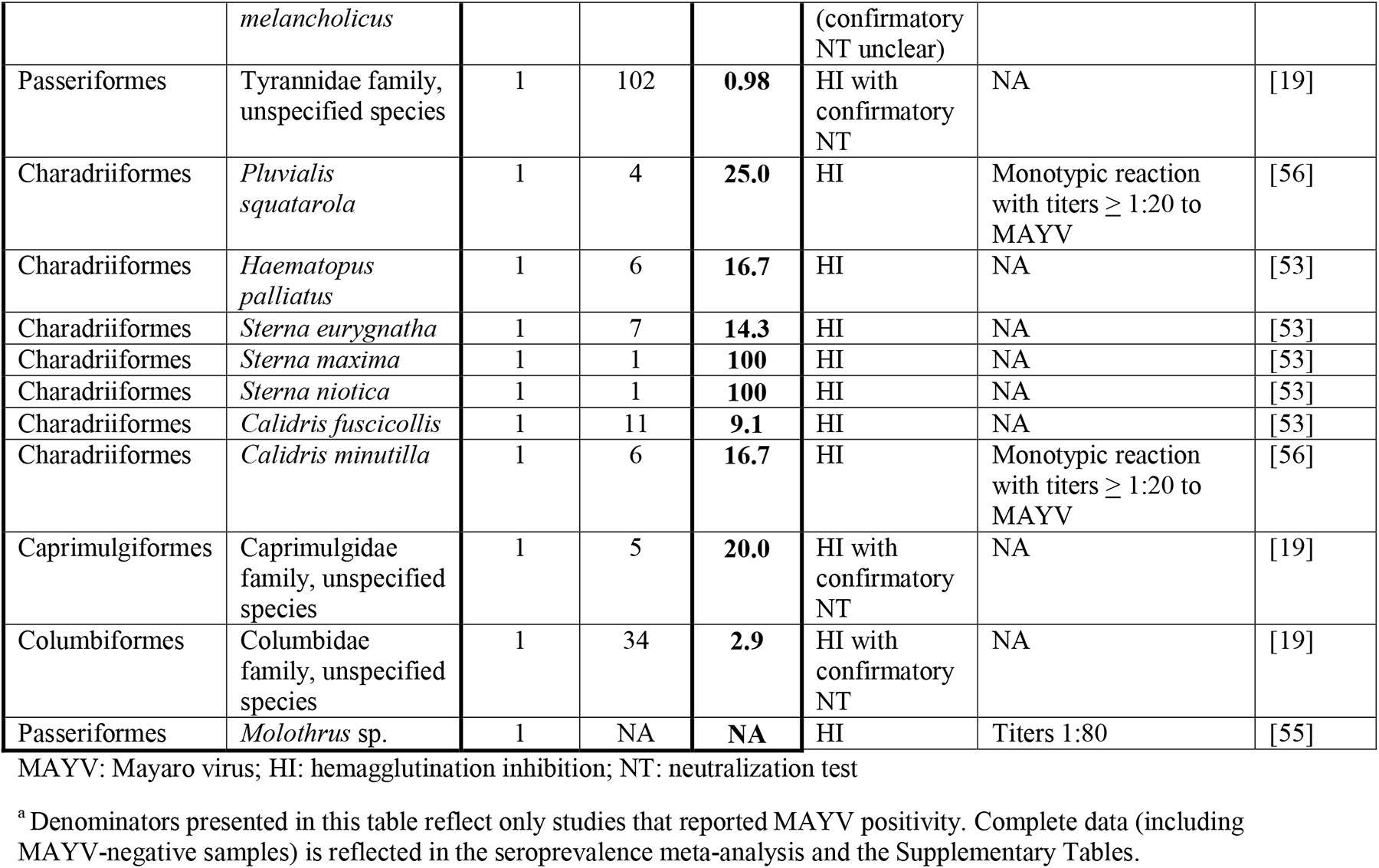
Evidence of MAYV infection in birds.

Additional wild-caught mammals with evidence of MAYV infection are presented in **Table 5**. Six rodent species as well as unidentified rodents in the *Echimys* and *Proechimys* genera were detected with MAYV antibodies in French Guiana [21], Peru [88], and Panama [20]. In addition, four species in the order Didelphimorphia, three species in the order Pilosa, and one species each in the orders Carnivora, Artiodactyla, and Cingulata were detected with MAYV antibodies in French Guiana [21] and Peru [88]. Additional positive samples were detected in the orders Rodentia, Didelphimorphia, and Pilosa although the species were not identified.

**Table 5.** Evidence of MAYV infection in mammals (excluding non-human primates)

| Order | Species | Positive (n) | Total (n)<sup>a</sup> | % Pos | Test method | Notes | Citation |
| --- | --- | --- | --- | --- | --- | --- | --- |
| Rodentia | Wild rodents, unspecified | 71 | 960 | **7.4** | HI | Results presented as a table from the Belem Virus Laboratory, but no further information is provided regarding the methods or species. | [52] |
| Didelphimorphia | Opossum, unspecified | 9 | 122 | **7.4** | HI | Results presented as a table from the Belem Virus Laboratory, but no further information is provided regarding the methods or species. | [52] |
| Pilosa | *Choloepus didactylus* | 7 | 26 | **26.9** | HI with confirmatory NT<sup>b</sup> | NA | [21] |
| Didelphimorphia | *Marmosa* sp. | 7 | 46 | **15.2** | HI | NA | [52] |
| Pilosa | *Tamandua tetradactyla* | 6 | 26 | **23.1** | HI with confirmatory NT<sup>b</sup> | NA | [21] |
| Cingulata | *Dasypus novemcinctus* | 4 | 40 | **10.0** | HI with confirmatory NT<sup>b</sup> | NA | [21] |
|  |  | 2 | 4 | **50.0** | ELISA with confirmatory plaque reduction NT | NA | [88] |
| Rodentia | *Dasyprocta leporina* | 5 | 29 | **17.2** | HI with confirmatory NT<sup>b</sup> | NA | [21] |
| Didelphimorphia | *Philander opossum* | 5 | 27 | **18.5** | HI with confirmatory NT<sup>b</sup> | NA | [21] |
| Rodentia | *Coendou prehensilis* | 3 | 26 | **11.5** | HI with confirmatory NT<sup>b</sup> | NA | [21] |
| Rodentia | *Dasyprocta punctata* | 3 | 5 | **60.0** | Plaque reduction NT | Samples considered positive if 90% plaque reduction by plasma 1:16 or weaker. The median positive titer was 1:128 (range 1:32-1:512). | [20] |
| Rodentia | *Dasyprocta fuliginosa* | 3 | 27 | **11.1** | ELISA with confirmatory plaque reduction NT | NA | [88] |
| Rodentia | *Coendou melanurus* | 2 | 15 | **13.3** | HI with confirmatory NT<sup>b</sup> | NA | [21] |
| Didelphimorphia | *Didelphis albiventris* | 2 | 19 | **10.5** | HI with confirmatory NT<sup>b</sup> | NA | [21] |
| Rodentia | *Echimys* sp. | 1 | 21 | **4.8** | HI with confirmatory NT<sup>b</sup> | NA | [21] |
| Rodentia | *Agouti paca* | 1 | 10 | **10.0** | ELISA with confirmatory plaque reduction NT | NA | [88] |
| Rodentia | *Proechimys* sp. | 1 | 18 | **5.6** | HI with confirmatory NT<sup>b</sup> | NA | [21] |
| Didelphimorphia | *Caluromys philander* | 1 | 5 | **20.0** | HI with confirmatory NT<sup>b</sup> | NA | [21] |
| Didelphimorphia | *Didelphis marsupialis* | 1 | 29 | **3.5** | HI with confirmatory NT<sup>b</sup> | NA | [21] |
| Carnivora | *Potos flavus* | 1 | 9 | **11.1** | HI with confirmatory NT<sup>b</sup> | NA | [21] |
| Artiodactyla | *Pecari tajacu* | 1 | 6 | **16.7** | ELISA with confirmatory plaque reduction NT | NA | [88] |
| Pilosa | *Bradypus tridactylus* | 1 | 29 | **3.5** | HI with confirmatory NT<sup>b</sup> | NA | [21] |
| Pilosa | *Bradypus* sp. | 1 | 3 | **33.3** | HI | NA | [52] |

MAYV: Mayaro virus; HI: hemagglutination inhibition; ELISA: enzyme-linked immunosorbent assay; RT-PCR: reverse transcription polymerase chain reaction; NT: neutralization test
<sup>a</sup> Denominators presented in this table reflect only studies that reported MAYV positivity. Complete data (including MAYV-negative samples) is reflected in the seroprevalence meta-analysis and the Supplementary Tables.
<sup>b</sup> Serum samples with titers >1:20 confirmed by seroneutralization. Positive reaction was considered with the total inhibition of the cytopathic effect in the cell monolayer.

Successful isolation of MAYV was reported from the following viremic animals: a silvery marmoset (*Callithrix argentata*) captured during a MAYV outbreak in Belterra, Brazil [19] and a migrating orchard oriole (*Icterus spurius*) captured in Louisiana [60]. In addition, the Belem Virus Laboratory reported MAYV isolation from two lizard species in 1963 [52] (*Tropidurus torquatus* and *Ameiva ameiva*) although no further information was provided regarding study methods or procedures.

The geographic distribution of animals (wild-caught, domestic, and sentinel) infected with MAYV is presented in **Fig 2**. The infected animals were identified in six countries overall, including Brazil, Peru, French Guiana, Colombia, Venezuela, and Panama, although the majority of infected animals were found in Brazil. Overall, 12 locations were geo-referenced as points, four locations as ADM1 polygons, 15 locations as ADM2 polygons, and two locations as custom polygons.

**Fig 2.**
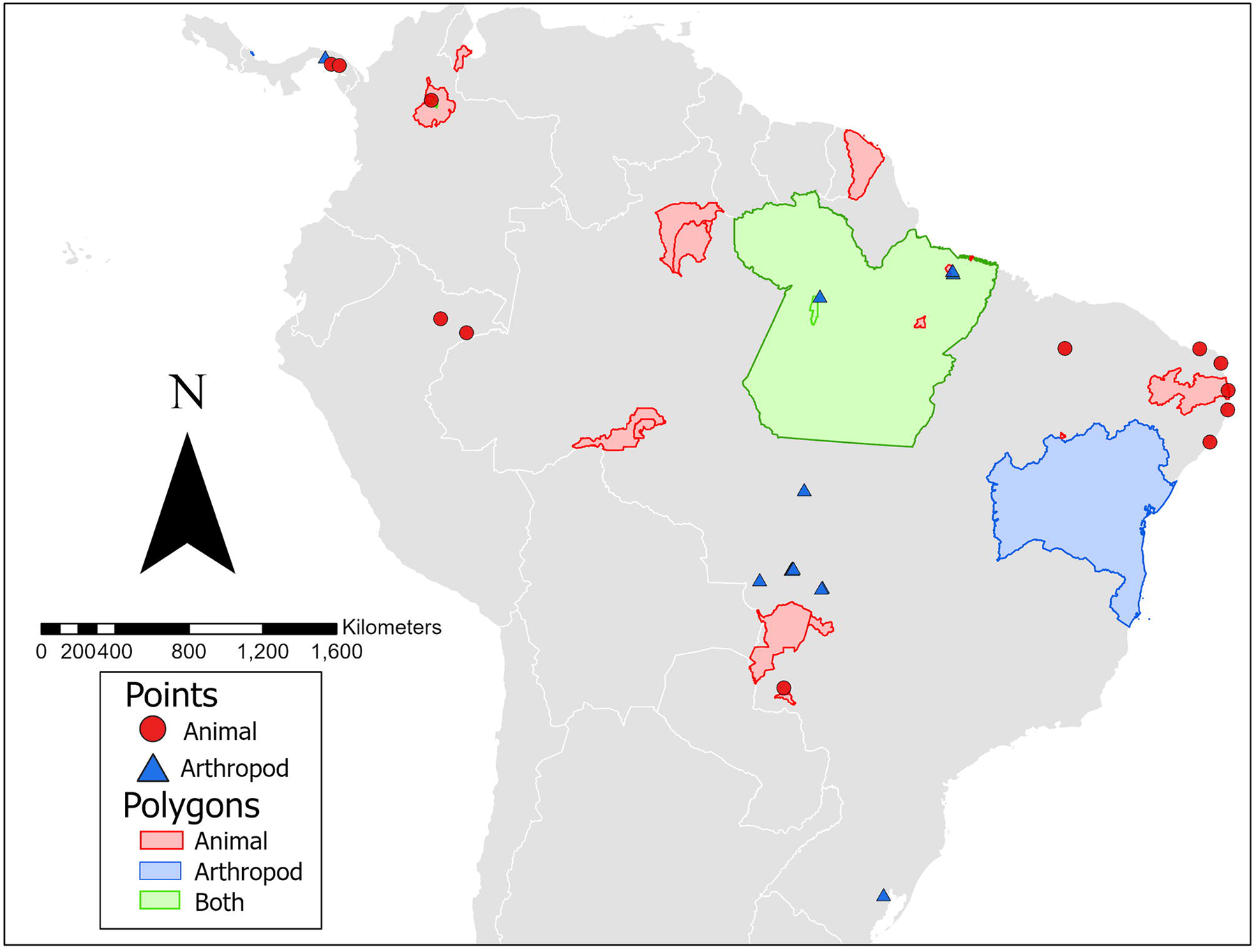
Georeferenced locations of MAYV positivity in non-human animals and arthropods. The finest spatial scale is presented where possible. One MAYV isolate detected in a migrating bird in Louisiana is not included in the map.

### MAYV in domestic or sentinel animals

Nine studies analyzed MAYV seroprevalence in domestic animals (equids, sheep, poultry, dogs, pigs, cattle, and buffaloes), and five studies analyzed MAYV seroprevalence in sentinel animals (monkeys, mice, and hamsters). Domestic and sentinel animals with evidence of MAYV positivity are reported in **Table 6** and complete results are reported in the **S4 Table**. In domestic animals, evidence of MAYV infection was detected in equids, cattle/buffalo, and dogs. Six studies assessed MAYV seroprevalence in Brazilian equids [54, 57, 63, 74, 85, 86], and antibodies against MAYV were detected in four of these studies. Notably, Gomes et al. [74] reported MAYV neutralizing antibodies in 48 equids out of 213 (23%) based on ELISA. However, only 16 of the 48 equids were considered positive based on the study’s diagnostic criterion of 4-fold greater plaque reduction NT_90_ titer than that of the other viruses under study. In addition, Casseb et al. [63] detected MAYV antibodies in 40 horses using HI, although only four of the 40 reactions were monotypic, and confirmatory NTs were not performed. Additional domestic animals with evidence of MAYV infection included cattle/buffalo (n=14/1103 positive reactions by HI; 5/14 monotypic reactions [62]) and dogs (n=2/7 positive reactions by HI [53]). In addition, neutralizing antibodies (plaque reduction NT_90_ titer >10) against MAYV were detected in three sheep in Brazil [86]. However, these animals did not meet the original study’s diagnostic criterion for MAYV diagnosis based on 4-fold greater plaque reduction NT_90_ titer than that of the other viruses under study. Evidence of MAYV infection was also detected by HI in two sentinel monkeys placed in the tree canopy in Panama [98], and one MAYV isolate was obtained from a sentinel hamster in Venezuela [82].

**Table 6.** Domestic and sentinel animals with evidence of MAYV infection.

| Animal Type | Total Positive | Number Tested<sup>a</sup> | % Pos | Test Method | Notes | Citation |
| --- | --- | --- | --- | --- | --- | --- |
| Domestic Equids | 16 | 213 | **7.5** | ELISA with confirmatory plaque reduction NT | Forty-eight horses had antibodies to MAYV by ELISA. Sixteen of 48 (33%) were considered positive by plaque reduction NT<sub>90</sub> for MAYV with titers 1:10 (n=12), 1:20 (n=3) and 1:40 (n=1). | [74] |
|  | 4 | 753 | **0.5** | HI | Forty reactions overall. Four of 40 reactions were monotypic while 36 of 40 were heterotypic. | [63] |
|  | 11 | 102 | **10.8** | HI | Not clear if the 11 reactions are monotypic or heterotypic. | [57] |
| | 10 | 748 | **1.5** | Plaque reduction NT | Forty-four horses had neutralizing antibody (titer ≥ 10) against MAYV, but only ten met the diagnostic criteria of 4-fold greater plaque reduction NT<sub>90</sub> titer than the three other viruses (VEEV, EEEV, WEEV). Positive samples had titers of 1:20 (n=6) and 1:40 (n=4) | [86] |
| Domestic Cattle/Buffalo | 5 | 1103 | **0.5** | HI | Positive reactions were considered any reaction with a titer equal to or greater than 1:20. Fourteen reactions overall, and five of 14 reactions were monotypic. | [62] |
| Domestic Dog | 2 | 7 | **28.6** | HI | N/A | [53] |
| Sentinel Hamster | 1 | N/A | **N/A** | RT-PCR |  | [82] |
| Sentinel Monkeys | 2 | 13 | **15.4** | HI | N/A | [98] |

MAYV: Mayaro virus; HI: hemagglutination inhibition; ELISA: enzyme-linked immunosorbent assay; RT-PCR: reverse transcription polymerase chain reaction; NT: neutralization test; VEEV: Venezuelan equine encephalitis virus; EEEV: Eastern equine encephalitis virus; WEEV: Western equine encephalitis virus
<sup>a</sup> Denominators presented in this table reflect only studies that reported MAYV positivity. Complete data (including MAYV-negative samples) are reflected in the seroprevalence meta-analysis and the Supplementary Tables.

### Pooled prevalence of MAYV in non-human vertebrate animals

Twenty-four studies overall were included in the pooled prevalence meta-analysis. Eight studies were excluded because they did not clearly state how many animals were tested for MAYV within each order [55, 66, 72, 84, 89, 93] or did not present serologic results [60, 70]. Another study was excluded because authors reported the number of “Group A” positive serum samples, but did not specify individual viruses [98]. Studies were also excluded if they only reported sequence data or only included sentinel animals [82, 91, 95, 100]. Finally, a study that sampled bats exclusively was excluded because no MAYV-positive samples were reported in the order Chiroptera [92].

Eleven orders of nonhuman vertebrate animals (including domestic equids) were included in the meta-analysis. Orders were excluded from the analysis due to insufficient sample size (N<10) or if no MAYV-positive samples were reported. These include the orders Apodiformes (MAYV prevalence: 0/3), Caprimulgiformes (MAYV prevalence: 1/6), Chiroptera (MAYV prevalence: 0/1546), Crocodilia (MAYV prevalence: 0/87), Cuculiformes (MAYV prevalence: 0/5), Galliformes (MAYV prevalence: 0/1), Gruiformes (MAYV prevalence: 0/2), Psittaciformes (MAYV prevalence: 0/3), Tinamiformes (MAYV prevalence: 0/2), Pelecaniformes (MAYV prevalence: 0/2), and Podicipediformes (MAYV prevalence: 0/2).

The primate order appeared in 14 studies that were included in the meta-analysis. When all positive samples were included, the pooled MAYV seroprevalence among primates was 13.1% (95% CI: 4.3-25.1%) according to the random effects model, with statistically significant heterogeneity across studies (*I^2^* = 95%, p < 0.01). After excluding positive samples that were not confirmed by NT, the pooled MAYV seroprevalence among primates decreased to 4.9 (95% CI: 0.0-15.2; *I^2^* = 96%; p < 0.01) according to the random effects model. When the analyses were repeated using the GLMM with logit transformation, seroprevalence estimates for primates decreased to 8.7% (95% CI: 3.1-22.0%) overall and to 0.7% (95% CI: 0.0-9.1%) when only NT-positive samples were included. Additional meta-analysis results for the various primate genera are presented in **S6 and S7 Tables**. The seroprevalence for the most frequently sampled primate genera was 32.2% (95% CI: 0.0-79.2%) for the *Alouatta* genus, 17.8% (95% CI: 8.6-28.5%) for the *Callithrix* genus, and 3.7% (95% CI: 0.0-11.1%) for the *Cebus/Sapajus* genus.

Meta-analysis results for additional non-human vertebrate orders are presented in **Table 7** and forest plots for mammal orders and avian orders are presented in **Figs 3 and 4**, respectively. When all positive samples were included in the analysis, the highest seroprevalence was observed in the orders Charadriiformes (prevalence: 7.1%; 95% CI: 2.2-13.8%) and Cingulata (prevalence: 3.0%; 95% CI: 0.0-24.5%). When the analysis was repeated using GLMM with logit transformation, the seroprevalence increased to 10.0% (95% CI: 2.7-30.8%) for the order Cingulata and 9.2% (95% CI: 4.4-18.2%) for the order Charadriiformes. All results of the sensitivity analysis using GLMM with logit transformation are reported in the **S5 Table**. An additional sensitivity analysis using fixed effects models is presented in the **S8 and S9 Tables**.

**Table 7.** Pooled Prevalence Table (Random effects with Freeman-Tukey double arcsine transformation)

| Order | Positives Included<sup>a</sup> | Studies (n) | Total (n) | Positive (n) | Pooled Prevalence (%) | 95% CI | I<sup>2</sup> (%) | $\tau^2$ | p-value |
| --- | --- | --- | --- | --- | --- | --- | --- | --- | --- |
| *Mammals* |  |  |  |  |  |  |  |  |  |
| Primate | HI and NT | 13 | 897 | 153 | 13.1 | 4.3; 25.1 | 95 | 0.0692 | <0.01 |
|  | NT only | 13 | 858 | 114 | 4.9 | 0.0; 15.2 | 96 | 0.0851 | <0.01 |
| Pilosa | HI and NT | 7 | 297 | 15 | 0.0 | 0.0; 6.6 | 84 | 0.0338 | <0.01 |
|  | NT only | 7 | 296 | 14 | 0.0 | 0.0; 3.9 | 82 | 0.0305 | <0.01 |
| Rodentia | HI and NT | 7 | 1557 | 90 | 1.3 | 0.0; 6.5 | 91 | 0.0160 | <0.01 |
|  | NT only | 7 | 1486 | 19 | 0.1 | 0.0; 3.7 | 90 | 0.0153 | <0.01 |
| Domestic Equids | HI and NT | 6 | 1955 | 41 | 1.1 | 0.0; 4.5 | 90 | 0.0085 | <0.01 |
|  | NT only | 6 | 1940 | 26 | 0.0 | 0.0; 1.9 | 90 | 0.0087 | <0.01 |
| Didelphimorphia | HI and NT | 6 | 369 | 25 | 2.0 | 0.0; 7.2 | 68 | 0.0101 | <0.01 |
|  | NT only | 6 | 353 | 9 | 0.1 | 0.0; 4.2 | 74 | 0.0141 | <0.01 |
| Carnivora Order | HI and NT | 5 | 40 | 2 | 0.1 | 0.0; 8.1 | 0 | 0 | 0.71 |
|  | NT only | 5 | 40 | 2 | 0.1 | 0.0; 8.1 | 0 | 0 | 0.71 |
| Cingulata Order | HI and NT | 4 | 70 | 6 | 3.0 | 0.0; 24.5 | 35 | 0.0198 | 0.20 |
|  | NT only | 4 | 70 | 6 | 3.0 | 0.0; 24.5 | 35 | 0.0198 | 0.20 |
| Artiodactyla | HI and NT | 2 | 26 | 1 | 2.3 | 0.0; 20.7 | 46 | 0.0172 | 0.17 |
|  | NT only | 2 | 26 | 1 | 2.3 | 0.0; 20.7 | 46 | 0.0172 | 0.17 |
| **Birds<sup>b</sup>** |  |  |  |  |  |  |  |  |  |
| Charadriiformes | HI and NT | 3 | 641 | 71 | 7.1 | 2.2; 13.8 | 61 | 0.0045 | 0.08 |
| Passeriformes | HI and NT | 4 | 1166 | 14 | 0.0 | 0.0; 0.0 | 27 | 0.0010 | 0.25 |
| Columbiformes | HI and NT | 4 | 171 | 35 | 2.2 | 0.0; 27.1 | 87 | 0.0591 | <0.01 |

MAYV: Mayaro virus; HI: hemagglutination inhibition; NT: neutralization test; CI: confidence interval
<sup>a</sup> The first analysis (HI and NT) included all positive samples, regardless of test method. A sensitivity analysis was conducted that included only positive samples that were confirmed with NT.
<sup>b</sup> Only one study reporting MAYV positivity in birds used confirmatory NT. Therefore, a sensitivity analysis was not conducted.

**Fig 3.**
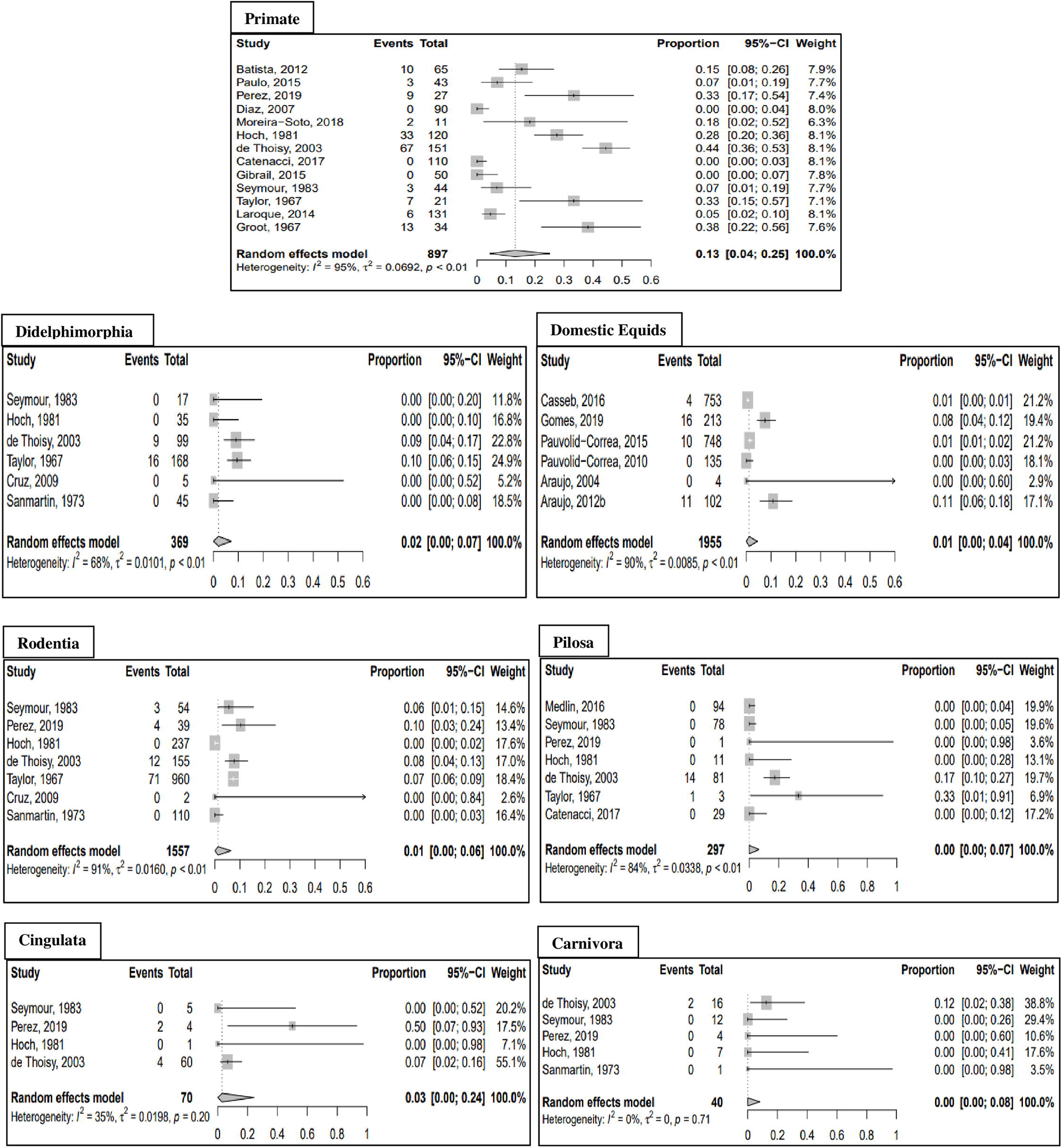
Forest plots of mammal orders from meta-analysis of pooled MAYV seroprevalence. Estimates are based on random effects model with Freeman-Tukey double arcsine transformation. All samples that tested MAYV-positive are included, regardless of test method.

**Fig 4.**
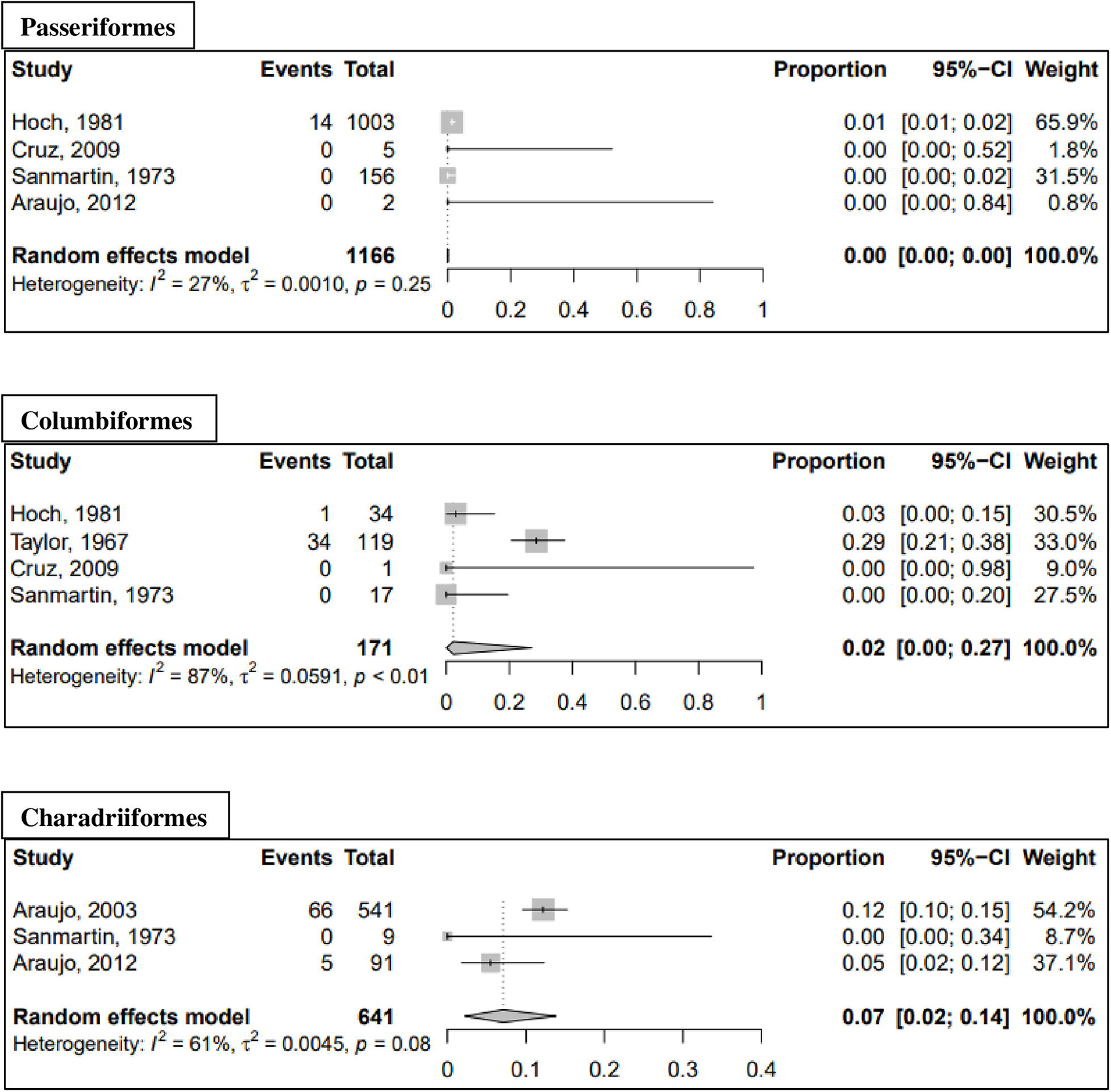
Forest plots of avian orders from meta-analysis of pooled MAYV seroprevalence. Estimates are based on random effects model with Freeman-Tukey double arcsine transformation. All samples that tested MAYV-positive are included, regardless of test method.

### MAYV in wild-caught arthropods

Twenty-eight of the studies in our systematic review analyzed MAYV infection in wild-caught arthropods. Seventeen (61%) of the 28 studies reported at least one arthropod that was positive for MAYV infection. Of the mosquito genera studied, seven were found to be infected with MAYV: *Aedes, Culex, Haemagogus, Psorophora, Sabethes, Wyeomyia*, and *Mansonia*. For detailed information on all infected mosquito species, see **Table 8**. The majority of infected vectors were identified using viral isolation techniques, although three studies reported MAYV positivity using RT-PCR alone. In addition, one study reported isolation of MAYV from an *Ixodes* tick [91] while another study reported isolation from a *Gigantolaelaps* mite [52]. Complete results, including studies that did not detect MAYV in arthropods, are reported in the **S10 Table.**

**Table 8.**
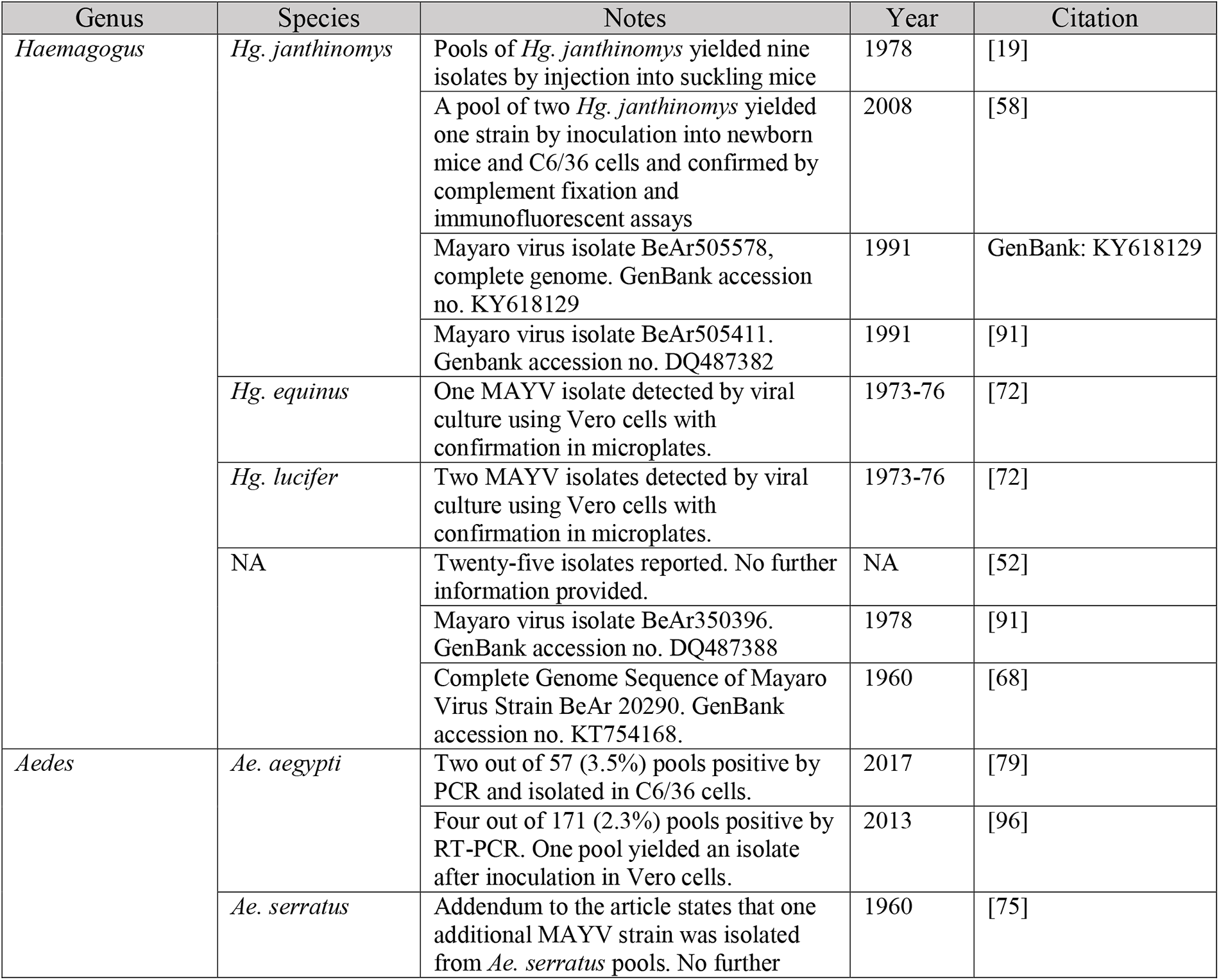
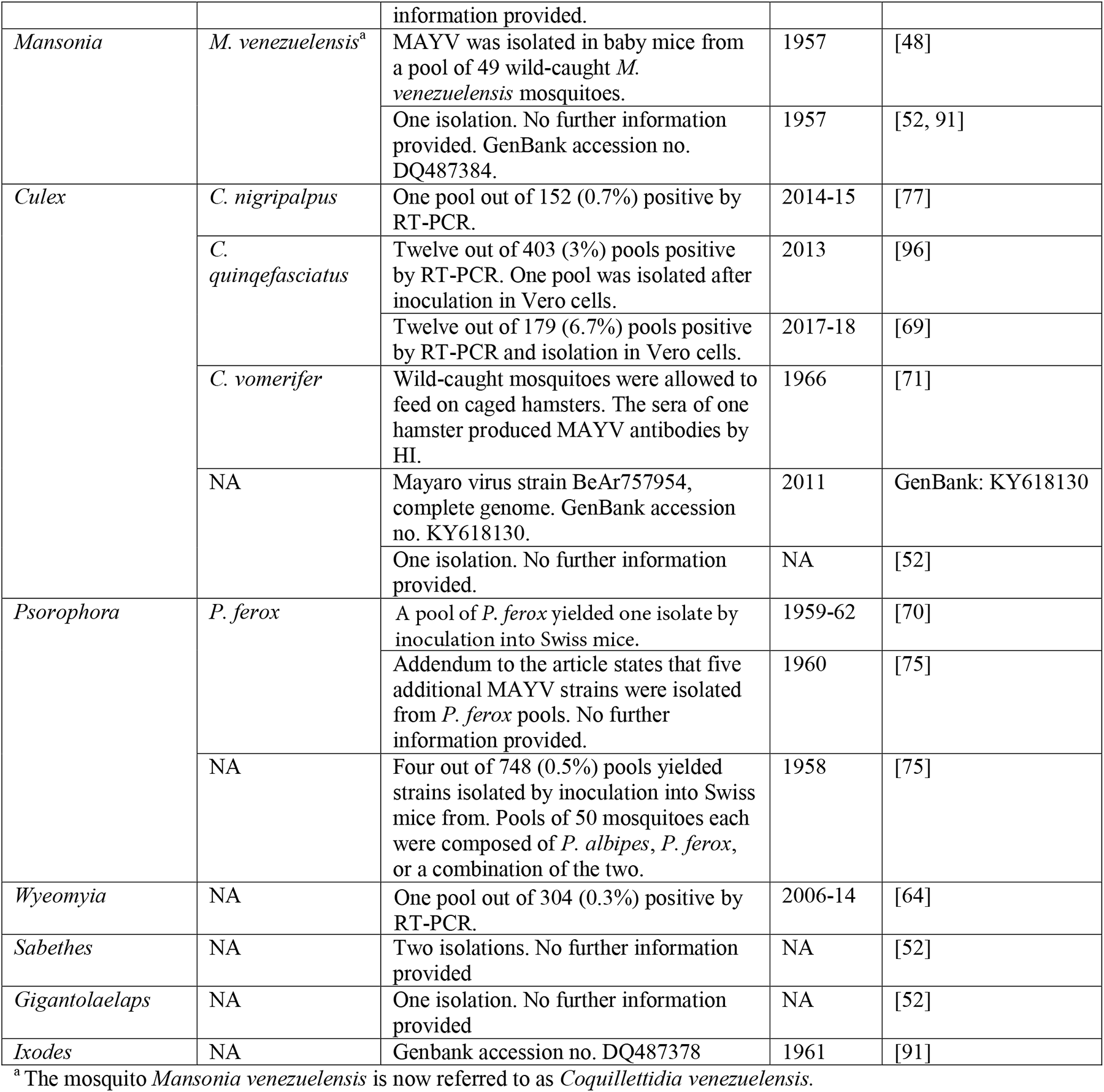
Evidence of MAYV infection in arthropods.

The geographic distribution of vectors infected with MAYV is presented in **Fig 2**. MAYV-positive arthropods were identified in four countries overall, including Brazil, Colombia, Panama, and Trinidad. Overall, 15 locations were geo-referenced as points, two locations as ADM1 polygons, two locations as ADM2 polygons, two locations as ADM3 polygons, and two as custom polygons.

### Analysis of publication bias

Publication bias was assessed among six animal orders (including domestic equids) and two primate genera. The results of Egger’s test did not reveal evidence of publication bias for the included studies. Therefore, the Trim fill technique was not carried out. Funnel plots are presented in **S1 and S2 Figures**, and results of Egger’s test are reported in the **S11 Table**.

## Discussion

To our knowledge, this study is the first attempt to systematically review the existing evidence of non-human animal reservoirs and arthropod vectors of MAYV, and the first study to quantitatively analyze the pooled seroprevalence of potential reservoirs. We identified 57 studies that assessed MAYV infection in non-human vertebrate animals and arthropods. Overall, the studies found evidence of MAYV infection in 12 wild-caught animal orders and seven arthropod genera across seven Latin American countries and the USA.

The majority of animal species that were found to be infected with MAYV belonged to the orders Primate and Charadriiformes (shorebirds). Several MAYV-positive species were also detected in the orders Rodentia, Didelphimorphia, and Pilosa. Overall, the highest MAYV pooled prevalence occurred in the Primate order. This finding points to the potential role of NHPs as an important reservoir in the MAYV transmission cycle.

The role of NHPs in sylvatic transmission cycles of arboviruses has been demonstrated with varying degrees of evidence [101]. Several arboviruses have been successfully isolated from wild NHPs, including dengue [102], CHIKV [103], and Zika [104] viruses. While isolation of a virus from NHPs is important for establishing the existence of a sylvatic cycle, it is difficult to achieve due to the short duration of viremia [101]. In our review, we identified only one study that successfully isolated MAYV from a NHP [19]. In the absence of viral detection, antibody seroprevalence has been used as evidence of the role of NHPs in sylvatic transmission cycles [105, 106]. Therefore, the high seroprevalence of MAYV among NHPs, including 52% seropositivity among *A. seniculus* monkeys in a 1994-95 survey in French Guiana [21], points to the potential importance of NHPs as MAYV reservoirs. Furthermore, Hoch et al. [19] reported substantial viremia in *C. argentata* marmosets that were experimentally infected with MAYV and noted that viremia titer was likely sufficient to infect vectors. Due to the high MAYV seroprevalence among marmosets during the Belterra outbreak, the isolation of MAYV from a single *C. argentata* marmoset, and the results of experimental infection studies, the authors concluded that marmosets were likely the amplifying hosts of MAYV.

The importance of birds in the MAYV transmission cycle was hypothesized following viral isolation from a migrating oriole (*Icterus spurius*) in Louisiana [60]. Avian species have been implicated as definitive or potential reservoirs of several Alphaviruses, including Sindbis virus [107], Ross River virus [108], and Eastern/Western equine encephalitis virus [109]. However, their role in MAYV transmission remains poorly understood. Our systematic review identified seven studies that found MAYV positivity in birds in the orders Passeriformes, Caprimulgiformes, Columbiformes, and Charadriiformes with relatively high seroprevalence reported in several bird species in the latter two orders [52, 53]. While some have theorized that MAYV has been introduced into certain areas by migratory birds [59], this hypothesis requires further study in order to elucidate the role of birds in MAYV transmission.

Although evidence of MAYV infection was detected in several vertebrate species, identifying the primary non-human animal reservoirs remains a difficult task. The precise definition of a disease “reservoir” has been a source of disagreement [17, 110]. One definition proposed by Haydon et al., (2002) defined a reservoir as “one or more epidemiologically connected populations or environments in which the pathogen can be permanently maintained and from which infection is transmitted to the defined target population” [17]. In addition, in 2005 Kuno and Chang outlined three basic criteria for the identification of reservoirs including isolation of the virus from the suspected reservoir population, high antibody prevalence in field-caught animals, and evidence of viremia in laboratory settings, although they posited that definitive identification of a reservoir requires evidence of long-term infection [111]. The role of various non-human vertebrates in the MAYV transmission cycle should be explored further in longitudinal seroprevalence surveys and experimental transmission studies in laboratory settings.

The sylvatic *Hg. janthinomys* mosquito has long been considered as the primary vector of MAYV. This is in part based on the isolation of MAYV from several pools of *Hg. janthinomys* mosquitoes in the context of a major MAYV outbreak in Belterra, Brazil in 1978 [19]. Our systematic review also identified several additional mosquito species including *Ae. aegypti* and *Cx. quinqefasciatus* with evidence of MAYV infection. A caveat, however, is that the isolation of a virus or detection of viral RNA through PCR is not sufficient to establish that arthropod as a biological vector [112], i.e. involved in the biological transmission of pathogens [111]. The World Health Organization (WHO) established three criteria to define a confirmed vector: (1) viral isolation in the absence of vertebrate blood; (2) biological transmission of the virus in experimental conditions; and (3) presence of certain temporal, geographic and other epidemiological or ecological parameters that allow transmission to occur [112]. Thus, certain arthropods that are capable of ingesting and transmitting a virus may not be established as confirmed vectors if the other parameters are not in place.

Experimental transmission studies support the role of *Ae. aegypti* as a possible MAYV vector with high MAYV infection rates and transmission potential [22-24, 113]. For example, Long et al., revealed *Ae. aegypti* to be a capable MAYV vector with a relatively short extrinsic incubation period [22]. Furthermore, MAYV titers in the saliva of *Ae. aegypti* were similar to other Alphavirus-vector systems including EEEV in *Culiseta melanura* and VEEV in *Ae. albopictus* and *Ae. taeniorhynchus.* In contrast, *Cx. quinquefasciatus* mosquitoes exhibited low MAYV infection rates and inability to transmit MAYV in laboratory settings [113]. It is also important to note that the competence of a given vector species to transmit MAYV may be impacted by the MAYV genotype that is present in a given area. In laboratory conditions, genotype L infection rates were significantly higher than genotype D infection rates among *Ae. aegypti* mosquitoes [113].

The spillover of MAYV into urban populations has been a source of concern for Latin American health authorities [114]. The implication is that anthropophilic, urban-dwelling mosquitoes like *Ae. aegypti* as effective vectors of MAYV would increase the potential for urban MAYV outbreaks [115]. Concerns of urban MAYV transmission were amplified after antibodies to MAYV were discovered in 33 of 631 sera (5.2%) in the city of Manaus, Brazil in 2007-08 [25] although it is unclear if humans can serve as amplification hosts. For example, Long et al. noted that the short duration of MAYV viremia and the relatively low viremic titers in humans reduces the probability of urban spread [22]. Our systematic review identified two recent studies conducted in the city of Cuiaba in which MAYV was isolated from pools of wild-caught *Ae. aegypti* mosquitoes [79, 96]. One of these studies also reported vertical transmission of MAYV [79]. This represents another mechanism that may lead to maintenance of the virus in urban mosquito populations. Although *Ae. aegypti* mosquitoes have not been conclusively implicated as MAYV vectors, the isolation of MAYV from wild-caught *Ae. aegypti* mosquitoes combined with the evidence of vector competence in laboratory settings [22-24, 113] suggests that MAYV could spill over into an urban cycle. This hypothesis requires further study to explore natural MAYV infection in city-dwelling mosquitoes and additional controlled vector competence studies.

Our systematic review revealed substantial heterogeneity across included studies, even within animal orders. Heterogeneity may complicate the interpretation of pooled seroprevalence estimates [38]. An additional limitation involves the validity of serological assays used to detect MAYV infection in animals. While plaque reduction NT is considered the “gold standard test” for detecting neutralizing antibodies to MAYV, some of the studies in the review instead relied on the less-specific HI test for antibody detection [101]. Furthermore, antibodies to other alphaviruses in the Semliki Forest serocomplex (e.g., CHIKV) may cross-react in serological tests [116]. Therefore, interpretation of seroprevalence estimates should be done with caution especially in the absence of confirmatory NT. Finally, unpublished data and articles with low quality scores were included in this review due to the paucity of eligible studies. Therefore, readers should consider the heterogeneity of study quality when interpreting the results of pooled seroprevalence estimates.

### Conclusions

MAYV is an emerging arbovirus that poses a major threat to human populations in Latin America. In order for public health authorities to effectively design MAYV surveillance and control programs, an understanding of the disease ecology is essential. This systematic review adds to existing knowledge regarding the potential animal reservoirs and arthropod vectors that are involved in the MAYV transmission cycle. These baseline data and maps of MAYV occurrence can direct risk emergence modeling and prediction efforts. Future studies involving experimental infection of primates and other non-human vertebrates are necessary to determine the animal species that may serve as amplifying hosts. Furthermore, additional experimental transmission studies may provide critical information regarding the potential for *Ae. aegypti* to facilitate urban spread of MAYV.

## Acknowledgments

We would like to thank Dr. Mauro Ramos for his assistance with reviewing Portuguese language articles. ELE is a Scientific researcher of the Consejo de Investigaciones Científicas y Tecnológicas (CONICET) from Argentina.

## FUNDING STATEMENT

This work was in part conducted by the Infectious Disease Clinical Research Program (IDCRP), a Department of Defense (DoD) program executed by the Uniformed Services University of the Health Sciences (USU) through a cooperative agreement with The Henry M. Jackson Foundation for the Advancement of Military Medicine, Inc. (HJF). This project has been supported with federal funds from the National Institute of Allergy and Infectious Diseases, National Institutes of Health (NIH), under Inter Agency Agreement Y1-Al-5072 and from the Defense Health Program, U.S. Department of Defense, under award HU0001190002.

A.A.’s participation in this work was supported by funding from Armed Forces Health Surveillance Branch - Global Emerging Infections Surveillance (GEIS) Project #P0044_20_NS and NASA Applied Sciences Program – Health and Air Quality, Grant #17-HAQ17-0065.

## DISCLAIMER

The contents of this publication are the sole responsibility of the author(s) and do not necessarily reflect the views, opinions or policies of Uniformed Services University of the Health Sciences (USUHS), the Department of Defense (DoD), the Departments of the Army, Navy, or Air Force. Mention of trade names, commercial products, or organizations does not imply endorsement by the U.S. Government.

## CONFLICT OF INTEREST

The authors declare no conflicts of interest.

## Supplementary Materials

**S1 Table. PRISMA Checklist**

**S2 Table. MAYV positivity by taxa of wild mammals in included studies S3 Table. MAYV positivity by taxa of wild birds in included studies**

**S4 Table. MAYV positivity in domestic or sentinel animals studied**

**S5 Table. Pooled Prevalence Table (Random effects using GLMM with logit transformation)**

**S6 Table. Primate Genera Pooled Prevalence Table (Random effects with Freeman-Tukey double arcsine transformation)**

**S7 Table. Primate Genera Pooled Prevalence Table (Random effects using GLMM with logit transformation)**

**S8 Table. Pooled Prevalence Table (Fixed effects with Freeman-Tukey double arcsine transformation)**

**S9 Table. Pooled Prevalence Table (Fixed effects using GLMM with logit transformation) S1 Fig. Funnel plots for estimates of MAYV seroprevalence in non-human animal reservoirs**

**S2 Fig: Funnel plots for estimates of MAYV seroprevalence in non-human primate genera**

**S10 Table. Complete arthropod results by genus**

**S11 Table. Egger’s test for publication bias**

## Notes

### Competing Interest Statement

The authors have declared no competing interest.

## References

1. Anderson CR, Downs WG, Wattley GH, Ahin NW, Reese AA. Mayaro virus: a new human disease agent. II. Isolation from blood of patients in Trinidad, B.W.I. Am J Trop Med Hyg. 1957;6(6):1012–6. Epub 1957/11/01. doi: 10.4269/ajtmh.1957.6.1012. PubMed PMID: 13487973.

2. Suhrbier A, Jaffar-Bandjee MC, Gasque P. Arthritogenic alphaviruses--an overview. Nat Rev Rheumatol. 2012;8(7):420–9. Epub 2012/05/09. doi: 10.1038/nrrheum.2012.64. PubMed PMID: 22565316.

3. Pan American Health Organization/World Health Organization. Epidemiological Alert: Mayaro Fever. Washington, D.C.: PAHO/WHO: 2019 May 1, 2019. Report No.

4. Causey OR, Maroja OM. Mayaro virus: a new human disease agent. III. Investigation of an epidemic of acute febrile illness on the river Guama in Para, Brazil, and isolation of Mayaro virus as causative agent. Am J Trop Med Hyg. 1957;6(6):1017–23. Epub 1957/11/01. PubMed PMID: 13487974.

5. LeDuc JW, Pinheiro FP, Travassos da Rosa AP. An outbreak of Mayaro virus disease in Belterra, Brazil. II. Epidemiology. Am J Trop Med Hyg. 1981;30(3):682–8. Epub 1981/05/01. doi: 10.4269/ajtmh.1981.30.682. PubMed PMID: 6266264.

6. Schaeffer M, Gajdusek DC, Lema AB, Eichenwald H. Epidemic jungle fevers among Okinawan colonists in the Bolivian rain forest. I. Epidemiology. Am J Trop Med Hyg. 1959;8(3):372–96. doi: 10.4269/ajtmh.1959.8.372.

7. Auguste AJ, Liria J, Forrester NL, Giambalvo D, Moncada M, Long KC, et al. Evolutionary and Ecological Characterization of Mayaro Virus Strains Isolated during an Outbreak, Venezuela, 2010. Emerg Infect Dis. 2015;21(10):1742–50. Epub 2015/09/25. doi: 10.3201/eid2110.141660. PubMed PMID: 26401714; PubMed Central PMCID: PMCPMC4593426.

8. Forshey BM, Guevara C, Laguna-Torres VA, Cespedes M, Vargas J, Gianella A, et al. Arboviral etiologies of acute febrile illnesses in Western South America, 2000-2007. PLoS Negl Trop Dis. 2010;4(8):e787. Epub 2010/08/14. doi: 10.1371/journal.pntd.0000787. PubMed PMID: 20706628; PubMed Central PMCID: PMCPMC2919378.

9. Jonkers AH, Spence L, Karbaat J. Arbovirus infections in Dutch military personnel stationed in Surinam. Further studies. Trop Geogr Med. 1968;20(3):251–6. Epub 1968/09/01. PubMed PMID: 5683357.

10. Navarrete-Espinosa J, Gomez-Dantes H. Arbovirus causales de fiebre hemorrágica en pacientes del Instituto Mexicano del Seguro Social. Rev Med Inst Mex Seguro Soc. 2006;44(4):347–53. Epub 2006/08/15. PubMed PMID: 16904038.

11. Groot H. Estudios sobre virus transmitidos por artropodos en Colombia. Rev Acad Colomb Cienc. 1964;12(46):191–217. doi: 10.18257/raccefyn.565.

12. Talarmin A, Chandler LJ, Kazanji M, de Thoisy B, Debon P, Lelarge J, et al. Mayaro virus fever in French Guiana: isolation, identification, and seroprevalence. Am J Trop Med Hyg. 1998;59(3):452–6. Epub 1998/09/28. doi: 10.4269/ajtmh.1998.59.452. PubMed PMID: 9749643.

13. Blohm G, Elbadry MA, Mavian C, Stephenson C, Loeb J, White S, et al. Mayaro as a Caribbean traveler: Evidence for multiple introductions and transmission of the virus into Haiti. Int J Infect Dis. 2019;87:151–3. Epub 2019/08/06. doi: 10.1016/j.ijid.2019.07.031. PubMed PMID: 31382049.

14. Izurieta RO, Macaluso M, Watts DM, Tesh RB, Guerra B, Cruz LM, et al. Hunting in the Rainforest and Mayaro Virus Infection: An emerging Alphavirus in Ecuador. J Glob Infect Dis. 2011;3(4):317–23. Epub 2012/01/10. doi: 10.4103/0974-777x.91049. PubMed PMID: 22223990; PubMed Central PMCID: PMCPMC3249982.

15. Plowright RK, Parrish CR, McCallum H, Hudson PJ, Ko AI, Graham AL, et al. Pathways to zoonotic spillover. Nat Rev Microbiol. 2017;15(8):502–10. Epub 2017/05/31. doi: 10.1038/nrmicro.2017.45. PubMed PMID: 28555073; PubMed Central PMCID: PMCPMC5791534.

16. Viana M, Mancy R, Biek R, Cleaveland S, Cross PC, Lloyd-Smith JO, et al. Assembling evidence for identifying reservoirs of infection. Trends Ecol Evol. 2014;29(5):270–9. Epub 2014/04/15. doi: 10.1016/j.tree.2014.03.002. PubMed PMID: 24726345; PubMed Central PMCID: PMCPMC4007595.

17. Haydon DT, Cleaveland S, Taylor LH, Laurenson MK. Identifying reservoirs of infection: a conceptual and practical challenge. Emerg Infect Dis. 2002;8(12):1468–73. Epub 2002/12/25. doi: 10.3201/eid0812.010317. PubMed PMID: 12498665; PubMed Central PMCID: PMCPMC2738515.

18. Pezzi L, Reusken CB, Weaver SC, Drexler JF, Busch M, LaBeaud AD, et al. GloPID-R report on Chikungunya, O’nyong-nyong and Mayaro virus, part I: Biological diagnostics. Antiviral Res. 2019;166:66–81. Epub 2019/03/25. doi: 10.1016/j.antiviral.2019.03.009. PubMed PMID: 30905821.

19. Hoch AL, Peterson NE, LeDuc JW, Pinheiro FP. An outbreak of Mayaro virus disease in Belterra, Brazil. III. Entomological and ecological studies. Am J Trop Med Hyg. 1981;30(3):689–98. Epub 1981/05/01. doi: 10.4269/ajtmh.1981.30.689. PubMed PMID: 6266265.

20. Seymour C, Peralta PH, Montgomery GG. Serologic evidence of natural togavirus infections in Panamanian sloths and other vertebrates. Am J Trop Med Hyg. 1983;32(4):854–61. Epub 1983/07/01. doi: 10.4269/ajtmh.1983.32.854. PubMed PMID: 6309027.

21. de Thoisy B, Gardon J, Salas RA, Morvan J, Kazanji M. Mayaro virus in wild mammals, French Guiana. Emerg Infect Dis. 2003;9(10):1326–9. Epub 2003/11/12. doi: 10.3201/eid0910.030161. PubMed PMID: 14609474; PubMed Central PMCID: PMCPMC3033094.

22. Long KC, Ziegler SA, Thangamani S, Hausser NL, Kochel TJ, Higgs S, et al. Experimental transmission of Mayaro virus by Aedes aegypti. Am J Trop Med Hyg. 2011;85(4):750–7. Epub 2011/10/07. doi: 10.4269/ajtmh.2011.11-0359. PubMed PMID: 21976583; PubMed Central PMCID: PMCPMC3183788.

23. Wiggins K, Eastmond B, Alto BW. Transmission potential of Mayaro virus in Florida Aedes aegypti and Aedes albopictus mosquitoes. Med Vet Entomol. 2018;32(4):436–42. Epub 2018/07/15. doi: 10.1111/mve.12322. PubMed PMID: 30006976.

24. Brustolin M, Pujhari S, Henderson C, Rasgon J. Emergent viruses and their interactions in Aedes aegypti: Mayaro and zika virus coinfected mosquitoes can successfully transmit both pathogens. Am J Trop Med Hyg. 2019;101(5):50. doi: 10.4269/ajtmh.abstract2019.

25. Mourao MP, Bastos Mde S, de Figueiredo RP, Gimaque JB, Galusso Edos S, Kramer VM, et al. Mayaro fever in the city of Manaus, Brazil, 2007-2008. Vector Borne Zoonotic Dis. 2012;12(1):42–6. Epub 2011/09/20. doi: 10.1089/vbz.2011.0669. PubMed PMID: 21923266; PubMed Central PMCID: PMCPMC3249893.

26. Page MJ, McKenzie JE, Bossuyt PM, Boutron I, Hoffmann TC, Mulrow CD, et al. The PRISMA 2020 statement: An updated guideline for reporting systematic reviews. Int J Surg. 2021;88:105906. Epub 2021/04/02. doi: 10.1016/j.ijsu.2021.105906. PubMed PMID: 33789826.

27. Clark K, Karsch-Mizrachi I, Lipman DJ, Ostell J, Sayers EW. GenBank. Nucleic Acids Res. 2016;44(D1):D67–D72. Epub 2015/11/20. doi: 10.1093/nar/gkv1276. PubMed PMID: 26590407.

28. Ding H, Gao YM, Deng Y, Lamberton PH, Lu DB. A systematic review and meta-analysis of the seroprevalence of Toxoplasma gondii in cats in mainland China. Parasit Vectors. 2017;10(1):27. Epub 2017/01/15. doi: 10.1186/s13071-017-1970-6. PubMed PMID: 28086987; PubMed Central PMCID: PMCPMC5237326.

29. Rodríguez-Monguí E, Cantillo-Barraza O, Prieto-Alvarado FE, Cucunubá ZM. Heterogeneity of Trypanosoma cruzi infection rates in vectors and animal reservoirs in Colombia: a systematic review and meta-analysis. Parasit Vectors. 2019;12(1):308. Epub 2019/06/22. doi: 10.1186/s13071-019-3541-5. PubMed PMID: 31221188; PubMed Central PMCID: PMCPMC6585012.

30. Guernier V, Goarant C, Benschop J, Lau CL. A systematic review of human and animal leptospirosis in the Pacific Islands reveals pathogen and reservoir diversity. PLoS Negl Trop Dis. 2018;12(5):e0006503. Epub 2018/05/15. doi: 10.1371/journal.pntd.0006503. PubMed PMID: 29758037; PubMed Central PMCID: PMCPMC5967813.

31. ESRI. ArcGIS Desktop: Release 10. Redlands, CA: Environmental Systems Research Institute.; 2011.

32. Acosta-Ampudia Y, Monsalve DM, Rodriguez Y, Pacheco Y, Anaya JM, Ramirez-Santana C. Mayaro: an emerging viral threat? Emerg Microbes Infect. 2018;7(1):163. Epub 2018/09/27. doi: 10.1038/s41426-018-0163-5. PubMed PMID: 30254258; PubMed Central PMCID: PMCPMC6156602.

33. Haidich AB. Meta-analysis in medical research. Hippokratia. 2010;14(Suppl 1):29–37. Epub 2011/04/14. PubMed PMID: 21487488; PubMed Central PMCID: PMCPMC3049418.

34. 34. Higgins JPT, Thomas J, Chandler J, Cumpston M, Li T, Page MJ, et al. Cochrane Handbook for Systematic Reviews of Interventions version 6.0 Cochrane; 2019. Available from: www.training.cochrane.org/handbook.

35. Barendregt JJ, Doi SA, Lee YY, Norman RE, Vos T. Meta-analysis of prevalence. J Epidemiol Community Health. 2013;67(11):974–8. Epub 2013//08/22. doi: 10.1136/jech-2013-203104. PubMed PMID: 23963506.

36. Schwarzer G, Chemaitelly H, Abu-Raddad LJ, Rücker G. Seriously misleading results using inverse of Freeman-Tukey double arcsine transformation in meta-analysis of single proportions. Res Synth Methods. 2019;10(3):476–83. Epub 2019/04/05. doi: 10.1002/jrsm.1348. PubMed PMID: 30945438; PubMed Central PMCID: PMCPMC6767151.

37. Warton DI, Hui FK. The arcsine is asinine: the analysis of proportions in ecology. Ecology. 2011;92(1):3–10. Epub 2011/05/13. doi: 10.1890/10-0340.1. PubMed PMID: 21560670.

38. Higgins JPT, Thompson SG, Deeks JJ, Altman DG. Measuring inconsistency in meta-analyses BMJ. 2003;327:557–60.

39. Quintana DS. From pre-registration to publication: a non-technical primer for conducting a meta-analysis to synthesize correlational data. Front Psychol. 2015;6(1549). doi: 10.3389/fpsyg.2015.01549.

40. Balduzzi S, Rücker G, Schwarzer G. How to perform a meta-analysis with R: a practical tutorial. Evid Based Ment Health. 2019;22(4):153–60. Epub 2019/09/30. doi: 10.1136/ebmental-2019-300117. PubMed PMID: 31563865.

41. Egger M, Davey Smith G, Schneider M, Minder C. Bias in meta-analysis detected by a simple, graphical test. BMJ. 1997;315(7109):629–34. Epub 1997/10/06. doi: 10.1136/bmj.315.7109.629. PubMed PMID: 9310563; PubMed Central PMCID: PMCPMC2127453.

42. Duval S, Tweedie R. Trim and fill: A simple funnel-plot-based method of testing and adjusting for publication bias in meta-analysis. Biometrics. 2000;56(2):455–63. Epub 2000/07/06. doi: 10.1111/j.0006-341x.2000.00455.x. PubMed PMID: 10877304.

43. Messina JP, Brady OJ, Pigott DM, Brownstein JS, Hoen AG, Hay SI. A global compendium of human dengue virus occurrence. Sci Data. 2014;1:140004. Epub 2014/01/01. doi: 10.1038/sdata.2014.4. PubMed PMID: 25977762; PubMed Central PMCID: PMCPMC4322574.

44. Pigott DM, Golding N, Messina JP, Battle KE, Duda KA, Balard Y, et al. Global database of leishmaniasis occurrence locations, 1960-2012. Sci Data. 2014;1:140036. Epub 2014/01/01. doi: 10.1038/sdata.2014.36. PubMed PMID: 25984344; PubMed Central PMCID: PMCPMC4432653.

45. Runfola D, Anderson A, Baier H, Crittenden M, Dowker E, Fuhrig S, et al. geoBoundaries: A global database of political administrative boundaries. PloS One. 2020;15(4):e0231866. Epub 2020/04/25. doi: 10.1371/journal.pone.0231866. PubMed PMID: 32330167; PubMed Central PMCID: PMCPMC7182183 Allen Hamilton, and Deloitte, respectively. This does not alter our adherence to PLOS ONE policies on sharing data and materials.

46. de Thoisy B, Vogel I, Reynes JM, Pouliquen JF, Carme B, Kazanji M, et al. Health evaluation of translocated free-ranging primates in French Guiana. Am J Primatol. 2001;54(1):1–16. Epub 2001/05/01. doi: 10.1002/ajp.1008. PubMed PMID: 11329164.

47. Aitken TH, Downs WG, Anderson CR, Spence L, Casals J. Mayaro virus isolated from a Trinidadian mosquito, Mansonia venezuelensis. Science (New York, NY). 1960;131(3405):986. Epub 1960/04/01. doi: 10.1126/science.131.3405.986. PubMed PMID: 13792204.

48. Aitken TH, Spence L, Jonkers AH, Downs WG. A 10-year survey of Trinidadian arthropods for natural virus infections (1953-1963). J Med Entomol. 1969;6(2):207–15. Epub 1969/05/01. doi: 10.1093/jmedent/6.2.207. PubMed PMID: 5807863.

49. Batista PM, Andreotti R, Almeida PS, Marques AC, Rodrigues SG, Chiang JO, et al. Detection of arboviruses of public health interest in free-living New World primates (Sapajus spp.; Alouatta caraya) captured in Mato Grosso do Sul, Brazil. Rev Soc Bras Med Trop. 2013;46(6):684–90. Epub 2014/01/30. doi: 10.1590/0037-8682-0181-2013. PubMed PMID: 24474008.

50. Paulo M, Renato A, Da Carneiro Rocha T, Eliane C, Navarro da Silva M. Serosurvey of arbovirus in free-living non-human primates (Sapajus spp.) in Brazil. J Environ Anal Chem. 2015;2(155):2380–91.1000155.

51. Woodall JP. Virus Research in Amazonia. Atas do Simpósio Sobre a Biota Amazônica; Para, Brazil1967. p. 31–63.

52. Taylor RM. Catalogue of arthropod-borne viruses of the world: a collection of data on registered arthropod-borne animal viruses: US Public Health Service; 1967.

53. Araujo FAA, Wada MY, da Silva EV, Cavalcante GC, Magalhaes VS, de Andrade Filho GV, et al. Primeiro inquérito sorológico em aves migratórias e nativas do Parque Nacional da Lagoa do Peixe/RS para detecção do vírus do Nilo Ocidental. In: Ministério da Saúde Secretaria de Vigilância em Saúde, editor. Boletim Eletrônico Epidemiologico, 2003.

54. Araújo FAA, Vianna RdST, Andrade Filho GVd, Melhado DL, Todeschini B, Cavalcante e Cavalcanti G, et al. Segundo inquérito sorológico em aves migratórias e residentes do parque nacional da Lagoa do Peixe/RS para detecção do vírus da Febre da Febre do Nilo Ocidental e outros vírus. In: Ministério da Saúde Secretaria de Vigilância em Saúde, editor. Boletim Eletrônico Epidemiologico, 2004.

55. Araújo FAA, Vianna RdST, Wada MY, Silva ÉVd, Doretto L, Cavalcante GCe, et al. Inquérito sorológico em aves migratórias e residentes de Galinhos/RN para detecção do vírus da Febre do Nilo Ocidental e outros vírus. In: Ministério da Saúde Secretaria de Vigilância em Saúde, editor. Boletim Eletrônico Epidemiológico, 2004.

56. Araujo FAA, Lima PC, Andrade MA, de Sá Jayme V, Ramos DG, Da Silveira SL. Soroprevalência de anticorpos “anti-arbovírus” de importância em saúde pública em aves selvagens, Brasil–2007 e 2008. Ciênc Anim Brasil. 2012;13(1):115–23. doi: 10.5216/cab.v13i1.16834.

57. Araujo FAA, Andrade MA, Jayme VS, Santos AL, Roman APM, Ramos DG, et al. Anticorpos antialfavírus detectados em equinos durante diferentes epizootias de encefalite equina, Paraíba, 2009. Rev Bras Ciênc Vet. 2012;19(1):80–5. doi: 10.4322/rbcv.2014.086.

58. Azevedo RS, Silva EV, Carvalho VL, Rodrigues SG, Neto JPN, Monteiro HA, et al. Mayaro fever virus, Brazilian amazon. Emerg Infect Dis. 2009;15(11):1830. doi: 10.3201/eid1511.090461.

59. Batista PM, Andreotti R, Chiang JO, Ferreira MS, Vasconcelos PF. Seroepidemiological monitoring in sentinel animals and vectors as part of arbovirus surveillance in the state of Mato Grosso do Sul, Brazil. Rev Soc Bras Med Trop. 2012;45(2):168–73. Epub 2012/04/27. doi: 10.1590/s0037-86822012000200006. PubMed PMID: 22534986.

60. Calisher CH, Gutierrez E, Maness KS, Lord RD. Isolation of Mayaro virus from a migrating bird captured in Louisiana in 1967. Bull Pan Am Health Organ. 1974;8(3):243–8. Epub 1974/01/01. PubMed PMID: 4418030.

61. Carrera JP, Cucunubá ZM, Neira K, Lambert B, Pittí Y, Liscano J, et al. Endemic and Epidemic Human Alphavirus Infections in Eastern Panama: An Analysis of Population-Based Cross-Sectional Surveys. Am J Trop Med Hyg. 2020. Epub 2020/10/31. doi: 10.4269/ajtmh.20-0408. PubMed PMID: 33124532.

62. Casseb AdR. Soroprevalência de anticorpos e padronização do teste ELISA sanduíche indireto para 19 tipos de arbovírus em herbívoros domésticos [Ph.D. Thesis]. Belém: Universidade Federal do Pará; 2010. Available from: http://repositorio.ufpa.br/jspui/handle/2011/4760.

63. Casseb AdR, Brito TC, Silva MRMd, Chiang JO, Martins LC, Silva SPd, et al. Prevalence of antibodies to equine alphaviruses in the State of Pará, Brazil. Arq Inst Biol. 2016;83. doi: 10.1590/1808-1657000202014.

64. Catenacci LS. Abordagem one health para vigilância de arbovirus na Mata Atlântica do sul da Bahia, Brasil. [Ph.D. Thesis]. Ananindeua: Instituto Evandro Chagas; 2017. Available from: https://patua.iec.gov.br/handle/iec/3073.

65. Cruz ACR, Prazeres AdSCd, Gama EC, Lima MFd, Azevedo RdSS, Casseb LMN, et al. Vigilância sorológica para arbovírus em Juruti, Pará, Brasil. Cadernos de saude publica. 2009;25(11):2517–23.

66. Degallier N, Travassos da Rosa AP, Vasconcelos PFC, Hervé JP, Sa Filho GC, Travassos da Rosa JFS, et al. Modifications of arbovirus transmission in relation to construction of dams in Brazilian Amazonia Journal of the Brazilian Association for the Advancement of Science. 1992;44.

67. Diaz LA, Diaz Mdel P, Almiron WR, Contigiani MS. Infection by UNA virus (Alphavirus; Togaviridae) and risk factor analysis in black howler monkeys (Alouatta caraya) from Paraguay and Argentina. Trans R Soc Trop Med Hyg. 2007;101(10):1039–41. Epub 2007/07/31. doi: 10.1016/j.trstmh.2007.04.009. PubMed PMID: 17658571.

68. Esposito DL, da Fonseca BA. Complete Genome Sequence of Mayaro Virus (Togaviridae, Alphavirus) Strain BeAr 20290 from Brazil. Genome Announc. 2015;3(6). Epub 2015/12/19. doi: 10.1128/genomeA.01372-15. PubMed PMID: 26679574; PubMed Central PMCID: PMCPMC4683219.

69. da Silva Ferreira R, de Toni Aquino da Cruz LC, Souza VJ, da Silva Neves NA, de Souza VC, Filho LCF, et al. Insect-specific viruses and arboviruses in adult male culicids from Midwestern Brazil. Infect Genet Evol. 2020:104561. Epub 2020/09/23. doi: 10.1016/j.meegid.2020.104561. PubMed PMID: 32961364.

70. Galindo P, Srihongse S, De Rodaniche E, Grayson MA. An ecological survey for arboviruses in Almirante, Panama, 1959-1962. Am J Trop Med Hyg. 1966;15(3):385–400. Epub 1966/05/01. doi: 10.4269/ajtmh.1966.15.385. PubMed PMID: 4380043.

71. Galindo P, Srihongse S. Transmission of arboviruses to hamsters by the bite of naturally infected Culex (Melanoconion) mosquitoes. Am J Trop Med Hyg. 1967;16(4):525–30. Epub 1967/07/01. doi: 10.4269/ajtmh.1967.16.525. PubMed PMID: 4952151.

72. Galindo P, Adames A, Peralta P, Johnson C, Read R. Impacto de la hidroeléctrica de Bayano en la transmisión de arbovirus. Rev Med Pan. 1983;8:89–134.

73. Gibrail MM. Detecção de anticorpos para arbovirus em primatas não humanos no município de Goiânia, Goiás [M.Sc. Thesis]. Goiânia: Universidade Federal de Goiás; 2015. Available from: https://repositorio.bc.ufg.br/tede/handle/tede/5552.

74. Gomes FA, Jansen AM, Machado RZ, Jesus Pena HF, Fumagalli MJ, Silva A, et al. Serological evidence of arboviruses and coccidia infecting horses in the Amazonian region of Brazil. PloS One. 2019;14(12):e0225895. Epub 2019/12/13. doi: 10.1371/journal.pone.0225895. PubMed PMID: 31830142.

75. Groot H, Morales A, Vidales H. Virus isolations from forest mosquitoes in San Vicente de Chucuri, Colombia. Am J Trop Med Hyg. 1961;10:397–402. Epub 1961/05/01. doi: 10.4269/ajtmh.1961.10.397. PubMed PMID: 13708940.

76. Henriques DA. Caracterização molecular de arbovírus isolados da fauna diptera nematocera do Estado de Rondônia (Amazônia ocidental brasileira) [Ph.D. Thesis]. São Paulo: Universidade de São Paulo; 2008. Available from: https://teses.usp.br/teses/disponiveis/42/42132/tde-27032009-124003/pt-br.php.

77. Kubiszeski JR. Arboviroses emergentes no município de Sinop-MT: pesquisa de vetores [Ph.D. Thesis]. Sinop: Universidade Federal de Mato Grosso; 2016. Available from: https://teses.usp.br/teses/disponiveis/42/42132/tde-27032009-124003/pt-br.php.

78. Laroque PO, Valença-Montenegro MM, Ferreira DRA, Chiang JO, Cordeiro MT, Vasconcelos PFC, et al. Levantamento soroepidemiológico para arbovírus em macaco-prego-galego (Cebus flavius) de vida livre no estado da Paraíba e em macaco-prego (Cebus libidinosus) de cativeiro do nordeste do Brasil. Pesq Vet Bras. 2014;34:462–8.

79. Maia LMS, Bezerra MCF, Costa MCS, Souza EM, Oliveira MEB, Ribeiro ALM, et al. Natural vertical infection by dengue virus serotype 4, Zika virus and Mayaro virus in Aedes (Stegomyia) aegypti and Aedes (Stegomyia) albopictus. Med Vet Entomol. 2019;33(3):437–42. Epub 2019/02/19. doi: 10.1111/mve.12369. PubMed PMID: 30776139.

80. Martinez D, Hernandez C, Munoz M, Armesto Y, Cuervo A, Ramirez JD. Identification of Aedes (Diptera: Culicidae) Species and Arboviruses Circulating in Arauca, Eastern Colombia. Front Ecol Evol. 2020;8. doi: 10.3389/fevo.2020.602190. PubMed PMID: WOS:000596835300001.

81. Medlin S, Deardorff ER, Hanley CS, Vergneau-Grosset C, Siudak-Campfield A, Dallwig R, et al. Serosurvey of Selected Arboviral Pathogens in Free-Ranging, Two-Toed Sloths (Choloepus Hoffmanni) and Three-Toed Sloths (Bradypus Variegatus) In Costa Rica, 2005-07. J Wildl Dis. 2016;52(4):883–92. Epub 2016/08/02. doi: 10.7589/2015-02-040. PubMed PMID: 27479900; PubMed Central PMCID: PMCPMC5189659.

82. Medina G, Garzaro DJ, Barrios M, Auguste AJ, Weaver SC, Pujol FH. Genetic diversity of Venezuelan alphaviruses and circulation of a Venezuelan equine encephalitis virus subtype IAB strain during an interepizootic period. Am J Trop Med Hyg. 2015;93(1):7–10. Epub 2015/05/06. doi: 10.4269/ajtmh.14-0543. PubMed PMID: 25940191; PubMed Central PMCID: PMCPMC4497907.

83. Moreira-Soto A, Carneiro ID, Fischer C, Feldmann M, Kummerer BM, Silva NS, et al. Limited Evidence for Infection of Urban and Peri-urban Nonhuman Primates with Zika and Chikungunya Viruses in Brazil. mSphere. 2018;3(1). doi: 10.1128/mSphere.00523-17. PubMed PMID: WOS:000425277500024.

84. Nunes MR, Barbosa TF, Casseb LM, Nunes Neto JP, Segura Nde O, Monteiro HA, et al. Eco-epidemiologia dos arbovirus na area de influencia da rodovia Cuiaba-Santarem (BR 163), Estado do Para, Brasil. Cad Saude Publica. 2009;25(12):2583–602. Epub 2010/03/02. doi: 10.1590/s0102-311x2009001200006. PubMed PMID: 20191150.

85. Pauvolid-Correa A, Tavares FN, Costa EV, Burlandy FM, Murta M, Pellegrin AO, et al. Serologic evidence of the recent circulation of Saint Louis encephalitis virus and high prevalence of equine encephalitis viruses in horses in the Nhecolandia sub-region in South Pantanal, Central-West Brazil. Mem Inst Oswaldo Cruz. 2010;105(6):829–33. Epub 2010/10/15. doi: 10.1590/s0074-02762010000600017. PubMed PMID: 20945001.

86. Pauvolid-Correa A, Juliano RS, Campos Z, Velez J, Nogueira RM, Komar N. Neutralising antibodies for Mayaro virus in Pantanal, Brazil. Mem Inst Oswaldo Cruz. 2015;110(1):125–33. Epub 2015/03/06. doi: 10.1590/0074-02760140383. PubMed PMID: 25742272; PubMed Central PMCID: PMCPMC4371226.

87. Pauvolid-Correa A. Estudo sobre arbovírus em populações de eqüinos e artrópodes na sub-região da Nhecolândia no Pantanal de Mato Grosso do Sul [M.Sc. Thesis]. Rio de Janeiro: Fundação Oswaldo Cruz; 2008. Available from: https://www.arca.fiocruz.br/handle/icict/21142.

88. Perez JG, Carrera JP, Serrano E, Pitti Y, Maguina JL, Mentaberre G, et al. Serologic Evidence of Zoonotic Alphaviruses in Humans from an Indigenous Community in the Peruvian Amazon. Am J Trop Med Hyg. 2019. Epub 2019/10/02. doi: 10.4269/ajtmh.18-0850. PubMed PMID: 31571566.

89. Pinheiro FP, Bensabath G, Andrade AH, Lins ZC, Fraihi H, Tang AT, et al. Infectious diseases along Brazil’s Trans-Amazon Highway: surveillance and research. Bull Pan Am Health Organ. 1974;8(111).

90. Pinheiro GG, Rocha MN, de Oliveira MA, Moreira LA, Andrade JD. Detection of Yellow Fever Virus in Sylvatic Mosquitoes during Disease Outbreaks of 2017-2018 in Minas Gerais State, Brazil. Insects. 2019;10(5). doi: 10.3390/insects10050136. PubMed PMID: WOS:000476846800018.

91. Powers AM, Aguilar PV, Chandler LJ, Brault AC, Meakins TA, Watts D, et al. Genetic relationships among Mayaro and Una viruses suggest distinct patterns of transmission. Am J Trop Med Hyg. 2006;75(3):461–9. Epub 2006/09/14. PubMed PMID: 16968922.

92. Price JL. Serological evidence of infection of Tacaribe virus and arboviruses in Trinidadian bats. Am J Trop Med Hyg. 1978;27(1 Pt 1):162–7. Epub 1978/01/01. doi: 10.4269/ajtmh.1978.27.162. PubMed PMID: 204207.

93. Ragan IK, Hartwig A, Bowen RA. Cold blood: Reptiles and amphibians as reservoir and over wintering hosts for arboviruses. Am J Trop Med Hyg. 2019;101(5):261. doi: 10.4269/ajtmh.abstract2019.

94. Sanmartín C, Mackenzie RB, Trapido H, Barreto P, Mullenax CH, Gutiérrez E, et al. Encefalitis equina venezolana en Colombia, 1967. Bol Oficina Sanit Panam. 1973;74(2):108-37. Epub 1973/02/01. PubMed PMID: 4265714.

95. Scherer WF, Madalengoitia J, Flores W, Acosta M. The first isolations of eastern encephalitis, group C, and Guama group arboviruses from the Peruvian Amazon region of western South America. Bull Pan Am Health Organ. 1975;9(1):19–26. Epub 1975/01/01. PubMed PMID: 238693.

96. Serra OP, Cardoso BF, Ribeiro AL, Santos FA, Slhessarenko RD. Mayaro virus and dengue virus 1 and 4 natural infection in culicids from Cuiaba, state of Mato Grosso, Brazil. Mem Inst Oswaldo Cruz. 2016;111(1):20–9. Epub 2016/01/20. doi: 10.1590/0074-02760150270. PubMed PMID: 26784852; PubMed Central PMCID: PMCPMC4727432.

97. Silva JWP. Aspectos ecológicos de vetores putativos do Vírus Mayaro e Vírus Oropuche em estratificação vertical e horizontal em ambientes florestais e antropizados em uma comunidade rural no Amazonas [M.Sc. Thesis]. Manaus, AM: Oswaldo Cruz Foundation, Instituto Leônidas and Maria Deane; 2017. Available from: https://www.arca.fiocruz.br/handle/icict/23337.

98. Srihongse S, Galindo P, Eldridge BF. A survey to assess potential human disease hazards along proposed sea level canal routes in Panama and Colombia. V. Arbovirus infection in non human vertebrates. Mil Med. 1974;139(6):449–53.

99. Tauro LB, Cardoso CW, Souza RL, Nascimento LC, Santos DRD, Campos GS, et al. A localized outbreak of Chikungunya virus in Salvador, Bahia, Brazil. Mem Inst Oswaldo Cruz. 2019;114:e180597. Epub 2019/03/08. doi: 10.1590/0074-02760180597. PubMed PMID: 30843962; PubMed Central PMCID: PMCPMC6396974.

100. Turell MJ, Gozalo AS, Guevara C, Schoeler GB, Carbajal F, Lopez-Sifuentes VM, et al. Lack of Evidence of Sylvatic Transmission of Dengue Viruses in the Amazon Rainforest Near Iquitos, Peru. Vector Borne Zoonotic Dis. 2019;19(9):685–9. Epub 2019/04/10. doi: 10.1089/vbz.2018.2408. PubMed PMID: 30964397; PubMed Central PMCID: PMCPMC6716187.

101. Valentine MJ, Murdock CC, Kelly PJ. Sylvatic cycles of arboviruses in non-human primates. Parasit Vectors. 2019;12(1):463. Epub 2019/10/04. doi: 10.1186/s13071-019-3732-0. PubMed PMID: 31578140; PubMed Central PMCID: PMCPMC6775655.

102. Cornet M, Saluzzo JF, Hervy JP, Digoutte JP, Germain M, Chauvancy MF. Dengue 2 au Sénégal oriental: une pousse épizootique en milieu selvatique; isolements du virus à partir de moustiques et d’un singe et considérations épidémiologiques. Cah Orstom Ser Ent Med Parasitol. 1984;22:313–23.

103. Diallo M, Thonnon J, Traore-Lamizana M, Fontenille D. Vectors of Chikungunya virus in Senegal: current data and transmission cycles. Am J Trop Med Hyg. 1999;60(2):281–6. Epub 1999/03/11. doi: 10.4269/ajtmh.1999.60.281. PubMed PMID: 10072152.

104. Dick GW, Kitchen SF, Haddow AJ. Zika virus. I. Isolations and serological specificity. Trans R Soc Trop Med Hyg. 1952;46(5):509–20. Epub 1952/09/01. doi: 10.1016/0035-9203(52)90042-4. PubMed PMID: 12995440.

105. Althouse BM, Guerbois M, Cummings DAT, Diop OM, Faye O, Faye A, et al. Role of monkeys in the sylvatic cycle of chikungunya virus in Senegal. Nat Commun. 2018;9(1):1046. Epub 2018/03/15. doi: 10.1038/s41467-018-03332-7. PubMed PMID: 29535306; PubMed Central PMCID: PMCPMC5849707.

106. Kading RC, Borland EM, Cranfield M, Powers AM. Prevalence of antibodies to alphaviruses and flaviviruses in free-ranging game animals and nonhuman primates in the greater Congo basin. J Wildl Dis. 2013;49(3):587–99. Epub 2013/06/20. doi: 10.7589/2012-08-212. PubMed PMID: 23778608.

107. Lundström JO, Lindström KM, Olsen B, Dufva R, Krakower DS. Prevalence of sindbis virus neutralizing antibodies among Swedish passerines indicates that thrushes are the main amplifying hosts. J Med Entomol. 2001;38(2):289–97. Epub 2001/04/12. doi: 10.1603/0022-2585-38.2.289. PubMed PMID: 11296837.

108. Stephenson EB, Peel AJ, Reid SA, Jansen CC, McCallum H. The non-human reservoirs of Ross River virus: a systematic review of the evidence. Parasit Vectors. 2018;11(1):188. Epub 2018/03/21. doi: 10.1186/s13071-018-2733-8. PubMed PMID: 29554936; PubMed Central PMCID: PMCPMC5859426.

109. Barba M, Fairbanks EL, Daly JM. Equine viral encephalitis: prevalence, impact, and management strategies. Vet Med (Auckl)). 2019;10:99–110. Epub 2019/09/10. doi: 10.2147/vmrr.S168227. PubMed PMID: 31497528; PubMed Central PMCID: PMCPMC6689664.

110. Kuno G, Mackenzie JS, Junglen S, Hubálek Z, Plyusnin A, Gubler DJ. Vertebrate reservoirs of arboviruses: myth, synonym of amplifier, or reality? Viruses. 2017;9(7):185.

111. Kuno G, Chang GJ. Biological transmission of arboviruses: reexamination of and new insights into components, mechanisms, and unique traits as well as their evolutionary trends. Clin Microbiol Rev. 2005;18(4):608–37. Epub 2005/10/15. doi: 10.1128/cmr.18.4.608-637.2005. PubMed PMID: 16223950; PubMed Central PMCID: PMCPMC1265912.

112. World Health Organization Scientific Group. Arthropod-borne and rodent-borne viral diseases. Geneva, Switzerland: World Health Organization, 1985.

113. Pereira TN, Carvalho FD, De Mendonça SF, Rocha MN, Moreira LA. Vector competence of Aedes aegypti, Aedes albopictus, and Culex quinquefasciatus mosquitoes for Mayaro virus. PLoS Negl Trop Dis. 2020;14(4):e0007518. Epub 2020/04/15. doi: 10.1371/journal.pntd.0007518. PubMed PMID: 32287269; PubMed Central PMCID: PMCPMC7182273.

114. Mackay IM, Arden KE. Mayaro virus: a forest virus primed for a trip to the city? Microbes Infect. 2016;18(12):724–34. Epub 2016/12/19. doi: 10.1016/j.micinf.2016.10.007. PubMed PMID: 27989728.

115. Figueiredo MLGd, Figueiredo LTM. Emerging alphaviruses in the Americas: Chikungunya and Mayaro. Revista da Sociedade Brasileira de Medicina Tropical. 2014;47(6):677–83. doi: 10.1590/0037-8682-0246-2014.

116. Hassing RJ, Leparc-Goffart I, Tolou H, van Doornum G, van Genderen PJ. Cross-reactivity of antibodies to viruses belonging to the Semliki forest serocomplex. Eurosurveillance. 2010;15(23).

