## Supplementary Material for "A systematic review and meta-analysis of the potential non-human animal reservoirs and arthropod vectors of the Mayaro virus"

**S1 Table. PRISMA Checklist**

| **Section and Topic** | **Item #** | **Checklist item** | **Location where item is reported** |
| --- | --- | --- | --- |
| **TITLE** | | |  |
| Title | 1 | Identify the report as a systematic review. | Page 1 |
| **ABSTRACT** | | |  |
| Abstract | 2 | See the PRISMA 2020 for Abstracts checklist. | Page 2 |
| **INTRODUCTION** | | |  |
| Rationale | 3 | Describe the rationale for the review in the context of existing knowledge. | Page 4-5 |
| Objectives | 4 | Provide an explicit statement of the objective(s) or question(s) the review addresses. | Page 5 |
| **METHODS** | | |  |
| Eligibility criteria | 5 | Specify the inclusion and exclusion criteria for the review and how studies were grouped for the syntheses. | Page 7 |
| Information sources | 6 | Specify all databases, registers, websites, organisations, reference lists and other sources searched or consulted to identify studies. Specify the date when each source was last searched or consulted. | Page 6 |
| Search strategy | 7 | Present the full search strategies for all databases, registers and websites, including any filters and limits used. | Page 6 |
| Selection process | 8 | Specify the methods used to decide whether a study met the inclusion criteria of the review, including how many reviewers screened each record and each report retrieved, whether they worked independently, and if applicable, details of automation tools used in the process. | Page 7 |
| Data collection process | 9 | Specify the methods used to collect data from reports, including how many reviewers collected data from each report, whether they worked independently, any processes for obtaining or confirming data from study investigators, and if applicable, details of automation tools used in the process. | Page 8 |
| Data items | 10a | List and define all outcomes for which data were sought. Specify whether all results that were compatible with each outcome domain in each study were sought (e.g. for all measures, time points, analyses), and if not, the methods used to decide which results to collect. | Page 7-8 |
|  | 10b | List and define all other variables for which data were sought (e.g. participant and intervention characteristics, funding sources). Describe any assumptions made about any missing or unclear information. | Page 7-8 |
| Study risk of bias assessment | 11 | Specify the methods used to assess risk of bias in the included studies, including details of the tool(s) used, how many reviewers assessed each study and whether they worked independently, and if applicable, details of automation tools used in the process. | Page 8-9 |
| Effect measures | 12 | Specify for each outcome the effect measure(s) (e.g. risk ratio, mean difference) used in the synthesis or presentation of results. | Page 10-11 |
| Synthesis methods | 13a | Describe the processes used to decide which studies were eligible for each synthesis (e.g. tabulating the study intervention characteristics and comparing against the planned groups for each synthesis (item #5)). | Page 10 |
|  | 13b | Describe any methods required to prepare the data for presentation or synthesis, such as handling of missing summary statistics, or data conversions. | Page 8 |
|  | 13c | Describe any methods used to tabulate or visually display results of individual studies and syntheses. | Page 8, 10 |
|  | 13d | Describe any methods used to synthesize results and provide a rationale for the choice(s). If meta-analysis was performed, describe the model(s), method(s) to identify the presence and extent of statistical heterogeneity, and software package(s) used. | Page 10-11 |
|  | 13e | Describe any methods used to explore possible causes of heterogeneity among study results (e.g. subgroup analysis, meta-regression). | NA |
|  | 13f | Describe any sensitivity analyses conducted to assess robustness of the synthesized results. | Page 10-11 |
| Reporting bias assessment | 14 | Describe any methods used to assess risk of bias due to missing results in a synthesis (arising from reporting biases). | Page 11 |
| Certainty assessment | 15 | Describe any methods used to assess certainty (or confidence) in the body of evidence for an outcome. | Page 11 |
| **RESULTS** | | |  |
| Study selection | 16a | Describe the results of the search and selection process, from the number of records identified in the search to the number of studies included in the review, ideally using a flow diagram. | Fig 1 |
|  | 16b | Cite studies that might appear to meet the inclusion criteria, but which were excluded, and explain why they were excluded. | Page 13 |
| Study characteristics | 17 | Cite each included study and present its characteristics. | Page 13-14 |
| Risk of bias in studies | 18 | Present assessments of risk of bias for each included study. | Table 2 |
| Results of individual studies | 19 | For all outcomes, present, for each study: (a) summary statistics for each group (where appropriate) and (b) an effect estimate and its precision n (e.g. confidence/credible interval), ideally using structured tables or plots. | Page 17-26, Table 7, Supplementary materials |
| Results of syntheses | 20a | For each synthesis, briefly summarise the characteristics and risk of bias among contributing studies. | Table 2 |
|  | 20b | Present results of all statistical syntheses conducted. If meta-analysis was done, present for each the summary estimate and its precision (e.g. confidence/credible interval) and measures of statistical heterogeneity. If comparing groups, describe the direction of the effect. | Table 7, Fig 3-4, Supplementary materials |
|  | 20c | Present results of all investigations of possible causes of heterogeneity among study results. | NA |
|  | 20d | Present results of all sensitivity analyses conducted to assess the robustness of the synthesized results. | Supplementary materials |
| Reporting biases | 21 | Present assessments of risk of bias due to missing results (arising from reporting biases) for each synthesis assessed. | Page 31, Supplementary materials |
| Certainty of evidence | 22 | Present assessments of certainty (or confidence) in the body of evidence for each outcome assessed. | NA |
| **DISCUSSION** | | |  |
| Discussion | 23a | Provide a general interpretation of the results in the context of other evidence. | Page 31-33 |
|  | 23b | Discuss any limitations of the evidence included in the review. | Page 35-36 |
|  | 23c | Discuss any limitations of the review processes used. | Page 35-36 |
|  | 23d | Discuss implications of the results for practice, policy, and future research. | Page 33-36 |
| **OTHER INFORMATION** | | |  |
| Registration and protocol | 24a | Provide registration information for the review, including register name and registration number, or state that the review was not registered. | Page 6 |
|  | 24b | Indicate where the review protocol can be accessed, or state that a protocol was not prepared. | NA |
|  | 24c | Describe and explain any amendments to information provided at registration or in the protocol. | NA |
| Support | 25 | Describe sources of financial or non-financial support for the review, and the role of the funders or sponsors in the review. | Page 36 |
| Competing interests | 26 | Declare any competing interests of review authors. | Page 37 |
| Availability of data, code and other materials | 27 | Report which of the following are publicly available and where they can be found: template data collection forms; data extracted from included studies; data used for all analyses; analytic code; any other materials used in the review. | Supplementary materials |

**S2 Table.** **MAYV positivity by taxa of wild mammals in included studies**

| **Family** | **Genus** | **Species** | **Common Name** | **MAYV Positive Samples** | **Total Positive** | **Total Tested^#^** | **Country of study*** |
| --- | --- | --- | --- | --- | --- | --- | --- |
| ***Order: Primate*** | | | | | | | |
| Aotidae | *Aotus* | *A. trivirgatus* | Three-striped night monkey | No | 0 | 6 | Panama [1, 2] |
|  |  | NA | NA | Yes | 1 | 4 | Colombia **[3]*** |
| Atelidae | *Alouatta* | *A. belzebul* | Red-handed howler | Yes | 1 | 1 | Brazil **[4]*** |
|  |  | *A. caraya* | Black Howler | Yes | 0 | 97 | Brazil [5, 6], Argentina/Paraguay [7] |
|  |  | *A. seniculus* | Venezuelan red howler | Yes | 52 | 99 | Peru **[8]***,  French Guiana **[9]*** |
|  |  | *A. villosa* | Guatemalan black howler | Yes | 3 | 5 | Panama **[1]*** [2] |
|  |  | NA | NA | Yes | 7 | 11 | Colombia **[3]*** |
|  | *Ateles* | *A. marginatus* | White-cheeked spider monkey | Yes | 1 | 1 | Brazil **[10]*** |
|  |  | NA | NA | No | 0 | 5 | Colombia [3] |
|  | *Lagothrix* | *L. poeppigii* | Silvery wooly monkey | Yes | 6 | 11 | Peru **[8]*** |
| Callithricidae | *Callithrix* | *C. jacchus* | Common marmoset | No | 0 | 1 | Brazil [10] |
|  |  | *C. argentata* | Silvery marmoset | Yes | 32 | 119 | Brazil **[4]*** |
|  |  | *C. penicillata* | Black-tufted marmoset | No | 0 | 3 | Brazil [6] |
|  | *Leontopithecus* | *L. chrysomelas* | Golden-headed lion tamarin | No | 0 | 103 | Brazil [11] |
|  | *Saguinas* | *S. oedipus* | Cotton-top tamarin | No | 0 | NA | Panama [2] |
|  |  | *S. geoffroyi* | Geoffrey’s tamarin | No | 0 | 32 | Panama [1] |
|  |  | *S. midas* | Red-handed tamarin | Yes | 8 | 42 | French Guiana **[9]*** |
| Cebidae | *Cebus* | *C. apella* | Tufted capuchin | Yes | 10 | 62 | Brazil **[5]*** |
|  |  | *C. capucinus* | Colombian white-faced capuchin | No | 0 | 1 | Panama [1, 2] |
|  |  | *C. albifrons* | White-fronted capuchin | No | 0 | 2 | Peru [8] |
|  |  | *C. libidinosus* | Black-striped capuchin | Yes | 6 | 142 | Brazil **[12]*** [6] |
|  |  | NA | NA | Yes | 4 | 13 | Colombia **[3]*** |
|  | *Sapajus* | *S. macrocephalus* | Large-headed capuchin | Yes | 1 | 6 | Peru **[8]*** |
|  |  | *S. flavius* | Blond capuchin | No | 0 | 32 | Brazil [10, 12] |
|  |  | *S. robustus* | Crested capuchin | No | 0 | 1 | Brazil [10] |
|  |  | *S. xanthosternos* | Golden-bellied capuchin | Yes | 1 | 2 | Brazil **[10]***[11] |
|  |  | NA | NA | Yes | 3 | 48 | Brazil **[13]*** [10] |
|  | *Saimiri* | *S. macrodon* | Ecuadorian squirrel monkey | No | 0 | 3 | Peru [8] |
|  |  | *S. sciureus* | Common squirrel monkey | Yes | 4 | 6 | French Guiana **[9]*** |
|  |  | NA | NA | Yes | 1 | 1 | Colombia **[3]*** |
| Pitheciidae | *Cacajao* | *C. calvus* | Bald uakari | Yes | 1 | 3 | Peru **[8]*** |
|  | *Callicebus* | *C. brunneus* | Brown titi | Yes | 1 | NA | Brazil **[14]*** |
|  |  | *C. donacophilus* | White-eared titi | No | 0 | 1 | Brazil [5] |
|  | *Pithecia* | *P. monachus* | Monk saki | No | 0 | 1 | Peru [8] |
|  |  | *P. pithecia* | White-faced saki | Yes | 4 | 5 | French Guiana **[9]*** |
| NA | NA | NA | NA | Yes | 7 | 53 | Brazil **[15]*[14]*[16]***, Panama/Colombia [17] |
| ***Order: Rodentia*** | | | | | | | |
| Cricetidae | *Neacomys* | *N. guianae* | Guianan neacomys | No | 0 | NA | Brazil [18] |
|  | *Nectomis* | *N. squamipes* | Atlantic Forest nectomys | No | 0 | NA | Brazil [18] |
|  | *Oryzomys* | *O. alfaroi* | Alfaro’s rice rat | No | 0 | 9 | Colombia [19] |
|  |  | *O. caliginosus* | Costa Rican dusky rice rat | No | 0 | 47 | Colombia [19] |
|  |  | *O. goeldi* | Large-headed rice rat | No | 0 | NA | Brazil [18] |
|  |  | *N/A* | NA | No | 0 | 458 | Panama/Colombia [17] |
|  | *Sigmodon* | *S. hispidus* | Hispid cotton rat | No | 0 | 183 | Panama [1, 20] |
|  | *Oxymycterus* | *O. amazonicus* | Amazonian hocicudo | No | 0 | NA | Brazil [18] |
|  | *Rhipidomys* | *R. latimanus* | Broad-footed climbing mouse | No | 0 | 7 | Colombia [19] |
|  | *Zygodontomys* | *Z. brevicauda* | Common cane mouse | No | 0 | 1 | Colombia [19] |
|  | NA | NA | NA | No | 0 | 150 | Brazil [4] |
| Cuniculidae | *Agouti* | *A. paca* | Lowland paca | Yes | 1 | 29 | Peru **[8]*,**  French Guiana [9]**,** Panama [1, 2], |
| Dasyproctidae | *Dasyprocta* | *D. fuliginosa* | Black agouti | Yes | 3 | 27 | Peru **[8]*** |
|  |  | *D. leporina* | Red-rumped agouti | Yes | 5 | 29 | French Guiana **[9]*** |
|  |  | *D. punctata* | Central American agouti | Yes | 3 | 5 | Panama **[1]*** [2] |
|  | *Myoprocta* | *M. acouchy* | Red acouchi | No | 0 | 29 | French Guiana [9] |
| Echimyidae | *Echimys* | NA | NA | Yes | 1 | 21 | French Guiana **[9]*** |
|  | *Proechimys* | *P. guyannensis* | Guyenne spiny rat | No | 0 | 2 | Brazil [18, 21] |
|  |  | *P. longicaudatus* | Long-tailed spiny rat | No | 0 | NA | Brazil [18] |
|  |  | *P. semispinosus* | Tome's spiny rat | No | 0 | 76 | Panama [1, 2, 20] |
|  |  | NA | NA | Yes | 1 | 149 | French Guiana **[9]*,**  Peru [8], Panama/Colombia [17], |
|  | NA | NA | N/A | No | 0 | 85 | Brazil [4] |
| Erethizontidae | *Coendou* | *C. melanurus* | Black-tailed hairy dwarf porcupine | Yes | 2 | 15 | French Guiana **[9]*** |
|  |  | *C. prehensilis* | Brazilian porcupine | Yes | 3 | 26 | French Guiana **[9]*** |
|  |  | *C. rothschildii* | Rothschild’s porcupine | No | 0 | 1 | Panama [1, 2] |
| Sciuridae | *Sciurus* | *S. granatensis* | Red-tailed squirrel | No | 0 | 11 | Panama [1, 2] |
|  |  | *S. igniventris* | Northern Amazon red squirrel | No | 0 | 1 | Peru [8] |
|  | NA | NA | NA | No | 0 | 2 | Brazil [4] |
| Muridae | *Mus* | *M. musculus* | House mouse | No | 0 | 2 | Colombia [19] |
|  | *Rattus* | *R. norvegicus* | Brown rat | No | 0 | 2 | Colombia [19] |
|  |  | *R. rattus* | Black rat | No | 0 | 40 | Brazil [18],  Colombia [19] |
|  |  | NA | NA | No | 0 | 2 | Colombia [19] |
| NA | NA | NA | NA | Yes | 71 | 1092 | Brazil **[15]*** [16] Panama/Colombia [17] |
| ***Order: Chiroptera*** | | | | | | | |
| Molossidae | *Molossus* | *M. ater* | Black mastiff bat | No | 0 | 165 | Trinidad [22] |
|  |  | *M. molossus* | Velvety free-tailed bat | No | 0 | 41 | Trinidad [22] |
|  | NA | NA | NA | No | 0 | 2 | Brazil [4] |
| Mormoopidae | *Pteronotus* | *P. davyi* | Davy’s naked-backed bat | No | 0 | 18 | Trinidad [22] |
|  |  | *P. parnellii* | Parnell’s mustached bat | No | 0 | 30 | Trinidad [22] |
| Natalidae | *Natalus* | *N. tumidirostris* | Trinidadian funnel-eared bat | No | 0 | 10 | Trinidad [22] |
| Phyllostomidae | *Anoura* | *A. geoffroyi* | Geoffroy’s tailless bat | No | 0 | 31 | Trinidad [22] |
|  | *Artibeus* | *A. cinereus* | Gervais’s fruit-eating bat | No | 0 | 22 | Trinidad [22] |
|  |  | *A. jamaicensis* | Jamaican fruit bat | No | 0 | 93 | Trinidad [22],  Panama [1] |
|  |  | *A. lituratus* | Great fruit-eating bat | No | 0 | 66 | Trinidad [22],  Panama [1],  Brazil [18],  Colombia [19] |
|  |  | *A. hartii* | Velvety fruit-eating bat | No | 0 | 1 | Colombia [19] |
|  | *Carollia* | *C. perspicillata* | Seba’s short-tailed bat | No | 0 | 192 | Trinidad [22],  Panama [1],  Brazil [18],  Colombia [19] |
|  | *Chiroderma* | *C. villosum* | Hairy big-eyed bat | No | 0 | 1 | Colombia [19] |
|  | *Glossophaga* | *G. soricina* | Pallas’s long-tongued bat | No | 0 | 47 | Trinidad [22],  Brazil [18],  Colombia [19] |
|  | *Phyllostomus* | *P. hastatus* | Greater spear-nosed bat | No | 0 | 170 | Trinidad [22] |
|  |  | *P. discolor* | Pale spear-nosed bat | No | 0 | N/A | Brazil [18] |
|  | *Sturnira* | NA | NA | No | 0 | 21 | Trinidad [22] |
|  |  | *S. lilium* | Little yellow-shouldered bat | No | 0 | 2 | Colombia [19] |
|  | *Vampyrops* | *V. helleri* | Heller’s broad-nosed bat | No | 0 | 30 | Trinidad [22] |
|  | NA | NA | NA | No | 0 | 172 | Brazil [4] |
| NA | NA | NA | NA | No | 0 | 431 | Panama [1], Panama/Colombia [17], Brazil [16] |
| ***Order: Cingulata*** | | | | | | | |
| Chlamyphoridae | *Cabassous* | *C. centralis* | Northern naked-tailed armadillo | No | 0 | 1 | Panama [1, 2] |
| Dasypodidae | *Dasypus* | *D. kappleri* | Greater long-nosed armadillo | No | 0 | 20 | French Guiana [9] |
|  |  | *D. novemcinctus* | Nine-banded armadillo | Yes | 6 | 48 | Peru **[8]***,  French Guiana **[9]***, Panama [1, 2] |
|  | NA | NA | NA | No | 0 | 1 | Brazil [4] |
| ***Order: Didelphimorphia*** | | | | | | | |
| Didelphidae | *Caluromys* | *C. philander* | Bare-tailed woolly opossum | Yes | 1 | 5 | French Guiana **[9]***, Brazil [18] |
|  |  | *C. derbianus* | Derby's wooly opossum | No | 0 | 3 | Colombia [19] |
|  | *Didelphis* | *D. albiventris* | White-eared opossum | Yes | 2 | 19 | French Guiana **[9]***, Brazil [18] |
|  |  | *D. marsupialis* | Common opossum | Yes | 1 | 87 | French Guiana **[9]***, Panama [1, 2],  Brazil [18, 21],  Colombia [19] |
|  |  | NA | NA | No | 0 | 43 | Panama/Colombia [17] |
|  | *Marmosa* | *M. cinerea* | Woolly mouse opossum | No | 0 | NA | Brazil |
|  |  | *M. murina* | Linnaeus's mouse opossum | No | 0 | 2 | Colombia [19] |
|  |  | NA | NA | Yes | 7 | 296 | Brazil **[15]***  Panama [1], Panama/Colombia [17], |
|  | *Metachirus* | *M. nudicaudatus* | Brown four-eyed opossum | No | 0 | 21 | Panama [1],  French Guiana [9],  Brazil [18] |
|  |  | NA | NA | No | 0 | 37 | Panama/Colombia [17] |
|  | *Monodelphis* | *M. americana* | Northern three-striped opossum | No | 0 | NA | Brazil [18] |
|  | *Philander* | *P. opossum* | Gray four-eyed opossum | Yes | 5 | 27 | French Guiana **[9]***, Brazil [18] |
|  |  | *P. nudicaudatus* | Gray and black four-eyed opossum | No | 0 | NA | Panama [2] |
|  | NA | NA | NA | Yes | 9 | 303 | Brazil **[15]*** [4]**,** Panama/Colombia [17] |
| ***Order: Carnivora*** | | | | | | | |
| Canidae | *Dusicyon* | *D. thous* | Crab-eating fox | No | 0 | 1 | Colombia [19] |
| Mustelidae | *Eira* | *E. barbara* | Tayra | No | 0 | N/A | Panama [2] |
|  | NA | NA | NA | No | 0 | 3 | Brazil [4] |
| Procyonidae | *Bassaricyon* | *B. gabii* | Olingo | No | 0 | NA | Panama [2] |
|  | *Bassariscus* | *B. sumichrasti* | Cacomistle | No | 0 | NA | Panama [2] |
|  | *Nasua* | *N. nasua* | South American coati | No | 0 | 9 | Panama [1, 2],  Peru [8] |
|  | *Potos* | *P. flavus* | Kinkajou | Yes | 1 | 16 | French Guiana **[9]***, Panama [1, 2] |
|  | *Procyon* | *P. canerivorus* | Crab-eating raccoon | No | 0 | NA | Panama [2] |
|  | NA | NA | NA | No | 0 | 4 | Brazil [4] |
| ***Order Artiodactyla*** | | | | | | | |
| Cervidae | *Mazama* | *M. americana* | Red brocket | No | 0 | 3 | Peru [8],  Panama [2] |
|  |  | NA | NA | No | 0 | 10 | French Guiana [9] |
| Tayassuidae | *Pecari* | *P. tajacu* | Collared peccary | Yes | 1 | 13 | Peru **[8]***,  French Guiana [9], Panama [2] |
| ***Order Crocodilia*** | | | | | | | |
| Alligatoridae | NA | NA | Caiman | No | 0 | 87 | Brazil [23] |
| ***Order Lagomorpha*** | | | | | | | |
| Leporidae | *Sylvilagus* | *S. brasiliensis* | Tapeti | No | 0 | 15 | Panama [1, 2] |
|  | *N/A* | *N/A* | N/A | No | 0 | 24 | Panama/Colombia [17] |
| ***Order Pilosa*** | | | | | | | |
| Bradypodidae | *Bradypus* | *B. variegatus* | Brown-throated sloth | No | 0 | 69 | Costa Rica [24],  Panama [1, 2],  Brazil [11] |
|  |  | *B. tridactylus* | Pale-throated sloth | Yes | 1 | 29 | French Guiana **[9]***, Brazil [18] |
|  |  | *B. torquatus* | Maned sloth | No | 0 | 22 | Brazil [11] |
|  |  | *N/A* | N/A | Yes | 1 | 4 | Brazil **[4]*,**  Peru [8] |
|  | *N/A* | *N/A* | N/A | No | 0 | 11 | Brazil [4] |
| Choloepodidae | *Choloepus* | *C. hoffmanni* | Hoffmann's two-toed sloth | No | 0 | 96 | Costa Rica [24],  Panama [1, 2] |
|  |  | *C. didactylus* | Linnaeus's two-toed sloth | Yes | 7 | 26 | French Guiana **[9]***, Brazil [18] |
| Myrmecophagidae | *Tamandua* | *T. mexicana* | Northern tamandua | No | 0 | NA | Panama [2] |
|  |  | *T. tetradactyla* | Southern tamandua | Yes | 6 | 40 | French Guiana **[9]***, Panama [1] |
| ***Order Squamata*** | | | | | | | |
| Teiidae | *Ameiva* | *A. ameiva* | South American ground lizard | Yes | 1 | NA | Brazil **[4]*** |
| Tropiduridae | *Tropidurus* | *T. torquatus* |  | Yes | 1 | NA | Brazil **[4]*** |
| Iguanidae | *Iguana* | NA | Iguana | No | 0 | 1 | Colombia [19] |
| NA | NA | NA | Lizard | No | 0 | 1 | Colombia [19] |
| ***Order Testudines*** | | | | | | | |
| Testudinidae | *Geochelone* | *G. denticulata* | Yellow-footed tortoise | No | 0 | NA | Brazil [18] |

NT: Neutralization test; HI: hemagglutination inhibition; MAYV: Mayaro virus

^#^Total is pooled across all studies. A value of NA indicates that a study reported testing an animal for MAYV but did not specify how many were tested.

^*^Indicates the location where the positive animal was found and the citation for the study that reported the positive animal.

**S3 Table. MAYV positivity by taxa of wild birds in included studies**

| **Family** | **Genus** | **Species** | **Common Name** | **MAYV Positive Samples** | **Total Positive** | **Total Tested^#^** | **Country of study*** |
| --- | --- | --- | --- | --- | --- | --- | --- |
| ***Order: Passeriformes*** | | | | | | | |
| Bombycillidae | *Bombycilla* | *B. cedrorum* | Cedar waxwing | No | 0 | 1 | USA [25] |
| Cardinalidae | *Cyanocompsa* | *C. cyanoides* | Blue-black grosbeak | No | 0 | 2 | Colombia [19],  Brazil [18] |
|  | *Guiraca* | *G. caerulea* | Blue grosbeak | No | 0 | 8 | USA [25] |
|  | *Habia* | *H. rubica* | Red-crowned ant tanager | No | 0 | NA | Brazil [18] |
|  | *Passerina* | *P. ciris* | Painted bunting | No | 0 | 12 | USA [25] |
|  |  | *P. cyanea* | Indigo bunting | No | 0 | 60 | USA [25] |
|  | *Pheucticus* | *P. ludovicianus* | Rose-breasted grosbeak | No | 0 | 52 | USA [25],  Colombia [19] |
|  | *Piranga* | *P.* *olivacea* | Scarlet tanager | No | 0 | 70 | USA [25] |
|  |  | *P. rubra* | Summer tanager | No | 0 | 110 | USA [25] |
|  | *Richmondena* | *R. cardinalis* | Northern cardinal | No | 0 | 8 | USA [25] |
| Conopophagidae | *Conopophaga* | *C. roberti* | Hooded gnateater | No | 0 | 1 | Brazil [21] |
|  |  | *C. aurita* | Chestnut-belted gnateater | No | 0 | NA | Brazil [18] |
| Dendrocolaptidae | *Hylexetastes* | *H. perrotii* | Red-billed woodcreeper | No | 0 | NA | Brazil [18] |
|  | *Xiphorhynchus* | *X. picus* | Straight-billed Woodcreeper | No | 0 | NA | Brazil [18] |
|  | NA | NA | NA | Yes | 1 | 97 | Brazil **[4]*** |
| Formicariidae | *Formicarius* | *F. analis* | Black-faced antthrush | No | 0 | NA | Brazil [18] |
|  | NA | NA | NA | Yes | 5 | 444 | Brazil **[4]*** |
| Fringillidae | *Cyanocompsa* | *C. cyanoides* | Blue-black grosbeak | No | 0 | 2 | Colombia [19] |
|  | *Spinus* | *S. psaltria* | Lesser goldfinch | No | 0 | 13 | Colombia [19] |
|  |  | NA | NA | No | 0 | 2 | Colombia [19] |
|  | *Tiaris* | NA | NA | No | 0 | 1 | Colombia [19] |
|  | NA | NA | NA | Yes | 6 | 131 | Brazil **[4]*** |
| Furnariidae | *Automolus* | *A. infuscatus* | Olive-backed foliage-gleaner | No | 0 | NA | Brazil [18] |
|  |  | *A. rufipileatus* | Chestnut-crowned Foliage-gleaner | No | 0 | NA | Brazil [18] |
|  | *Deconychura* | *D. longicauda* | Long-tailed woodcreeper | No | 0 | NA | Brazil [18] |
|  | *Dendrocincla* | *D. fuliginosa* | Plain-brown woodcreeper | No | 0 | NA | Brazil [18] |
|  | *Glyphorynchus* | *G. spirurus* | Wedge-billed woodcreeper | No | 0 | NA | Brazil [18] |
|  | *Philydor* | *P. pyrrhodes* | Cinnamon-rumped foliage-gleaner | No | 0 | NA | Brazil [18] |
|  | *Sclerurus* | *S. mexicanus* | Tawny-throated leaftosser | No | 0 | NA | Brazil [18] |
|  | *Synallaxis* | *S. albescens* | Pale-breasted spinetail | No | 0 | 1 | Colombia [19] |
|  |  | *S. gujanensis* | Plain-crowned Spinetail | No | 0 | NA | Brazil [18] |
|  | *Xiphorhynchus* | *X. ocellatus* | Ocellated woodcreeper | No | 0 | 1 | Brazil [21] |
|  |  | *X. spixii* | Spix's woodcreeper | No | 0 | NA | Brazil [18] |
| Grallariidae | *Myrmothera* | *M. campanisona* | Thrush-like antpitta | No | 0 | 1 | Brazil [21] |
| Hirundinidae | *Hirundo* | *H. rustica* | Barn swallow | No | 0 | 4 | USA [25] |
|  | *Notiochelidon* | *N. cyanoleuca* | Blue-and-white swallow | No | 0 | 3 | Colombia [19] |
|  | *Stelgidopteryx* | *S. ruficollis* | Southern rough-winged swallow | No | 0 | 1 | Colombia [19] |
| Icteridae | *Agelaius* | *A. icterocephalus* | Yellow-hooded blackbird | No | 0 | 3 | Colombia [19] |
|  |  | *A. phoeniceus* | Red-winged blackbird | No | 0 | 65 | USA [25] |
|  | *Cacicus* | *C. cela* | Yellow-rumped cacique | No | 0 | NA | Brazil [18] |
|  | *Dolichonyx* | *D. orizivorus* | Bobolink | No | 0 | 8 | USA [25] |
|  | *Icterus* | *I. galbula* | Baltimore oriole | No | 0 | 2 | USA [25] |
|  |  | *I. prosthemelas* | Black-cowled oriole | No | 0 | 2 | Panama [20] |
|  |  | *I. spurius* | Orchard oriole | Yes | 1 | 223 | USA **[25]*** |
| Mimidae | *Dumetella* | *D. carolinensis* | Gray catbird | No | 0 | 134 | USA [25] |
| Motacillidae | *Anthus* | *A. hellmayri* | Hellmayr’s pippit | No | 0 | NA | Brazil [26] |
| Paridae | *Parus* | *P. bicolor* | Tufted titmouse | No | 0 | 1 | USA [25] |
| Parulidae | *Dendroica* | *D. striata* | Blackpoll warbler | No | 0 | 10 | USA [25] |
|  |  | *D. magnolia* | Magnolia warbler | No | 0 | 4 | USA [25] |
|  |  | *D. petechia* | American yellow warbler | No | 0 | 4 | USA [25] |
|  | *Geothlypis* | *G. aequinoctialis* | Masked yellowthroat | No | 0 | NA | Brazil [18] |
|  |  | *G. semiflava* | Olive-crowned yellowthroat | No | 0 | 1 | Colombia [19] |
|  |  | *G. trichas* | Common yellowthroat | No | 0 | 2 | USA [25] |
|  | *Helmitheros* | *H. vermivorus* | Worm-eating warbler | No | 0 | 5 | USA [25] |
|  | *Limnothlypis* | *L. swainsonii* | Swainson's warbler | No | 0 | 2 | USA [25] |
|  | *Mniotilta* | *M. varia* | Black-and-white warbler | No | 0 | 5 | USA [25] |
|  | *Oporornis* | *O. formosus* | Kentucky warbler | No | 0 | 6 | USA [25] |
|  |  | *O. philadelphiae* | Mourning warbler | No | 0 | 1 | Colombia [19] |
|  | *Protonotaria* | *P. citrea* | Prothonotary warbler | No | 0 | 36 | USA [25] |
|  | *Seiurus* | *S. aurocapillus* | Ovenbird | No | 0 | 31 | USA [25] |
|  |  | *S. motacilla* | Louisiana waterthrush | No | 0 | 1 | USA [25] |
|  |  | *S. noveboracensis* | Northern waterthrush | No | 0 | 35 | USA [25] |
|  | *Setophaga* | *S. ruticilla* | American redstart | No | 0 | 12 | USA [25] |
|  | *Vermivora* | *V. peregrina* | Tennessee warbler | No | 0 | 1 | USA [25] |
|  |  | *V. pinus* | Blue-winged warbler | No | 0 | 2 | USA [25] |
|  | *Wilsonia* | *W. citrina* | Hooded warbler | No | 0 | 13 | USA [25] |
| Passerellidae | *Arremon* | *A. tactiturnus* | Pectoral sparrow | Yes | 1 | NA | Brazil **[14]*** [18] |
|  | *Zonotrichia* | *Z. albicollis* | White-throated sparrow | No | 0 | 8 | USA [25] |
|  |  | *Z. capensis* | Rufous-collared sparrow | No | 0 | 20 | Colombia [19] |
| Pipridae | *Pipra* | *P. fasciicauda* | Band-tailed manakin | No | 0 | NA | Brazil [18] |
|  |  | *P. rubrocapilla* | Red-headed manakin | No | 0 | NA | Brazil [18] |
|  | NA | NA | NA | Yes | 1 | 229 | Brazil **[4]*** |
| Thamnophilidae | *Cercomacra* | *C. tyrannina* | Dusky antbird | Yes | 1 | N/A | Brazil **[14]*** |
|  |  | *C. nigrescens* | Blackish antbird | No | 0 | NA | Brazil [18] |
|  | *Formicivora* | *F. grisea* | Southern white-fringed antwren | Yes | 1 | NA | Brazil **[14]*** |
|  | *Hylophylax* | *H. naevia* | Spot-backed antbird | No | 0 | NA | Brazil [18] |
|  | *Hypocnemis* | *H. cantator* | Guianan warbling antbird | No | 0 | NA | Brazil [18] |
|  | *Phlegopsis* | *P. nigromaculata* | Black-spotted bare-eye | No | 0 | NA | Brazil [18] |
|  | *Percnostola* | *P. rufifrons* | Black-headed antbird | No | 0 | NA | Brazil [18] |
|  | *Pygiptila* | *P. stellaris* | Spot-winged antshrike | No | 0 | NA | Brazil [18] |
|  | *Pyriglena* | *P. leuconota* | East Amazonian fire-eye | No | 0 | NA | Brazil [18] |
|  | *Taraba* | *T. major* | Great antshrike | No | 0 | NA | Brazil [18] |
|  | *Thamnomanes* | *T. caesius* | Cinereous antshrike | No | 0 | NA | Brazil [18] |
|  | *Thamnophilus* | *T. aethiops* | White-shouldered antshrike | No | 0 | 1 | Brazil [18, 21] |
|  |  | *T. amazonicus* | Amazonian antshrike | No | 0 | NA | Brazil [18] |
| Thraupidae | *Coereba* | *C. flaveola* | Bananaquit | No | 0 | 2 | Colombia [19] |
|  | *Oryzoborus* | *O. angolensis* | Chestnut-bellied seed finch | No | 0 | NA | Brazil [18] |
|  | *Saltator* | *S. albicollis* | Lesser Antillean saltator | No | 0 | 7 | Colombia [19] |
|  |  | *S. coerulescens* | Greyish saltator | No | 0 | NA | Brazil [18] |
|  |  | *S. maximus* | Buff-throated saltator | No | 0 | NA | Brazil [18] |
|  | *Sicalis* | *S. luteola* | Grassland yellow finch | No | 0 | NA | Brazil [26] |
|  | *Ramphocelus* | *R. carbo* | Silver-beaked tanager | No | 0 | NA | Brazil [18] |
|  |  | *R. dimidiatus* | Crimson-backed tanager | No | 0 | 1 | Colombia [19] |
|  |  | *R. paserinii* | Scarlet-rumped tanager | No | 0 | 201 | Panama [20] |
|  | *Sporophila* | *S. caerulescens* | Double-collared seedeater | No | 0 | NA | Brazil [18] |
|  |  | *S. intermedia* | Grey seedeater | No | 0 | 11 | Colombia [19] |
|  |  | *S. minuta* | Ruddy-breasted seedeater | No | 0 | 5 | Colombia [19] |
|  |  | *S. nigricollis* | Yellow-bellied seedeater | No | 0 | 5 | Colombia [19] |
|  |  | NA | NA | No | 0 | 7 | Colombia [19] |
|  | *Tachyphonus* | *T. rufus* | White-lined tanager | No | 0 | NA | Brazil [18] |
|  | *Thraupis* | *T. episcopus* | Blue-gray tanager | No | 0 | 6 | Colombia [19],  Brazil [18] |
|  |  | *T. palmarum* | Palm tanager | No | 0 | NA | Brazil [18] |
|  | *Volatinia* | *V. jacarina* | Blue-black grassquit | No | 0 | 1 | Colombia [19] |
| Tityridae | *Onychorhynchus* | *O. coronatus* | Amazonian Royal Flycatcher | No | 0 | NA | Brazil [18] |
|  | *Schiffornis* | *S. turdina* | Brown-winged schiffornis | No | 0 | 1 | Brazil [21] |
| Troglodytidae | *Microcerculus* | *M. marginatus* | Scaly-breasted Wren | No | 0 | NA | Brazil [18] |
|  | *Thryothorus* | *T. genibarbis* | Moustached wren | No | 0 | NA | Brazil [18] |
|  |  | *T. ludovicianus* | Carolina wren | No | 0 | 1 | USA [25] |
|  | *Troglodytes* | *T. aedon* | House wren | No | 0 | 3 | Colombia [19] |
| Turdidae | *Catharus* | *C. ustulatus* | Swainson's thrush | No | 0 | 12 | Colombia [19],  USA [25] |
|  | *Hylocichla* | *H. fuscescens* | Veery | No | 0 | 30 | USA [25] |
|  |  | *H. minima* | Gray-cheeked thrush | No | 0 | 43 | USA [25] |
|  |  | *H. mustellina* | Wood thrush | No | 0 | 16 | USA [25] |
|  | *Turdus* | *T. fumigatus* | Cocoa thrush | No | 0 | NA | Brazil [18] |
|  |  | *T. grayi* | Clay-colored thrush | No | 0 | 43 | Panama [20] |
|  |  | *T. ignobilis* | Black-billed thrush | No | 0 | 22 | Colombia [19] |
|  |  | NA | NA | No | 0 | 2 | Colombia [19] |
| Tyrannidae | *Attila* | *A. spadiceus* | Bright-rumped attila | No | 0 | NA | Brazil [18] |
|  | *Contopus* | *C. virens* | Eastern wood pewee | No | 0 | 2 | USA [25] |
|  | *Elaenia* | *E. flavogaster* | Yellow-bellied elaenia | No | 0 | 10 | Colombia [19] |
|  | *Empidonax* | *E. virescens* | Acadian flycatcher | No | 0 | 5 | USA [25] |
|  | *Fluvicola* | *F. pica* | Pied water tyrant | No | 0 | 2 | Colombia [19] |
|  | *Mecocerculus* | *M. leucophrys* | White-throated tyrannulet | No | 0 | NA | Brazil [18] |
|  | *Myiozetetes* | *M. crinitus* | Great crested flycatcher | No | 0 | 11 | USA [25] |
|  |  | *M. granadensis* | Grey-capped flycatcher | No | 0 | 16 | Panama [20] |
|  |  | *M. similis* | Social flycatcher | No | 0 | 17 | Panama [20] |
|  |  | *M. cayanensis* | Rusty-margined flycatcher | No | 0 | 4 | Colombia [19] |
|  |  | NA | NA | No | 0 | 21 | Panama [20] |
|  | *Pitangus* | *P. sulphuratus* | Great kiskadee | No | 0 | 7 | Brazil [27], Colombia [19] |
|  | *Pyrocephalus* | *P. rubinus* | Scarlet flycatcher | No | 0 | 1 | Colombia [19] |
|  | *Todirostrum* | *T. cinereum* | Common tody-flycatcher | No | 0 | 1 | Colombia [19] |
|  |  | NA | NA | No | 0 | 1 | Colombia [19] |
|  | *Tyrannus* | *T. melancholicus* | Tropical kingbird | Yes | 1 | 4 | Brazil **[14]*** [27]**,** Colombia [19] |
|  |  | *T. tyrannus* | Eastern kingbird | No | 0 | 29 | USA [25] |
|  | NA | NA | NA | Yes | 1 | 103 | Brazil **[4]*,**  Colombia [19] |
| Vireonidae | *Vireo* | *V. altiloquus* | Black-whiskered Vireo | No | 0 | 3 | USA [25] |
|  |  | *V. flavifrons* | Yellow-throated vireo | No | 0 | 4 | USA [25] |
|  |  | *V. griseus* | White-eyed vireo | No | 0 | 4 | USA [25] |
|  |  | *V. olivaceus* | Red-eyed vireo | No | 0 | 158 | Colombia [19],  USA [25] |
| ***Order Pelecaniformes*** | | | | | | | |
| Ardeidae | *Bubulcus* | *B. ibis* | Cattle egret | No | 0 | 1 | Colombia [19] |
|  | *Butorides* | *B. virescens* | Green heron | No | 0 | 20 | Panama [20] |
|  | *Egretta* | *E. caerulea* | Little blue heron | No | 0 | 14 | Panama [20] |
|  | *Tigrisoma* | *T. lineatum* | Rufescent tiger heron | No | 0 | 1 | Peru [8] |
| ***Order Piciformes*** | | | | | | | |
| Bucconidae | *Malacoptila* | *M. rufa* | Rufous-necked puffbird | No | 0 | NA | Brazil [18] |
| Picidae | *Chrysoptilus* | *C. punctigula* | Spot-breasted woodpecker | No | 0 | 1 | Colombia [19] |
|  | *Dendrocopos* | *D. villosus* | Hairy Woodpecker | No | 0 | 3 | USA [25] |
|  | *Dryocopus* | *D. lineatus* | Lineated woodpecker | No | 0 | NA | Brazil [18] |
|  | *Picummus* | NA | NA | No | 0 | 1 | Colombia [19] |
|  | *Sphyrapicus* | *S. varius* | Yellow-bellied sapsucker | No | 0 | 3 | USA [25] |
| Ramphastidae | *Ramphastos* | *R. sulfuratus* | Keel-billed toucan | No | 0 | 28 | Panama [20] |
| ***Order: Charadriiformes*** | | | | | | | |
| Charadriidae | *Charadrius* | *C. collaris* | Collared plover | No | 0 | 8 | Brazil [27] |
|  |  | *C. semipalmatus* | Semipalmated plover | No | 0 | 3 | Brazil [26, 27] |
|  |  | *C. wilsonia* | Wilson's plover | No | 0 | 1 | Brazil [27] |
|  | *Pluvialis* | *P. squatarola* | Grey plover | Yes | 1 | 4 | Brazil **[27]*** [26] |
| Hematopodidae | *Haematopus* | *H. palliatus* | American oystercatcher | Yes | 1 | 6 | Brazil **[28]*** |
| Jacanidae | *Jacana* | *J. jacana* | Wattled jacana | No | 0 | 6 | Colombia [19] |
| Laridae | *Rynchops* | *R.niger* | Black skimmer | No | 0 | 9 | Brazil [26-28] |
|  | *Sterna* | *S. eurygnatha* | Cayenne tern | Yes | 1 | 7 | Brazil **[28]*** [26] |
|  |  | *S. hirundo* | Common tern | Yes | 23 | 342 | Brazil **[28]*** [26] |
|  |  | *S. hirundinaceae* | South American tern | No | 0 | NA | Brazil [26] |
|  |  | *S. maxima* | Royal tern | Yes | 1 | 1 | Brazil **[28]*** |
|  |  | *S. nilotica* | Gull-billed tern | Yes | 1 | 1 | Brazil **[28]*** |
|  |  | *S. superciliaris* | Yellow-billed tern | Yes | 2 | 12 | Brazil **[28]*** [26, 27] |
|  |  | *S. trudeaui* | Snowy-crowned tern | Yes | 12 | 56 | Brazil **[28]*** [26] |
| Recurvirostridae | *Himantopus* | *H. himantopus* | Black-winged stilt | No | 0 | NA | Brazil [26] |
| Scolopacidae | *Actitis* | *A. macularius* | Spotted sandpiper | Yes | 2 | 26 | Brazil **[27]*,**  USA [25],  Colombia [19] |
|  | *Arenaria* | *A. interpres* | Ruddy turnstone | Yes | 8 | 36 | Brazil **[28]*** [26, 27] |
|  | *Calidris* | *C. alba* | Sanderling | No | 0 | NA | Brazil [26] |
|  |  | *C. canutus* | Red knot | Yes | 7 | 54 | Brazil **[28]*** [26, 27] |
|  |  | *C. fuscicollis* | White-rumped sandpiper | Yes | 1 | 11 | Brazil **[28]*** [26] |
|  |  | *C. minutilla* | Least sandpiper | Yes | 1 | 6 | Brazil **[27]*** |
|  |  | *C. pusilla* | Semipalmated sandpiper | Yes | 1 | 30 | Brazil **[27]*** |
|  | *Limosa* | *L. haemastica* | Hudsonian godwit | Yes | 5 | 17 | Brazil **[28]*** [26] |
|  | *Tringa* | *T. flavipes* | Lesser yellowlegs | Yes | 4 | 5 | Brazil **[28]*** [26] |
|  |  | *T. melanoleuca* | Greater yellowlegs | No | 0 | 1 | Brazil [26, 27] |
|  |  | *T. solitaria* | Solitary sandpiper | No | 0 | 1 | Colombia [19] |
| ***Order Caprimulgiformes*** | | | | | | | |
| Caprimulgidae | *Caprimulgus* | NA | NA | No | 0 | 1 | Colombia [19] |
|  | NA | NA | NA | Yes | 1 | 5 | Brazil **[4]*** |
| ***Order Columbiformes*** | | | | | | | |
| Columbidae | *Geotrygon* | *G. montana* | Ruddy quail-dove | No | 0 | 1 | Brazil [21] [18] |
|  |  | *G. violacea* | Violaceous quail-dove | No | 0 | NA | Brazil [18] |
|  | *Columbina* | *C. talpacoti* | Ruddy ground dove | No | 0 | 6 | Colombia [19] |
|  |  | *C. passerina* | Common ground dove | No | 0 | 6 | Colombia [19], Brazil [18] |
|  | *Leptotila* | *L. rufaxilla* | Grey-fronted dove | No | 0 | NA | Brazil [18] |
|  |  | *L. plumbeiceps* | Grey-headed dove | No | 0 | 5 | Colombia [19] |
|  |  | NA | NA | No | 0 | 2 | Colombia [19] |
|  | *Columbigallina* | NA | NA | Yes | 34 | 121 | Brazil **[15]***,  Colombia [19] |
|  | NA | NA | NA | Yes | 1 | 34 | Brazil **[4]*** |
| ***Order Coraciiformes*** | | | | | | | |
| Momotidae | *Momotus* | *M. momota* | Amazonian motmot | No | 0 | NA | Brazil [18] |
| Alcedinidae | *Chloroceryle* | *C. inda* | Green-and-rufous kingfisher | No | 0 | NA | Brazil [18] |
| ***Order Podicipediformes*** | | | | | | | |
| Podicipedidae | *Podiceps* | *P. dominicus* | Least grebe | No | 0 | 2 | Colombia [19] |
|  |  | *P. major* | Great grebe | No | 0 | NA | Brazil [26] |
| ***Order Psittaciformes*** | | | | | | | |
| Psittacidae | *Ara* | *A. ararauna* | Blue and yellow macaw | No | 0 | 2 | Peru [8] |
|  | *Forpus* | *F. conspicillatus* | Spectacled parrotlet | No | 0 | 1 | Colombia [19] |
| ***Order Cuculiformes*** | | | | | | | |
| Cuculidae | *Coccyzus* | *C. americanus* | Yellow-billed cuckoo | No | 0 | 40 | USA [25] |
|  |  | *C. erythropthalmus* | Black-billed cuckoo | No | 0 | 3 | USA [25] |
|  |  | *C. pumilus* | Dwarf cuckoo | No | 0 | 1 | Colombia [19] |
|  |  | NA | NA | No | 0 | 2 | Colombia [19] |
|  | *Crotophaga* | *C. ani* | Smooth-billed ani | No | 0 | 2 | Colombia [19] |
|  |  | *C. sulcirostris* | Groove-billed ani | No | 0 | 16 | Panama [20] |
| ***Order Galliformes*** | | | | | | | |
| Cracidae | *Ortalis* | *O. guttata* | Speckled chachalaca | No | 0 | 1 | Peru [8] |
| ***Order Strigiformes*** | | | | | | | |
| Strigidae | *Glaucidium* | *G. brasilianum* | Ferruginous pygmy owl | No | 0 | NA | Brazil [18] |
|  | *Otus* | *O. choliba* | Tropical screech owl | No | 0 | 3 | Colombia [19] |
| ***Order Apodiformes*** | | | | | | | |
| Trochilidae | *Gloucis* | *G. aenea* | Bronzy hermit | No | 0 | 1 | Colombia [19] |
|  | NA | NA | NA | No | 0 | 2 | Colombia [19] |
| ***Order Gruiformes*** | | | | | | | |
| Rallidae | *Gallinula* | *G. chloropus* | Common moorhen | No | 0 | 3 | Colombia [19] |
|  | *Rallus* | *R. longirostris* | Mangrove rail | No | 0 | 2 | Brazil [27] |
| ***Order Tinamiformes*** | | | | | | | |
| Tinamidae | *Tinamus* | *T. major* | Great tinamou | No | 0 | 2 | Peru [8] |

NT: Neutralization test; HI: hemagglutination inhibition; MAYV: Mayaro virus

^#^Total is pooled across all studies. A value of NA indicates that a study reported testing an animal for MAYV but did not specify how many were tested.

^*^Indicates the location where the positive animal was found and the citation for the study that reported the positive animal.

**S4 Table. MAYV positivity in domestic or sentinel animals studied**

| **Animal Type** | **MAYV positive Samples?** | **Total Positive** | **Total Tested^#^** | **Positivity Confirmed by NT?^%^** | **Country of study*** |
| --- | --- | --- | --- | --- | --- |
| Domestic Horse | Yes | 26 | 1096 | Yes | Brazil **[29]* [23]*** [30] |
|  |  | 51 | 859 | No | Brazil **[31]* [32]*** [26] |
| Domestic Cattle/Buffalo | Yes | 14 | 1103 | No | Brazil **[33]*,**  Colombia [19] |
| Domestic Dog | Yes | 2 | 22 | No | Brazil **[28]*** [26]  Colombia [19] |
| Sentinel Monkeys | Yes | 2 | 14 | No | Panama **[17]*** |
| Domestic Donkey | Yes | 1 | 24 | Yes | Brazil **[23]*** |
| Sentinel Hamster | Yes | 1 | NA | NA (RT-PCR) | Venezuela **[34]*** |
| Domestic Mule | No | 0 | 30 | NA | Brazil [23] |
| Domestic Unspecified Equid | No | 0 | 3 | NA | Brazil [23] |
| Domestic Sheep | No | 0^^^ | 622 | NA | Brazil [23, 33] |
| Domestic Hen/Chicken | No | 0 | 67 | NA | Brazil [4, 26, 28]  Colombia [19] |
| Domestic Pig | No | 0 | 13 | NA | Brazil [26],  Colombia [19] |
| Domestic Duck | No | 0 | 11 | NA | Brazil [4, 26] |
| Sentinel *Aotus nancymae* monkeys | No | 0 | 20 | NA | Peru [35] |

NT: Neutralization test; HI: hemagglutination inhibition; RT-PCR: reverse transcription polymerase chain reaction; MAYV: Mayaro virus

^%^*Yes* indicates that MAYV positivity was confirmed with an NT. *No* indicates that MAYV positivity was based on HI test only. *NA* indicates that no positivity was reported.

^#^Total is pooled across all studies. A value of NA indicates that a study reported testing an animal for MAYV but did not specify how many were tested.

^*^Indicates the location where the positive animal was found and the citation for the study that reported the positive animal.

^^^Neutralizing antibodies detected but did not meet the study’s diagnostic criteria.

**S5 Table. Pooled prevalence table (random effects using GLMM with logit transformation)**

| **Order** | **Positives Included^1^** | **Studies (n)** | **Total (n)** | **Positive (n)** | **Pooled Prev. (%)** | **95% CI** | **I^2^ (%)** | ***τ*^2^** | **P-value** |
| --- | --- | --- | --- | --- | --- | --- | --- | --- | --- |
| ***Mammals*** | | | | | | | | | |
| Primate | HI and NT | 13 | 897 | 153 | 8.7 | 3.1; 22.0 | 96 | 3.2572 | <0.01 |
|  | NT only | 13 | 858 | 114 | 0.7 | 0.0; 9.1 | 98 | 11.7937 | 0.05 |
| Pilosa | HI and NT | 7 | 297 | 15 | 0.5 | 0.0; 22.4 | 90 | 9.8670 | 1.00 |
|  | NT only | 7 | 296 | 14 | 0.1 | 0.0; 72.1 | 92 | 13.7165 | 1.00 |
| Rodentia | HI and NT | 7 | 1557 | 90 | 2.3 | 0.5; 10.3 | 93 | 2.6550 | 0.99 |
|  | NT only | 7 | 1486 | 19 | 0.4 | 0.0; 8.7 | 95 | 9.4525 | 0.99 |
| Domestic Equids | HI and NT | 6 | 1955 | 41 | 1.7 | 0.4; 6.6 | 92 | 2.0474 | <0.01 |
|  | NT only | 6 | 1940 | 26 | 0.2 | 0.0; 6.4 | 96 | 7.6678 | <0.01 |
| Didelphimorphia | HI and NT | 6 | 369 | 25 | 2.0 | 0.2; 20.2 | 85 | 2.1787 | 1.00 |
|  | NT only | 6 | 353 | 9 | 0.1 | 0.0; 51.4 | 90 | 10.4733 | 1.00 |
| Carnivora Order | HI and NT | 5 | 40 | 2 | 4.8 | 0.5; 32.4 | 3 | 0.0522 | 1.00 |
|  | NT only | 5 | 40 | 2 | 4.8 | 0.5; 32.4 | 3 | 0.0522 | 1.00 |
| Cingulata Order | HI and NT | 4 | 70 | 6 | 10.0 | 2.7; 30.8 | 34 | 0.5502 | 0.14 |
|  | NT only | 4 | 70 | 6 | 10.0 | 2.7; 30.8 | 34 | 0.5502 | 0.14 |
| Artiodactyla | HI and NT | 2 | 26 | 1 | 3.8 | 0.5; 22.8 | 0 | 0 | 1.00 |
|  | NT only | 2 | 26 | 1 | 3.8 | 0.5; 22.8 | 0 | 0 | 1.00 |
| ***Birds^2^*** | | | | | | | | | |
| Charadriiformes | HI and NT | 3 | 641 | 71 | 9.2 | 4.4; 18.2 | 29 | 0.0846 | 0.19 |
| Passeriformes | HI and NT | 4 | 1166 | 14 | 1.2 | 0.7; 2.0 | 0 | 0 | 1.00 |
| Columbiformes | HI and NT | 4 | 171 | 35 | 5.0 | 0.5; 36.8 | 74 | 2.4841` | 0.10 |

MAYV: Mayaro virus; HI: hemagglutination inhibition; NT: neutralization test; CI: confidence interval

^1^ The first analysis (HI and NT) included all positive samples, regardless of test method. A sensitivity analysis was conducted that included only positive samples that were confirmed with NT.

^2^ Only one study reporting MAYV positivity in birds used confirmatory NT. Therefore, a sensitivity analysis was not conducted.

**S6 Table. Primate genera pooled prevalence table (random effects with Freeman-Tukey double arcsine transformation)**

| **Primate Genus** | **Positives Included^1^** | **Studies (n)** | **Total (n)** | **Positive (n)** | **Pooled Prev. (%)** | **95% CI** | **I^2^ (%)** | ***τ*^2^** | **P-value** |
| --- | --- | --- | --- | --- | --- | --- | --- | --- | --- |
| *Cebus/Sapajus* | HI and NT | 9 | 316 | 25 | 3.7 | 0.0; 11.1 | 61 | 0.0132 | <0.01 |
|  | NT only | 9 | 293 | 2 | 0.0 | 0.0; 0.0 | 13 | 0.0014 | 0.32 |
| *Alouatta* | HI and NT | 8 | 213 | 63 | 32.2 | 0.0; 79.2 | 95 | 0.2257 | <0.01 |
|  | NT only | 8 | 206 | 56 | 20.8 | 0.0; 68.9 | 94 | 0.2263 | <0.01 |
| *Callithrix* | HI and NT | 3 | 123 | 32 | 17.8 | 8.6; 28.5 | 0 | 0 | 0.54 |
|  | NT only | 3 | 123 | 32 | 17.8 | 8.6; 28.5 | 0 | 0 | 0.54 |
| *Saguinus* | HI and NT | 2 | 74 | 8 | 6.3 | 0.0; 35.5 | 90 | 0.0628 | <0.01 |
|  | NT only | 2 | 74 | 8 | 6.3 | 0.0; 35.5 | 90 | 0.0628 | <0.01 |
| *Lagothrix* | HI and NT | 1 | 11 | 6 | 54.5 | 24.2; 83.3 | NA | NA | NA |
|  | NT only | 1 | 11 | 6 | 54.5 | 24.2; 83.3 | NA | NA | NA |
| *Saimiri* | HI and NT | 3 | 10 | 5 | 45.9 | 0.0; 100.0 | 62 | 0.1254 | 0.07 |
|  | NT only | 2 | 9 | 4 | 30.3 | 0.0; 98.5 | 76 | 0.1696 | 0.04 |
| *Aotus* | HI and NT | 2 | 10 | 1 | 6.3 | 0.0; 43.6 | 35 | 0.0253 | 0.21 |
|  | NT only | 2 | 9 | 0 | 0.0 | 0.0; 20.0 | 0 | 0 | 0.84 |

MAYV: Mayaro virus; HI: hemagglutination inhibition; NT: neutralization test; CI: confidence interval

^1^ The first analysis (HI and NT) included all positive samples, regardless of test method. A sensitivity analysis was conducted that included only positive samples that were confirmed with NT.

**S7 Table. Primate genera pooled prevalence table (random effects using GLMM with logit transformation)**

| **Primate Genus** | **Positives Included^1^** | **Studies (n)** | **Total (n)** | **Positive (n)** | **Pooled Prev. (%)** | **95% CI** | **I^2^ (%)** | ***τ*^2^** | **P-value** |
| --- | --- | --- | --- | --- | --- | --- | --- | --- | --- |
| *Cebus/Sapajus* | HI and NT | 9 | 316 | 25 | 7.5 | 3.5; 15.3 | 59 | 0.6576 | 0.15 |
|  | NT only | 9 | 293 | 2 | 0.3 | 0.0; 9.0 | 72 | 4.7055 | 1.00 |
| *Alouatta* | HI and NT | 8 | 213 | 63 | 24.0 | 2.2; 81.6 | 94 | 10.4411 | 1.00 |
|  | NT only | 8 | 206 | 56 | 10.4 | 0.3; 79.7 | 94 | 16.7758 | 1.00 |
| *Callithrix* | HI and NT | 3 | 123 | 32 | 26.0 | 19.0; 34.5 | 0 | 0 | 1.00 |
|  | NT only | 3 | 123 | 32 | 26.0 | 19.0; 34.5 | 0 | 0 | 1.00 |
| *Saguinus* | HI and NT | 2 | 74 | 8 | 3.7 | 0.1; 63.4 | 77 | 3.7403 | 1.00 |
|  | NT only | 2 | 74 | 8 | 3.7 | 0.1; 63.4 | 77 | 3.7403 | 1.00 |
| *Lagothrix* | HI and NT | 1 | 11 | 6 | 54.5 | 26.8; 79.7 | NA | NA | NA |
|  | NT only | 1 | 11 | 6 | 54.5 | 26.8; 79.7 | NA | NA | NA |
| *Saimiri* | HI and NT | 3 | 10 | 5 | 47.7 | 8.9; 89.5 | 51 | 1.7960 | 1.00 |
|  | NT only | 2 | 9 | 4 | 31.1 | 1.6; 92.8 | 60 | 2.3144 | 1.00 |
| *Aotus* | HI and NT | 2 | 10 | 1 | 10.0 | 1.4; 46.7 | 0 | 0 | 1.00 |
|  | NT only | 2 | 9 | 0 | NA^2^ | NA | NA | NA | NA |

MAYV: Mayaro virus; GLMM: generalized linear mixed model; HI: hemagglutination inhibition; NT: neutralization test; CI: confidence interval

^1^ The first analysis (HI and NT) included all positive samples, regardless of test method. A sensitivity analysis was conducted that included only positive samples that were confirmed with NT.

^2^ Cannot fit model due to zero events.

**S8 Table. Pooled prevalence table (****fixed effects with Freeman-Tukey double arcsine transformation)**

| **Order** | **Positives Included^1^** | **Studies (n)** | **Total (n)** | **Positive (n)** | **Pooled Prev. (%)** | **95% CI** | **I^2^ (%)** | ***τ*^2^** | **P-value** |
| --- | --- | --- | --- | --- | --- | --- | --- | --- | --- |
| ***Mammals*** | | | | | | | | | |
| Primate | HI and NT | 13 | 897 | 153 | 11.9 | 9.8; 14.2 | 95 | 0.0692 | <0.01 |
|  | NT only | 13 | 858 | 114 | 6.5 | 4.8; 8.4 | 96 | 0.0851 | <0.01 |
| Pilosa | HI and NT | 7 | 297 | 15 | 0.0 | 0.0; 0.1 | 84 | 0.0338 | <0.01 |
|  | NT only | 7 | 296 | 14 | 0.0 | 0.0; 0.0 | 82 | 0.0305 | <0.01 |
| Rodentia | HI and NT | 7 | 1557 | 90 | 2.1 | 1.2; 3.1 | 91 | 0.0160 | <0.01 |
|  | NT only | 7 | 1486 | 19 | 0.0 | 0.0; 0.0 | 90 | 0.0153 | <0.01 |
| Domestic Equids | HI and NT | 6 | 1955 | 41 | 0.2 | 0.0; 0.6 | 90 | 0.0085 | <0.01 |
|  | NT only | 6 | 1940 | 26 | 0.0 | 0.0; 0.0 | 90 | 0.0087 | <0.01 |
| Didelphimorphia | HI and NT | 6 | 369 | 25 | 4.2 | 2.0; 6.9 | 68 | 0.0101 | <0.01 |
|  | NT only | 6 | 353 | 9 | 0.1 | 0.0; 1.3 | 74 | 0.0141 | <0.01 |
| Carnivora Order | HI and NT | 5 | 40 | 2 | 0.1 | 0.0; 8.1 | 0 | 0 | 0.71 |
|  | NT only | 5 | 40 | 2 | 0.1 | 0.0; 8.1 | 0 | 0 | 0.71 |
| Cingulata Order | HI and NT | 4 | 70 | 6 | 0.5 | 0.0; 7.4 | 35 | 0.0198 | 0.20 |
|  | NT only | 4 | 70 | 6 | 0.5 | 0.0; 7.4 | 35 | 0.0198 | 0.20 |
| Artiodactyla | HI and NT | 2 | 26 | 1 | 1.6 | 0.0; 12.7 | 46 | 0.0172 | 0.17 |
|  | NT only | 2 | 26 | 1 | 1.6 | 0.0; 12.7 | 46 | 0.0172 | 0.17 |
| ***Birds*^2^** | | | | | | | | | |
| Charadriiformes | HI and NT | 3 | 641 | 71 | 9.5 | 7.2; 12.2 | 61 | 0.0045 | 0.08 |
| Passeriformes | HI and NT | 4 | 1166 | 14 | 0.0 | 0.0; 0.0 | 27 | 0.0010 | 0.25 |
| Columbiformes O | HI and NT | 4 | 171 | 35 | 11.7 | 5.6; 19.0 | 87 | 0.0591 | <0.01 |

MAYV: Mayaro virus; HI: hemagglutination inhibition; NT: neutralization test; CI: confidence interval

^1^ The first analysis included all positive samples, regardless of test method. The second analysis included only the positive samples that were confirmed with NT.

^2^ Only one study reporting MAYV positivity in birds used confirmatory NT.

**S9 Table. Pooled prevalence table (fixed effects using GLMM with logit transformation)**

| **Order** | **Positives Included^1^** | **Studies (n)** | **Total (n)** | **Positive (n)** | **Pooled Prev. (%)** | **95% CI** | **I^2^ (%)** | ***τ*^2^** | **P-value** |
| --- | --- | --- | --- | --- | --- | --- | --- | --- | --- |
| ***Mammals*** | | | | | | | | | |
| Primate | HI and NT | 13 | 897 | 153 | 17.1 | 14.7; 19.7 | 96 | 3.2572 | <0.01 |
|  | NT only | 13 | 858 | 114 | 13.3 | 11.2; 15.7 | 98 | 11.7937 | 0.05 |
| Pilosa | HI and NT | 7 | 297 | 15 | 5.1 | 3.1; 8.2 | 90 | 9.8670 | 1.00 |
|  | NT only | 7 | 296 | 14 | 4.7 | 2.8; 7.8 | 92 | 13.7165 | 1.00 |
| Rodentia | HI and NT | 7 | 1557 | 90 | 5.8 | 4.7; 7.1 | 93 | 2.6550 | 0.99 |
|  | NT only | 7 | 1486 | 19 | 1.3 | 0.8; 2.0 | 95 | 9.4525 | 0.99 |
| Domestic Equids | HI and NT | 6 | 1955 | 41 | 2.1 | 1.5; 2.8 | 92 | 2.0474 | <0.01 |
|  | NT only | 6 | 1940 | 26 | 1.3 | 0.9; 2.0 | 96 | 7.6678 | <0.01 |
| Didelphimorphia | HI and NT | 6 | 369 | 25 | 6.8 | 4.6; 9.8 | 85 | 2.1787 | 1.00 |
|  | NT only | 6 | 353 | 9 | 2.5 | 1.3; 4.8 | 90 | 10.4733 | 1.00 |
| Carnivora Order | HI and NT | 5 | 40 | 2 | 5.0 | 1.3; 17.9 | 3 | 0.0522 | 1.00 |
|  | NT only | 5 | 40 | 2 | 5.0 | 1.3; 17.9 | 3 | 0.0522 | 1.00 |
| Cingulata Order | HI and NT | 4 | 70 | 6 | 8.6 | 3.9; 17.8 | 34 | 0.5502 | 0.14 |
|  | NT only | 4 | 70 | 6 | 8.6 | 3.9; 17.8 | 34 | 0.5502 | 0.14 |
| Artiodactyla | HI and NT | 2 | 26 | 1 | 3.8 | 0.5; 22.8 | 0 | 0 | 1.00 |
|  | NT only | 2 | 26 | 1 | 3.8 | 0.5; 22.8 | 0 | 0 | 1.00 |
| ***Birds^28^*** | | | | | | | | | |
| Charadriiformes | HI and NT | 3 | 641 | 71 | 11.1 | 8.9; 13.7 | 29 | 0.0846 | 0.19 |
| Passeriformes | HI and NT | 4 | 1166 | 14 | 1.2 | 0.7; 2.0 | 0 | 0 | 1.00 |
| Columbiformes | HI and NT | 4 | 171 | 35 | 20.5 | 15.1; 27.2 | 74 | 2.4841` | 0.10 |

MAYV: Mayaro virus; HI: hemagglutination inhibition; NT: neutralization test; CI: confidence interval

^1^ The first analysis included all positive samples, regardless of test method. The second analysis included only the positive samples that were confirmed with NT.

^2^ Only one study reporting MAYV positivity in birds used confirmatory NT.

**S1 Fig. Funnel plots for estimates of MAYV seroprevalence in non-human animal reservoirs**

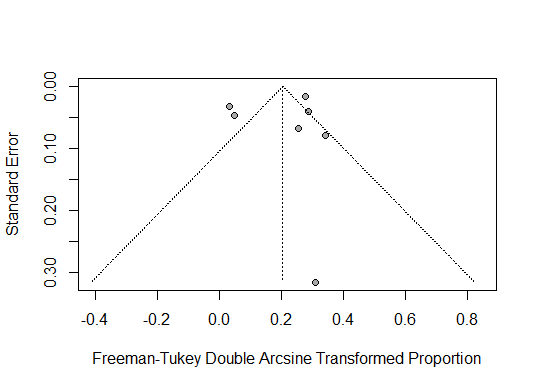

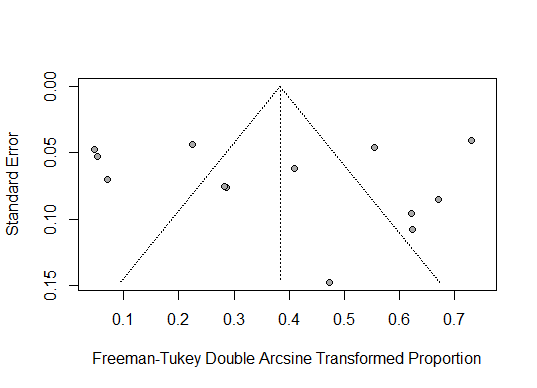

A. B.

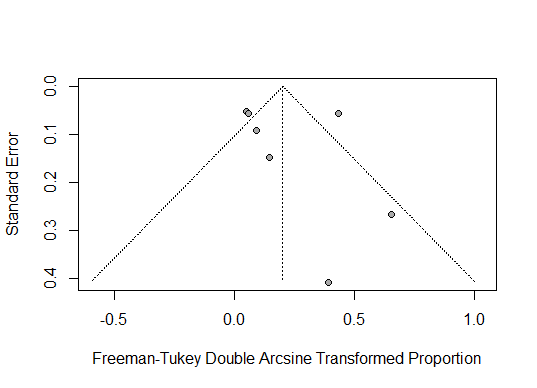

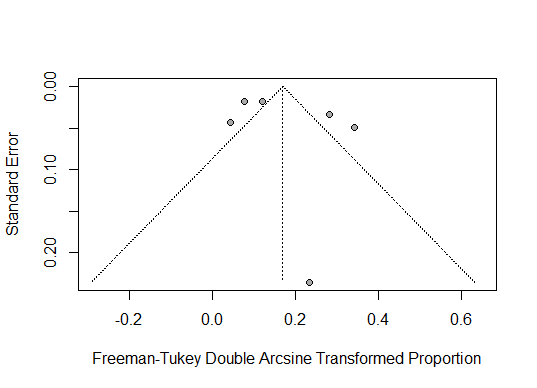
C. D.

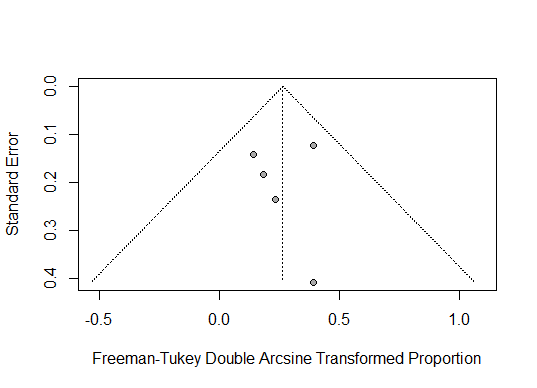

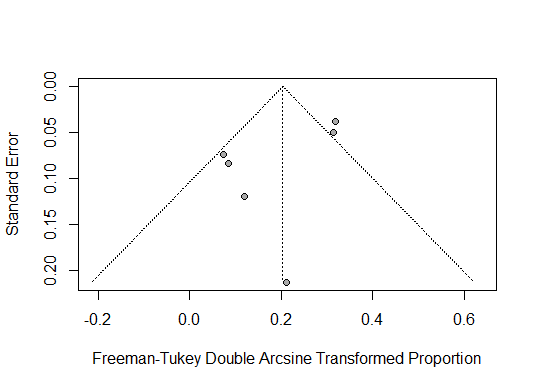
E. F.

Funnel plots presented for A) Primate order, B) Rodentia order, C) Domestic equids, D) Pilosa order, E) Didelphimorphia order, F) Carnivora order

**S2 Fig. Funnel plots for estimates of MAYV seroprevalence in non-human primate genera**

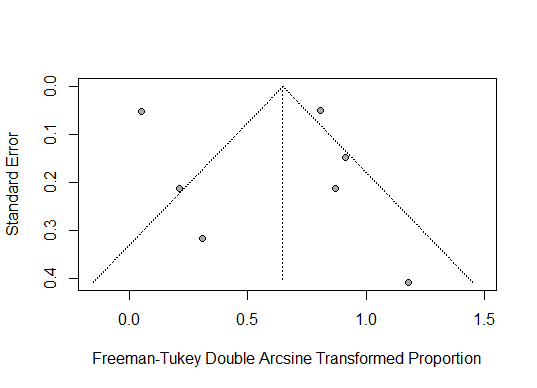
**A.**

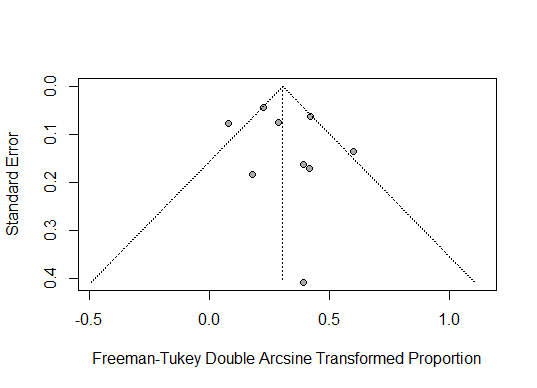
**B.**

Funnel plots presented for A) *Alouatta* genus and B) *Cebus/Sapajus* genus

**S10 Table. Complete arthropod results by genus**

| **Study** | **Test Method** | **Genus** | **Total (n)** | **MAYV Detected?** |
| --- | --- | --- | --- | --- |
| Aitken, 1969 [36] | Virus isolation | *Acanthocera* | 1 | No |
|  |  | *Aedomyia* | 12 | No |
|  |  | *Aedes* | 367994 | No |
|  |  | *Amblyomma* | 6550 | No |
|  |  | *Anopheles* | 66073 | No |
|  |  | *Boophilus* | 1759 | No |
|  |  | *Chrysops* | 87 | No |
|  |  | *Culex* | 238771 | No |
|  |  | *Culicoides* | 10079 | No |
|  |  | *Deinocerites* | 15 | No |
|  |  | *Diachlorus* | 1078 | No |
|  |  | *Fahrenholzia* | 187 | No |
|  |  | *Gigantolaelaps* | 5312 | No |
|  |  | *Haemagogus* | 18561 | No |
|  |  | *Hoplopleura* | 2080 | No |
|  |  | *Ixodes* | 78 | No |
|  |  | *Leucotabanus* | 18 | No |
|  |  | *Limatus* | 50730 | No |
|  |  | *Mansonia* | 127963 | Yes |
|  |  | *Ornithodoros* | 1922 | No |
|  |  | *Orthopodomyia* | 19 | No |
|  |  | *Philornis* | 179 | No |
|  |  | *Phlebotomus* | 12841 | No |
|  |  | *Phoniomyia* | 63689 | No |
|  |  | *Polygenis* | 386 | No |
|  |  | *Psorophora* | 177136 | No |
|  |  | *Rhipicephalus* | 270 | No |
|  |  | *Sabethes* | 5059 | No |
|  |  | *Simulium* | 546 | No |
|  |  | *Stibasoma* | 35 | No |
|  |  | *Stomoxys* | 23 | No |
|  |  | *Xenopsylla* | 7 | No |
|  |  | *Tabanus* | 72 | No |
|  |  | *Trichoprosopon* | 20500 | No |
|  |  | *Uranotaenia* | 71 | No |
|  |  | *Wyeomyia* | 346093 | No |
| Azevedo, 2009 [37] | Virus isolation | *Haemagogus* | 188 | Yes |
|  |  | Other^a^ | 644 | No |
| Batista, 2012 [5] | Virus isolation | *Aedes* | 59 | No |
|  |  | *Culex* | 8 | No |
|  |  | *Haemagogus* | 11 | No |
|  |  | *Psorophora* | 9 | No |
|  |  | *Sabethes* | 35 | No |
| Carrera, 2020 [38] | RT-PCR | *Culex* | 113 | No |
| Catenacci, 2017 [11] | RT-PCR | *Aedes* | 1 | No |
|  |  | *Culex* | 3 | No |
|  |  | *Flebotominae* | 11 | No |
|  |  | *Haemagogus* | 17 | No |
|  |  | *Limatus* | 83 | No |
|  |  | *Mansonia* | 1 | No |
|  |  | *Psorophora* | 1 | No |
|  |  | *Sabethes* | 6 | No |
|  |  | *Runchomya* | 1 | No |
|  |  | *Wyeomyia* | 115 | Yes |
| Degallier, 1992 [14] | Virus isolation | NA | 2005069 | No |
| Esposito, 2015 [39] | Virus isolation | *Haemagogus* | NA | Yes |
| Ferreira, 2020 [40] | RT-PCR; Virus isolation | *Aedes* | 1139 | No |
|  |  | *Culex* | 9429 | Yes |
|  |  | *Psorophora* | 1 | No |
| Galindo, 1966 [20] | Virus isolation | *Aedes* | 50616 | No |
|  |  | *Anopheles* | 4515 | No |
|  |  | *Culex* | 166921 | No |
|  |  | *Mansonia* | 59693 | No |
|  |  | *Phlebotomus* | 29651 | No |
|  |  | *Psorophora* | 28087 | Yes |
|  |  | *Sabethes* | 4332 | No |
|  |  | *Trichoprosopon* | 1901 | No |
| Galindo, 1967 [41] | Virus isolation | *Culex* | 11829 | Yes |
| Galindo, 1983 [2] | Virus isolation | *Aedes* | NA | No |
|  |  | *Aedomyia* | NA | No |
|  |  | *Anopheles* | NA | No |
|  |  | *Culex* | NA | No |
|  |  | *Culicoides* | NA | No |
|  |  | *Haemagogus* | NA | Yes |
|  |  | *Lutzomyia* | NA | No |
|  |  | *Mansonia* | NA | No |
|  |  | *Wyeomyia* | NA | No |
| GenBank KY618129 | Virus isolation | *Haemagogus* | NA | Yes |
| GenBank KY618130 | Virus isolation | *Culex* | NA | Yes |
| Groot, 1961 [42] | Virus isolation | *Aedes* | 14524 | No |
|  |  | *Anopheles* | 355 | No |
|  |  | *Culex* | 2420 | No |
|  |  | *Haemagogus* | 444 | No |
|  |  | *Limatus* | 236 | No |
|  |  | *Mansonia* | 2218 | No |
|  |  | *Psorophora* | 20111 | Yes |
|  |  | *Sabethes* | 275 | No |
|  |  | *Trichoprosopon* | 659 | No |
|  |  | *Wyeomyia* | 322 | No |
| Henriques, 2008 [43] | RT-PCR | *Aedes* | 1971 | No |
|  |  | *Aedomyia* | 23 | No |
|  |  | *Anopheles* | 8997 | No |
|  |  | *Cerathopogonidae* | 735 | No |
|  |  | *Coquilletidia* | 2710 | No |
|  |  | *Culex* | 7864 | No |
|  |  | *Culiseta* | 12 | No |
|  |  | *Deinocerites* | 208 | No |
|  |  | *Haemagogus* | 226 | No |
|  |  | *Limatus* | 67 | No |
|  |  | *Mansonia* | 6469 | No |
|  |  | *Psychodidae* | 4820 | No |
|  |  | *Psorophora* | 2233 | No |
|  |  | *Sabethes* | 18 | No |
|  |  | *Shannoniana* | 27 | No |
|  |  | *Simuliidae* | 828 | No |
|  |  | *Tabanidae* | 139 | No |
|  |  | *Trichoposopon* | 103 | No |
|  |  | *Uranotaenia* | 15 | No |
|  |  | *Wyeomyia* | 54 | No |
| Hoch, 1981 [4] | Virus isolation | *Aedes* | 11 | No |
|  |  | *Culex* | 424 | No |
|  |  | *Culicoides* | 3609 | No |
|  |  | *Foricypomyia* | 88 | No |
|  |  | *Haemagogus* | 2284 | Yes |
|  |  | *Limatus* | 2192 | No |
|  |  | *Lutzomyia* | 574 | No |
|  |  | *Orthopodomyia* | 69 | No |
|  |  | *Psorophora* | 77 | No |
|  |  | *Sabethes* | 465 | No |
|  |  | *Trichoprosopon* | 244 | No |
|  |  | *Wyeomyia* | 630 | No |
| Kubiszeski, 2017 [44] | RT-PCR; Virus isolation | *Aedomyia* | 90 | No |
|  |  | *Aedes* | 7 | No |
|  |  | *Culex* | 634 | Yes |
|  |  | *Coquilletidia* | 2 | No |
|  |  | *Haemagogus* | 1 | No |
|  |  | *Ochelorotatus* | 26 | No |
|  |  | *Psorophora* | 2 | No |
|  |  | *Uranotaenia* | 16 | No |
| Maia, 2019 [45] | RT-PCR; Virus isolation | *Aedes* | 4786 | Yes |
| Martinez, 2020 [46] | RT-PCR | *Aedes* | 169 | No |
| Pauvolid-Correa, 2008 [47] |  | *Aedomyia* | 1 | No |
|  |  | *Amblyomma* | 30 | No |
|  |  | *Anocentor* | 40 | No |
|  |  | *Anopheles* | 350 | No |
|  |  | *Coquillettidia* | 35 | No |
|  |  | *Culex* | 156 | No |
|  |  | *Mansonia* | 1799 | No |
|  |  | *Ochlerotatus* | 14 | No |
|  |  | *Psorophora* | 1325 | No |
|  |  | *Sabethes* | 4 | No |
| Pinheiro, 1974 [16] | Virus isolation | *Aedes* | NA | No |
|  |  | *Anopheles* | NA | No |
|  |  | *Culex* | NA | No |
|  |  | *Psorophora* | NA | No |
|  |  | *Trichoprosopon* | NA | No |
|  |  | *Uranotaenia* | NA | No |
|  |  | *Wyeomyia* | NA | No |
| Pinheiro, 2019 [48] | RT-PCR | *Aedes* | 125 | No |
|  |  | *Aedomyia* | 5 | No |
|  |  | *Anopheles* | 1 | No |
|  |  | *Culex* | 46 | No |
|  |  | *Haemagogus* | 200 | No |
|  |  | *Limatus* | 166 | No |
|  |  | *Psorophora* | 273 | No |
|  |  | *Runchomyia* | 3 | No |
|  |  | *Sabethes* | 42 | No |
|  |  | *Wyeomyia* | 6 | No |
| Powers, 2006 [49] | Virus isolation | *Haemagogus* | NA | Yes |
|  |  | *Ixodes* | NA | Yes |
|  |  | *Mansonia* | NA | Yes |
| SanMartin, 1973 [19] | Virus isolation | *Aedes* | 309 | No |
|  |  | *Aedomyia* | 9 | No |
|  |  | *Anopheles* | 302 | No |
|  |  | *Culex* | 13590 | No |
|  |  | *Limatus* | 1 | No |
|  |  | *Mansonia* | 13119 | No |
|  |  | *Psorophora* | 83 | No |
|  |  | *Uranotaenia* | 3 | No |
|  |  | *Wyeomyia* | 21 | No |
| Scherer, 1975 [50] | Virus isolation | NA | 18500 | No |
| Serra, 2016 [51] | RT-PCR; Virus isolation | *Aedes* | 1089 | Yes |
|  |  | *Culex* | 3433 | Yes |
|  |  | *Galindomyia* | 1 | No |
|  |  | *Limatus* | 7 | No |
|  |  | *Mansonia* | 4 | No |
|  |  | *Psorophora* | 21 | No |
|  |  | *Sabethes* | 1 | No |
|  |  | *Uranotaenia* | 1 | No |
| Silva, 2017 [52] | RT-PCR | *Aedes* | 77 | No |
|  |  | *Aedomyia* | 3 | No |
|  |  | *Anopheles* | 209 | No |
|  |  | *Chagasia* | 7 | No |
|  |  | *Coquillettidia* | 125 | No |
|  |  | *Culex* | 862 | No |
|  |  | *Haemagogus* | 78 | No |
|  |  | *Jhonbelkinia* | 73 | No |
|  |  | *Limatus* | 81 | No |
|  |  | *Mansonia* | 3 | No |
|  |  | *Ochlerotatus* | 695 | No |
|  |  | *Psorophora* | 1230 | No |
|  |  | *Runchomyia* | 1 | No |
|  |  | *Sabethes* | 83 | No |
|  |  | *Trichoprosopon* | 12 | No |
|  |  | *Uranotaenia* | 25 | No |
|  |  | *Wyeomyia* | 186 | No |
| Tauro, 2019 [53] | RT-PCR; Virus isolation | *Aedes* | 26 | No |
|  |  | *Culex* | 99 | No |
| Taylor, 1967 [15] | Virus isolation | *Culex* | NA | Yes |
|  |  | *Gigantolaelaps* | NA | Yes |
|  |  | *Haemagogus* | NA | Yes |
|  |  | *Mansonia* | NA | Yes |
|  |  | *Sabethes* | NA | Yes |

^a^ Includes *Wyeomyia*, *Aedes*, *Sabethes*, and *Limatus*

**S11 Table. Egger’s test for publication bias**

|  | **Studies (n)^a^** | **Pooled Prev.** | **95% CI** | **Egger Test p-value** |
| --- | --- | --- | --- | --- |
| **Primate order** | 13 | 13.1 | 4.3; 25.1 | **0.8945** |
| ***Cebus/Sapajus* genus** | 9 | 7.5 | 3.5; 15.3 | **0.4024** |
| ***Alouatta* genus** | 8 | 24.0 | 2.2; 81.6 | **0.6077** |
| **Pilosa order** | 7 | 0.0 | 0.0; 6.6 | **0.6759** |
| **Rodentia order** | 7 | 1.3 | 0.0; 6.5 | **0.6299** |
| **Domestic Equids** | 6 | 1.1 | 0.0; 4.5 | **0.3543** |
| **Didelphimorphia order** | 6 | 2.0 | 0.0; 7.2 | **0.1446** |
| **Carnivora order** | 5 | 0.1 | 0.0; 8.1 | **0.8822** |

^a^ Egger’s test was only conducted for meta-analyses that include five or more studies. Egger’s test should be interpreted with caution if 10 studies or less are included in the analysis [54].

References

1.	Seymour C, Peralta PH, Montgomery GG. Serologic evidence of natural togavirus infections in Panamanian sloths and other vertebrates. Am J Trop Med Hyg. 1983;32(4):854-61. Epub 1983/07/01. doi: 10.4269/ajtmh.1983.32.854. PubMed PMID: 6309027.

2.	Galindo P, Adames A, Peralta P, Johnson C, Read R. Impacto de la hidroeléctrica de Bayano en la transmisión de arbovirus. Rev Med Pan. 1983;8:89-134.

3.	Groot H. Estudios sobre virus transmitidos por artropodos en Colombia. Rev Acad Colomb Cienc. 1964;12(46):191-217. doi: 10.18257/raccefyn.565.

4.	Hoch AL, Peterson NE, LeDuc JW, Pinheiro FP. An outbreak of Mayaro virus disease in Belterra, Brazil. III. Entomological and ecological studies. Am J Trop Med Hyg. 1981;30(3):689-98. Epub 1981/05/01. doi: 10.4269/ajtmh.1981.30.689. PubMed PMID: 6266265.

5.	Batista PM, Andreotti R, Chiang JO, Ferreira MS, Vasconcelos PF. Seroepidemiological monitoring in sentinel animals and vectors as part of arbovirus surveillance in the state of Mato Grosso do Sul, Brazil. Rev Soc Bras Med Trop. 2012;45(2):168-73. Epub 2012/04/27. doi: 10.1590/s0037-86822012000200006. PubMed PMID: 22534986.

6.	Gibrail MM. Detecção de anticorpos para arbovirus em primatas não humanos no município de Goiânia, Goiás [M.Sc. Thesis]. Goiânia: Universidade Federal de Goiás; 2015. Available from: <https://repositorio.bc.ufg.br/tede/handle/tede/5552>.

7.	Diaz LA, Diaz Mdel P, Almiron WR, Contigiani MS. Infection by UNA virus (Alphavirus; Togaviridae) and risk factor analysis in black howler monkeys (Alouatta caraya) from Paraguay and Argentina. Trans R Soc Trop Med Hyg. 2007;101(10):1039-41. Epub 2007/07/31. doi: 10.1016/j.trstmh.2007.04.009. PubMed PMID: 17658571.

8.	Perez JG, Carrera JP, Serrano E, Pitti Y, Maguina JL, Mentaberre G, et al. Serologic Evidence of Zoonotic Alphaviruses in Humans from an Indigenous Community in the Peruvian Amazon. Am J Trop Med Hyg. 2019. Epub 2019/10/02. doi: 10.4269/ajtmh.18-0850. PubMed PMID: 31571566.

9.	de Thoisy B, Gardon J, Salas RA, Morvan J, Kazanji M. Mayaro virus in wild mammals, French Guiana. Emerg Infect Dis. 2003;9(10):1326-9. Epub 2003/11/12. doi: 10.3201/eid0910.030161. PubMed PMID: 14609474; PubMed Central PMCID: PMCPMC3033094.

10.	Moreira-Soto A, Carneiro ID, Fischer C, Feldmann M, Kummerer BM, Silva NS, et al. Limited Evidence for Infection of Urban and Peri-urban Nonhuman Primates with Zika and Chikungunya Viruses in Brazil. mSphere. 2018;3(1). doi: 10.1128/mSphere.00523-17. PubMed PMID: WOS:000425277500024.

11.	Catenacci LS. Abordagem one health para vigilância de arbovirus na Mata Atlântica do sul da Bahia, Brasil. [Ph.D. Thesis]. Ananindeua: Instituto Evandro Chagas; 2017. Available from: <https://patua.iec.gov.br/handle/iec/3073>.

12.	Laroque PO, Valença-Montenegro MM, Ferreira DRA, Chiang JO, Cordeiro MT, Vasconcelos PFC, et al. Levantamento soroepidemiológico para arbovírus em macaco-prego-galego (Cebus flavius) de vida livre no estado da Paraíba e em macaco-prego (Cebus libidinosus) de cativeiro do nordeste do Brasil. Pesq Vet Bras. 2014;34:462-8.

13.	Paulo M, Renato A, Da Carneiro Rocha T, Eliane C, Navarro da Silva M. Serosurvey of arbovirus in free-living non-human primates (Sapajus spp.) in Brazil. J Environ Anal Chem. 2015;2(155):2380-91.1000155.

14.	Degallier N, Travassos da Rosa AP, Vasconcelos PFC, Hervé JP, Sa Filho GC, Travassos da Rosa JFS, et al. Modifications of arbovirus transmission in relation to construction of dams in Brazilian Amazonia Journal of the Brazilian Association for the Advancement of Science. 1992;44.

15.	Taylor RM. Catalogue of arthropod-borne viruses of the world: a collection of data on registered arthropod-borne animal viruses: US Public Health Service; 1967.

16.	Pinheiro FP, Bensabath G, Andrade AH, Lins ZC, Fraihi H, Tang AT, et al. Infectious diseases along Brazil's Trans-Amazon Highway: surveillance and research. Bull Pan Am Health Organ. 1974;8(111).

17.	Srihongse S, Galindo P, Eldridge BF. A survey to assess potential human disease hazards along proposed sea level canal routes in Panama and Colombia. V. Arbovirus infection in non human vertebrates. Mil Med. 1974;139(6):449-53.

18.	Nunes MR, Barbosa TF, Casseb LM, Nunes Neto JP, Segura Nde O, Monteiro HA, et al. Eco-epidemiologia dos arbovirus na area de influencia da rodovia Cuiaba-Santarem (BR 163), Estado do Para, Brasil. Cad Saude Publica. 2009;25(12):2583-602. Epub 2010/03/02. doi: 10.1590/s0102-311x2009001200006. PubMed PMID: 20191150.

19.	Sanmartín C, Mackenzie RB, Trapido H, Barreto P, Mullenax CH, Gutiérrez E, et al. Encefalitis equina venezolana en Colombia, 1967. Bol Oficina Sanit Panam. 1973;74(2):108-37. Epub 1973/02/01. PubMed PMID: 4265714.

20.	Galindo P, Srihongse S, De Rodaniche E, Grayson MA. An ecological survey for arboviruses in Almirante, Panama, 1959-1962. Am J Trop Med Hyg. 1966;15(3):385-400. Epub 1966/05/01. doi: 10.4269/ajtmh.1966.15.385. PubMed PMID: 4380043.

21.	Cruz ACR, Prazeres AdSCd, Gama EC, Lima MFd, Azevedo RdSS, Casseb LMN, et al. Vigilância sorológica para arbovírus em Juruti, Pará, Brasil. Cadernos de saude publica. 2009;25(11):2517-23.

22.	Price JL. Serological evidence of infection of Tacaribe virus and arboviruses in Trinidadian bats. Am J Trop Med Hyg. 1978;27(1 Pt 1):162-7. Epub 1978/01/01. doi: 10.4269/ajtmh.1978.27.162. PubMed PMID: 204207.

23.	Pauvolid-Correa A, Juliano RS, Campos Z, Velez J, Nogueira RM, Komar N. Neutralising antibodies for Mayaro virus in Pantanal, Brazil. Mem Inst Oswaldo Cruz. 2015;110(1):125-33. Epub 2015/03/06. doi: 10.1590/0074-02760140383. PubMed PMID: 25742272; PubMed Central PMCID: PMCPMC4371226.

24.	Medlin S, Deardorff ER, Hanley CS, Vergneau-Grosset C, Siudak-Campfield A, Dallwig R, et al. Serosurvey of Selected Arboviral Pathogens in Free-Ranging, Two-Toed Sloths (Choloepus Hoffmanni) and Three-Toed Sloths (Bradypus Variegatus) In Costa Rica, 2005-07. J Wildl Dis. 2016;52(4):883-92. Epub 2016/08/02. doi: 10.7589/2015-02-040. PubMed PMID: 27479900; PubMed Central PMCID: PMCPMC5189659.

25.	Calisher CH, Gutierrez E, Maness KS, Lord RD. Isolation of Mayaro virus from a migrating bird captured in Louisiana in 1967. Bull Pan Am Health Organ. 1974;8(3):243-8. Epub 1974/01/01. PubMed PMID: 4418030.

26.	Araújo FAA, Vianna RdST, Andrade Filho GVd, Melhado DL, Todeschini B, Cavalcante e Cavalcanti G, et al. Segundo inquérito sorológico em aves migratórias e residentes do parque nacional da Lagoa do Peixe/RS para detecção do vírus da Febre da Febre do Nilo Ocidental e outros vírus. In: Ministério da Saúde Secretaria de Vigilância em Saúde, editor. Boletim Eletrônico Epidemiologico, 2004.

27.	Araujo FAA, Lima PC, Andrade MA, de Sá Jayme V, Ramos DG, Da Silveira SL. Soroprevalência de anticorpos “anti-arbovírus” de importância em saúde pública em aves selvagens, Brasil–2007 e 2008. Ciênc Anim Brasil. 2012;13(1):115-23. doi: 10.5216/cab.v13i1.16834.

28.	Araujo FAA, Wada MY, da Silva EV, Cavalcante GC, Magalhaes VS, de Andrade Filho GV, et al. Primeiro inquérito sorológico em aves migratórias e nativas do Parque Nacional da Lagoa do Peixe/RS para detecção do vírus do Nilo Ocidental. In: Ministério da Saúde Secretaria de Vigilância em Saúde, editor. Boletim Eletrônico Epidemiologico, 2003.

29.	Gomes FA, Jansen AM, Machado RZ, Jesus Pena HF, Fumagalli MJ, Silva A, et al. Serological evidence of arboviruses and coccidia infecting horses in the Amazonian region of Brazil. PloS One. 2019;14(12):e0225895. Epub 2019/12/13. doi: 10.1371/journal.pone.0225895. PubMed PMID: 31830142.

30.	Pauvolid-Correa A, Tavares FN, Costa EV, Burlandy FM, Murta M, Pellegrin AO, et al. Serologic evidence of the recent circulation of Saint Louis encephalitis virus and high prevalence of equine encephalitis viruses in horses in the Nhecolandia sub-region in South Pantanal, Central-West Brazil. Mem Inst Oswaldo Cruz. 2010;105(6):829-33. Epub 2010/10/15. doi: 10.1590/s0074-02762010000600017. PubMed PMID: 20945001.

31.	Araujo FAA, Andrade MA, Jayme VS, Santos AL, Roman APM, Ramos DG, et al. Anticorpos antialfavírus detectados em equinos durante diferentes epizootias de encefalite equina, Paraíba, 2009. Rev Bras Ciênc Vet. 2012;19(1):80-5. doi: 10.4322/rbcv.2014.086.

32.	Casseb AdR, Brito TC, Silva MRMd, Chiang JO, Martins LC, Silva SPd, et al. Prevalence of antibodies to equine alphaviruses in the State of Pará, Brazil. Arq Inst Biol. 2016;83. doi: 10.1590/1808-1657000202014.

33.	Casseb AdR. Soroprevalência de anticorpos e padronização do teste ELISA sanduíche indireto para 19 tipos de arbovírus em herbívoros domésticos [Ph.D. Thesis]. Belém: Universidade Federal do Pará; 2010. Available from: <http://repositorio.ufpa.br/jspui/handle/2011/4760>.

34.	Medina G, Garzaro DJ, Barrios M, Auguste AJ, Weaver SC, Pujol FH. Genetic diversity of Venezuelan alphaviruses and circulation of a Venezuelan equine encephalitis virus subtype IAB strain during an interepizootic period. Am J Trop Med Hyg. 2015;93(1):7-10. Epub 2015/05/06. doi: 10.4269/ajtmh.14-0543. PubMed PMID: 25940191; PubMed Central PMCID: PMCPMC4497907.

35.	Turell MJ, Gozalo AS, Guevara C, Schoeler GB, Carbajal F, Lopez-Sifuentes VM, et al. Lack of Evidence of Sylvatic Transmission of Dengue Viruses in the Amazon Rainforest Near Iquitos, Peru. Vector Borne Zoonotic Dis. 2019;19(9):685-9. Epub 2019/04/10. doi: 10.1089/vbz.2018.2408. PubMed PMID: 30964397; PubMed Central PMCID: PMCPMC6716187.

36.	Aitken TH, Spence L, Jonkers AH, Downs WG. A 10-year survey of Trinidadian arthropods for natural virus infections (1953-1963). J Med Entomol. 1969;6(2):207-15. Epub 1969/05/01. doi: 10.1093/jmedent/6.2.207. PubMed PMID: 5807863.

37.	Azevedo RS, Silva EV, Carvalho VL, Rodrigues SG, Neto JPN, Monteiro HA, et al. Mayaro fever virus, Brazilian amazon. Emerg Infect Dis. 2009;15(11):1830. doi: 10.3201/eid1511.090461.

38.	Carrera JP, Cucunubá ZM, Neira K, Lambert B, Pittí Y, Liscano J, et al. Endemic and Epidemic Human Alphavirus Infections in Eastern Panama: An Analysis of Population-Based Cross-Sectional Surveys. Am J Trop Med Hyg. 2020. Epub 2020/10/31. doi: 10.4269/ajtmh.20-0408. PubMed PMID: 33124532.

39.	Esposito DL, da Fonseca BA. Complete Genome Sequence of Mayaro Virus (Togaviridae, Alphavirus) Strain BeAr 20290 from Brazil. Genome Announc. 2015;3(6). Epub 2015/12/19. doi: 10.1128/genomeA.01372-15. PubMed PMID: 26679574; PubMed Central PMCID: PMCPMC4683219.

40.	da Silva Ferreira R, de Toni Aquino da Cruz LC, Souza VJ, da Silva Neves NA, de Souza VC, Filho LCF, et al. Insect-specific viruses and arboviruses in adult male culicids from Midwestern Brazil. Infect Genet Evol. 2020:104561. Epub 2020/09/23. doi: 10.1016/j.meegid.2020.104561. PubMed PMID: 32961364.

41.	Galindo P, Srihongse S. Transmission of arboviruses to hamsters by the bite of naturally infected Culex (Melanoconion) mosquitoes. Am J Trop Med Hyg. 1967;16(4):525-30. Epub 1967/07/01. doi: 10.4269/ajtmh.1967.16.525. PubMed PMID: 4952151.

42.	Groot H, Morales A, Vidales H. Virus isolations from forest mosquitoes in San Vicente de Chucuri, Colombia. Am J Trop Med Hyg. 1961;10:397-402. Epub 1961/05/01. doi: 10.4269/ajtmh.1961.10.397. PubMed PMID: 13708940.

43.	Henriques DA. Caracterização molecular de arbovírus isolados da fauna diptera nematocera do Estado de Rondônia (Amazônia ocidental brasileira) [Ph.D. Thesis]. São Paulo: Universidade de São Paulo; 2008. Available from: <https://teses.usp.br/teses/disponiveis/42/42132/tde-27032009-124003/pt-br.php>.

44.	Kubiszeski JR. Arboviroses emergentes no município de Sinop-MT: pesquisa de vetores [Ph.D. Thesis]. Sinop: Universidade Federal de Mato Grosso; 2016. Available from: <https://teses.usp.br/teses/disponiveis/42/42132/tde-27032009-124003/pt-br.php>.

45.	Maia LMS, Bezerra MCF, Costa MCS, Souza EM, Oliveira MEB, Ribeiro ALM, et al. Natural vertical infection by dengue virus serotype 4, Zika virus and Mayaro virus in Aedes (Stegomyia) aegypti and Aedes (Stegomyia) albopictus. Med Vet Entomol. 2019;33(3):437-42. Epub 2019/02/19. doi: 10.1111/mve.12369. PubMed PMID: 30776139.

46.	Martinez D, Hernandez C, Munoz M, Armesto Y, Cuervo A, Ramirez JD. Identification of Aedes (Diptera: Culicidae) Species and Arboviruses Circulating in Arauca, Eastern Colombia. Front Ecol Evol. 2020;8. doi: 10.3389/fevo.2020.602190. PubMed PMID: WOS:000596835300001.

47.	Pauvolid-Correa A. Estudo sobre arbovírus em populações de eqüinos e artrópodes na sub-região da Nhecolândia no Pantanal de Mato Grosso do Sul [M.Sc. Thesis]. Rio de Janeiro: Fundação Oswaldo Cruz; 2008. Available from: <https://www.arca.fiocruz.br/handle/icict/21142>.

48.	Pinheiro GG, Rocha MN, de Oliveira MA, Moreira LA, Andrade JD. Detection of Yellow Fever Virus in Sylvatic Mosquitoes during Disease Outbreaks of 2017-2018 in Minas Gerais State, Brazil. Insects. 2019;10(5). doi: 10.3390/insects10050136. PubMed PMID: WOS:000476846800018.

49.	Powers AM, Aguilar PV, Chandler LJ, Brault AC, Meakins TA, Watts D, et al. Genetic relationships among Mayaro and Una viruses suggest distinct patterns of transmission. Am J Trop Med Hyg. 2006;75(3):461-9. Epub 2006/09/14. PubMed PMID: 16968922.

50.	Scherer WF, Madalengoitia J, Flores W, Acosta M. The first isolations of eastern encephalitis, group C, and Guama group arboviruses from the Peruvian Amazon region of western South America. Bull Pan Am Health Organ. 1975;9(1):19-26. Epub 1975/01/01. PubMed PMID: 238693.

51.	Serra OP, Cardoso BF, Ribeiro AL, Santos FA, Slhessarenko RD. Mayaro virus and dengue virus 1 and 4 natural infection in culicids from Cuiaba, state of Mato Grosso, Brazil. Mem Inst Oswaldo Cruz. 2016;111(1):20-9. Epub 2016/01/20. doi: 10.1590/0074-02760150270. PubMed PMID: 26784852; PubMed Central PMCID: PMCPMC4727432.

52.	Silva JWP. Aspectos ecológicos de vetores putativos do Vírus Mayaro e Vírus Oropuche em estratificação vertical e horizontal em ambientes florestais e antropizados em uma comunidade rural no Amazonas [M.Sc. Thesis]. Manaus, AM: Oswaldo Cruz Foundation, Instituto Leônidas and Maria Deane; 2017. Available from: <https://www.arca.fiocruz.br/handle/icict/23337>.

53.	Tauro LB, Cardoso CW, Souza RL, Nascimento LC, Santos DRD, Campos GS, et al. A localized outbreak of Chikungunya virus in Salvador, Bahia, Brazil. Mem Inst Oswaldo Cruz. 2019;114:e180597. Epub 2019/03/08. doi: 10.1590/0074-02760180597. PubMed PMID: 30843962; PubMed Central PMCID: PMCPMC6396974.

54.	Egger M, Davey Smith G, Schneider M, Minder C. Bias in meta-analysis detected by a simple, graphical test. BMJ. 1997;315(7109):629-34. Epub 1997/10/06. doi: 10.1136/bmj.315.7109.629. PubMed PMID: 9310563; PubMed Central PMCID: PMCPMC2127453.
